# Ohm’s Law for increasing fitness gene expression with selection pressure

**DOI:** 10.1101/693234

**Authors:** Marta Ciechonska, Marc Sturrock, Alice Grob, Gerald Larrouy-Maumus, Vahid Shahrezaei, Mark Isalan

**Author notes:** To whom correspondence should be addressed: Mark Isalan and Vahid Shahrezaei, Imperial College London, London SW7 2AZ, UK. Contributed equally (Marta Ciechonska, Marc Sturrock).

## Abstract

Natural selection relies on genotypic and phenotypic adaptation in response to fluctuating environmental conditions and is the key to predicting and preventing drug resistance. Whereas classic persistence is all-or-nothing, here we show for the first time that an antibiotic resistance gene displays linear dose-responsive selection for increased expression in proportion to rising antibiotic concentration in *E. coli*. Furthermore, we observe the general nature of an instantaneous phenotypic selection process upon bactericidal and bacteriostatic antibiotic treatment, as well as an amino acid synthesis pathway enzyme under a range of auxotrophic conditions. To explain this phenomenon, we propose an analogy to Ohm’s law in electricity (V=IR) where fitness pressure acts similarly to voltage (V), gene expression to current (I), and resistance (R) to cellular machinery constraints. Lastly, mathematical modelling approaches reveal that the emergent gene expression mechanism requires variation in mRNA and protein production within an isogenic population, and cell ‘memory’ from positive feedbacks between growth and expression of any fitness-inducing gene.

---

Fitness-induced phenotypic variability occurs when individual cells within a genetically identical population spontaneously shift to a fitter state of gene expression. This eliminates both the need for selection based on genetic mutation and the high cost of evolving and maintaining a specialised sensory apparatus for every eventuality^1–3^. Such transient phenotypic selection, rooted in heterogenous expression of a fitness gene during changing environmental conditions, can result in selection according to need at the population level^4–9^. Due to its non-permanent nature, this behaviour is difficult to isolate and characterise, however, it appears to be a common survival strategy, as suggested in reports of several unrelated phenomena. These include: phenotypic selection for transient upregulation of a kanamycin resistance gene in *Salmonella*^10^, and the multiple resistance activator MarA in *E. coli* in the presence of antibiotics^11^; increased expression of an essential histidine biosynthesis gene in *E.coli* grown in the absence of this amino acid^12^; and cancer cell population survival in the presence of high concentrations of puromycin, based on a positive feedback-dependent heterogeneity of fitness gene expression^13^. These observations point to a general reversible phenotypic selection mechanism, the detailed characterization of which may aid in the prediction of population-level adaptation dynamics of antimicrobial and cancer chemotherapy resistance development. Here, we apply synthetic biology approaches and generate quantitative data to guide mathematical modelling in search of conditions underpinning this emergent gene expression (EGE) behaviour. We report a novel dose-dependent relationship of fitness gene expression in response to increasing fitness pressure, which is transient and is based exclusively on phenotypic selection. We thus propose a new concept in understanding cell population-level antibiotic resistance, which is distinct from - and complements - well-documented survival strategies such as persistence and heteroresistance^14,15^.

To isolate EGE, we applied a fitness pressure, using antibiotic challenge with the protein synthesis inhibitor chloramphenicol (Cm), and analysed phenotypic adaptation output based on changing expression levels of chloramphenicol acetyl transferase (CAT). We integrated wild type (WT) and mutant versions of *gfp-cat* fusion constructs, driven by an arabinose-inducible promoter, into the *E. coli* genome (Fig. 1a). We expected that as Cm concentration increased, cells expressing higher levels of GFP-CAT would be more fit; consequently, *cat* expression and GFP fluorescence within the entire population would rise in a dose-dependent manner (Fig. 1b). Additionally, we reasoned that strains expressing less-active mutant versions of CAT would require higher expression levels than WT, in order to acetylate and neutralize equal amounts of antibiotic.

**Fig. 1:**
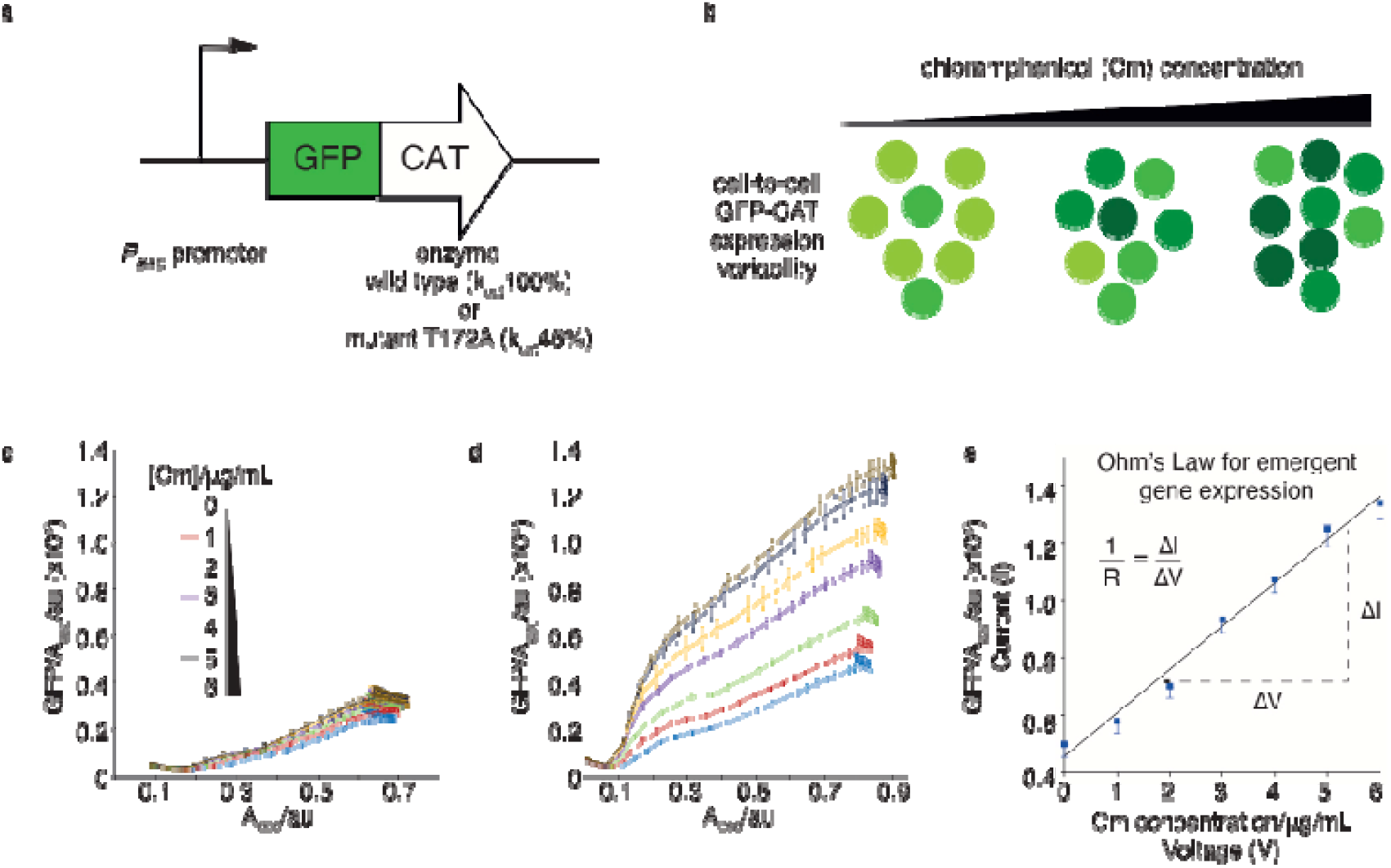
Emergent gene expression (EGE) increases linearly with rising antibiotic concentration in analogy to Ohm’s law. (a) Schematic of GFP-chloramphenicol acetyl transferase (CAT) expression cassettes used in this study with inducible (*P_BAD_*) promoter driving expression of wild type (wt) or mutant *cat_T172A_* (relative acetylation efficiencies of k_cat_100% and k_cat_ 46% respectively). (b) Schematic of emergent gene expression for antibiotic resistance. GFP-CAT concentration per cell is indicated by a range of light to dark green; three separate populations are shown where higher *gfp-cat* expression is spontaneously selected in the presence of increasing chloramphenicol (Cm) concentration, to increase cell fitness. (c,d) Emergent *gfp-cat* expression with a weakly-induced *P_BAD_* promoter (0.005% arabinose). Graphs show GFP/A_600_ per well of populations expressing genome-integrated wt *gfp-cat* (c) and mutant *gfp-cat _T172A_* (d). These were treated with 0, 1, 2, 3, 4, 5, or 6 μg/mL Cm to induce EGE, monitored for 12 hrs and compared at corresponding Abs_600_; n=3 biological replicates. (e) Ohm’s law analogy for emergent gene expression, where GFP-CAT expression is analogous to current (I), Cm concentration to voltage (V), and the slope of the linear relationship between these, representing cellular propensity to increase EGE, is analogous to conductance (1/R). The data points are the maximal emergent gene expression values in (d).

We challenged populations of cells weakly induced to express either the WT *gfp-cat* cassette or the mutant (T172A, ~46% of k_cat_ WT activity^16^), with increasing concentrations of chloramphenicol, and collected growth and GFP fluorescence time-series measurements. As expected, this resulted in slight upregulation of WT *gfp-cat* expression (Fig. 1c), and in strong, non-transient, Cm-dependent increases in mutant GFP-CAT_T172A_ production, well above the amount induced by arabinose alone (Fig. 1d). Strikingly, in all cases the increase in fitness gene (*cat*) expression was linearly correlated to rising fitness pressure (Cm concentration) (Fig. 1e).

We find it helpful to understand this observation using an analogy taken from electrical conductivity. In Ohm’s law, current (I) is proportional to voltage (V, electrical pressure), with resistance (R) being the opposition to electron flow. Here, by analogy, the fitness pressure (Cm concentration, “voltage”) drives a proportional increase in *gfp-cat* expression (“current”) with the metabolic and resource cost of gene expression accounting for the “resistance”, while the slope of the graph (1/R, Fig. 1e) gives the “conductance” of the system, or the propensity of the cell to increase EGE per unit fitness pressure.

Next, we applied a series of controls to test the EGE hypothesis. These included addition of a constitutively expressed *cat* gene to the *gfp-cat*_T172A_-expressing strain, thus relieving the fitness pressure and obviating the need for EGE. (Supplementary Data Fig. 1). Furthermore, *gfp* expression did not increase upon Cm treatment in a strain encoding a functionally inactive *gfp-cat_H193A_* mutant, suggesting that EGE selects only those cells which contain increased amounts of a functional fitness-conferring protein, and that Cm does not directly activate the promoter (Supplementary Data Fig. 1). We also found this phenotypic selection effect to be reversible and reproducible over several short rounds of antibiotic challenge and washout, reflecting the inherent flexibility of the mechanism (Supplementary Data Fig. 2). In addition, RT-qPCR analysis showed that *gfp-cat* transcripts were specifically upregulated upon Cm treatment, while expression of housekeeping genes, sigma factors, and other antibiotic treatment response genes remained constant (Supplementary Data Fig. 3). This suggests that phenotypic selection based on Cm-induced EGE is specific to the *gfp-cat* gene, that the level of upregulation is reversible and that it is related to the per-molecule activity of the fitness-conferring enzyme.

To understand the mechanisms underlying EGE, we established minimal requirements necessary to recapitulate this behaviour *in silico*. We constructed agent-based models that included growth and division of cells exhibiting stochastic gene expression of a fitness-inducing gene^17^. Previously established global links from both transcription and translation to cell growth were incorporated^5,18^. We also assumed that the antibiotic Cm impedes cell growth and translation and that CAT acetylates the antibiotic Cm.

As transcription and translation levels may be regulated by cellular growth rate^17^, we performed a comparison of 10 models exploring a range of assumptions for gene expression and growth regulation. We used Approximate Bayesian Computation (ABC)^19^ in conjunction with gene expression and growth kinetics data from our microplate experiments to constrain the parameters and perform model selection for most accurate capture of EGE behaviour (Fig. 2 a, b and Supplementary Data Video 1). We found that growth-dependent dilution and a global positive feedback coupling translation rate to cell growth were essential (Supplementary Data Table 1); positive feedback likely extends the lifetime of protein fluctuations beyond the dilution time set by the cell cycle^20^. The candidate models also explicitly required both transcription and translation components to produce EGE, as removing the mRNA variable did not produce sufficient levels of noise for phenotypic selection to act upon (Supplementary Figs. 8 to 11).

Importantly, without further fitting, the model was able to predict a Cm-dependent unimodal increase in CAT expression across the entire distribution of the population (Fig. 2c), which we validated using flow cytometry (Fig. 2d). The model similarly predicted experimentally observed increasing *gfp-cat* mRNA levels in the presence of increasing Cm concentration (Fig. 2e, Supplementary Data Fig. 3), was able to capture the time-course of external antibiotic depletion as observed by mass spectrometry (Fig. 2f), and fully replicated the reversibility of EGE upon removal of Cm (Supplementary Figs. 21 and 22).

**Fig. 2:**
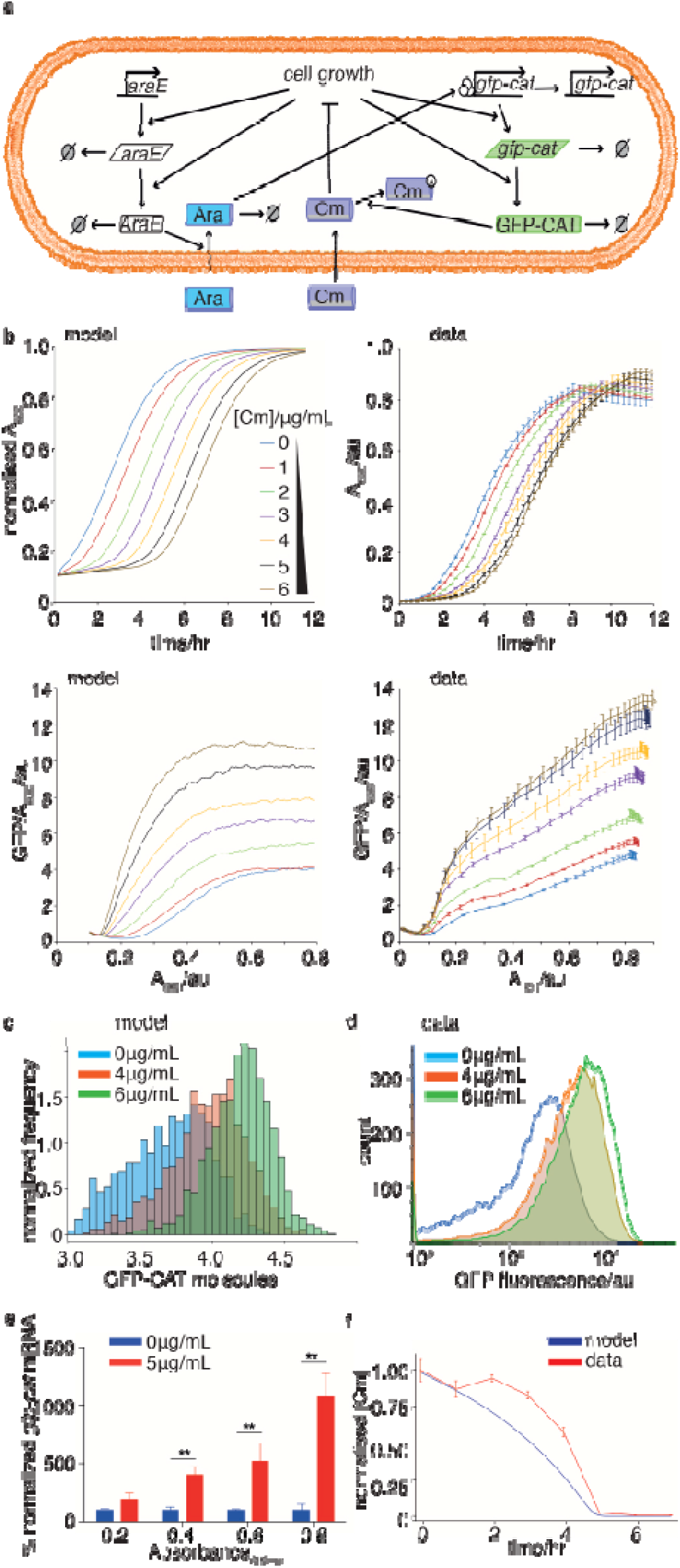
Experimental verification of the computational emergence model. (a) Schematic for mathematical modelling of the inducible promoter system for a single cell. The reactions capture *gfp-cat* expression and degradation, *araE* expression and degradation, arabinose import, degradation and activation of the *P_BAD_* promoter, as well as chloramphenicol (Cm) import and acetylation by GFP-CAT. Intracellular dynamics were coupled to a second model scale which captured cell division, partitioning and logistic growth of the cell population (more details in Supplementary Material Section 3 and Supplementary Figs. 16 to 20). (b) Comparing model outputs with experimental data for growth (normalised A_600_ data) and GFP-CAT molecules per cell (GFP/A_600_), for increasing Cm concentrations, over 12 hrs. Best fit parameters were taken from ABC parameter inference and initial conditions (initial number of cells and GFP expression levels) were taken from the experimental data displayed in right hand panels. mRNA levels were assumed to be zero initially. (c,d) Comparing model outputs with experimental data for population distributions of emergent gene expression. GFP-CAT molecule distributions are shown for increasing Cm concentrations. X-axis (c) is displayed on a log10 scale and y-axis is scaled such that the total area of the histograms sum to 1. The flow cytometry analysis (d) of the genome-integrated *gfp-cat_T172A_* mutant matches the model prediction (c). Cells in (d) were weakly induced with 0.005% arabinose, treated with 0, 4 or 6 μg/mL Cm for 12hrs, and fixed in 2% paraformaldehyde; n=3 biological replicates. (e) RT-qPCR assay of *gfp-cat_T172A_* mRNA expression in populations induced with 0.005% arabinose, treated with 0 (blue) or 5 (red) μg/mL Cm, and harvested at A_600_ of 0.2, 0.4, 0.6 and 0.8 (Mean ± SEm; n=3 biological replicates). Asterisks represent p values: ** = p<0.01. (f) Cm concentration in cultures of the genome-integrated *gfp-cat_T172A_* mutant strain, measured by LC-MS at one-hour intervals (orange). This is compared with the predicted Cm concentration from simulation of the refined inducible promoter model (Supplementary Fig. 23). Both time series are normalised to their maximal values.

We were thus able to build an accurate predictive model of EGE, explaining the associated fitness gene expression increases in the presence of a selection pressure. Based on these findings, we propose that population-level noise in gene expression ensures existence of cells with a range of fitness and higher expression of a fitness gene results in faster division times in the presence of a corresponding fitness pressure. Furthermore, positive feedback produces a memory effect in daughter cells, thus making phenotypic selection and EGE possible.

Fitness-induced gene expression effects in antibiotic resistance have been reported previously, however, due to the transiency of this phenomenon, collection of corresponding fine-grained experimental data for model fitting is difficult^21,22^. In this study, we used decreased-activity fitness gene mutants that enhance EGE to observe and quantify dose-responses. Similarly, using mathematical modelling, we revealed that to observe maximal EGE magnitude, one needs to use intermediate fitness gene strength (*cat_T172A_*) and intermediate pressure (Cm) (Fig. 3a). This interplay between fitness strength and pressure resulted in theoretically detectable bands or islands of maximal emergent gene expression within the fitness parameter landscape (Fig. 3a).

**Fig. 3:**
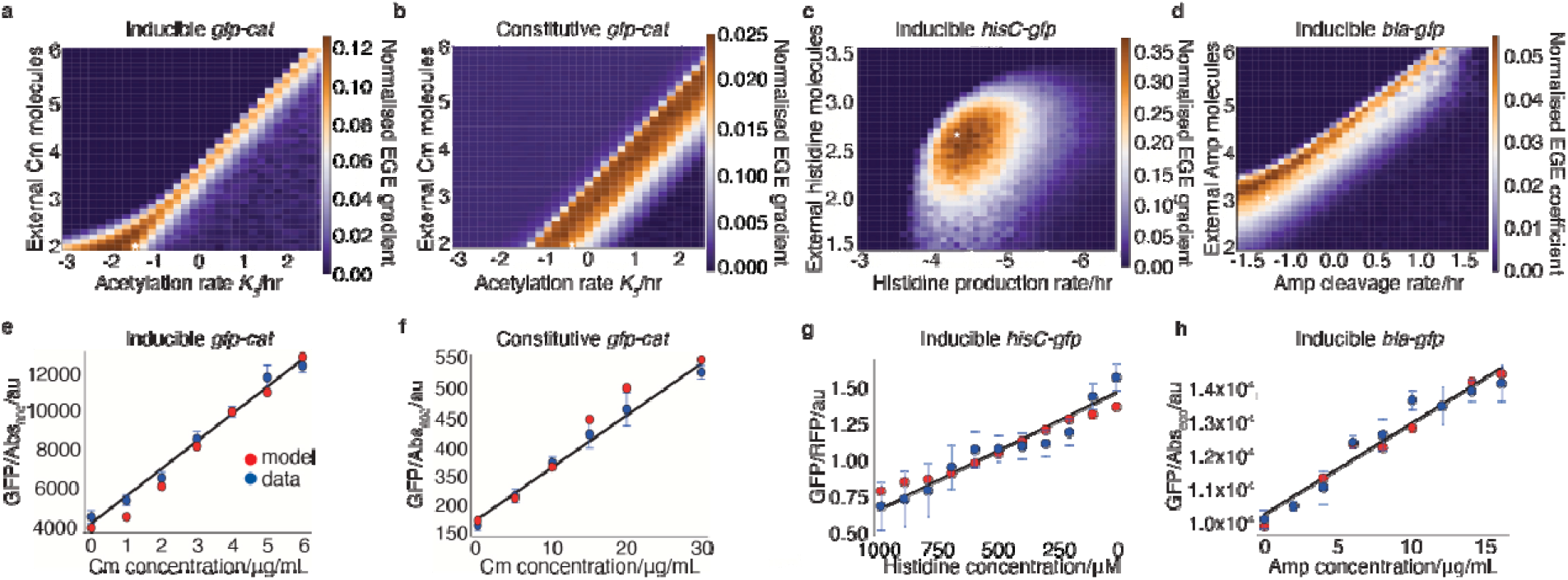
Bands and islands of maximal emergent gene expression, in antibiotic resistance and histidine auxotrophy, in parameter space. Heatmaps showing the effect of varying acetylation rate and chloramphenicol (Cm) dosage on a normalised emergent gene expression (EGE) gradient of *gfp-cat*, for (a) inducible (see Supplementary Figs. 16 to 19) and (b) constitutive promoter models (see Supplementary Figs. 12 to 15). Six different simulations were computed per model, corresponding to cells being treated with Cm ((a): 0C, 1C, 2C, 3C, 4C, 5C or 6C; (b): 0C, 5C, 10C, 15C, 20C or 30C; where C is the number of external Cm molecules). (c) Heatmap showing effect of varying histidinol-phosphate aminotransferase activity on a normalised emergent gene expression (EGE) gradient of *gfp-hisC*. Linear regression is applied to the HisC-GFP/RFP levels yielded from simulations of model of HisC dynamics (see Supplementary Fig. 24) generated using decreasing concentrations of histidine from 1000 to 0μM for each time point, to obtain the maximum gradient. (d) Heatmap showing the effect of varying the cleavage rate and concentration of ampicillin (Amp). Linear regression is applied to the *gfp-bla_L74N_* expression levels yielded from simulations of model of *blaL74N* dynamics (see Supplementary Fig. 25) generated using concentrations of Amp from 0, 2, 4, 6, 8, 10, 12, 14, and 16 μg/mL for each time point, to obtain the maximum gradient. White asterisks highlight the parameter combinations found to provide the best model fits to the microplate reader data. (e, f, g, h), Linear regression fits of microplate reader data showing linearity of GFP expression relationship (‘Ohm’s law’) with Cm treatment dosage, for both (e) inducible and (f) constitutive cases of *cat-gfp_T172A_* expression. (e) shows mean (± SDm) GFP expression (GFP/A_600_ per well) in populations treated with 0, 1, 2, 3, 4, 5 or 6 μg/mL Cm at t = 12 hrs (blue points) with linear regression fit overlaid (black solid line). These experimental data were compared with the predicted mean GFP-CAT molecules per cell for inducible promoter model simulations at same time point (red points). (f) is similar, for populations treated with 0, 5, 10, 15, 20, or 30 μg/mL Cm at t = 3 hrs (blue points), compared with model predictions (red points) of the constitutively expressed *gfp-cat_T172A_*. (g) Linear regression fit of *gfp* expression normalised to *rfp* expression in cells grown in minimal media with decreasing concentrations of histidine from 1000 to 0μM (red dots) compared with the predicted mean GFP-HisC/RFP molecule production at the respective concentrations. (h) Linear regression fit of inducible *gfp-bla_L74N_* expression (± SDm) in populations treated with 0, 2, 4, 6, 8, 10, 12, 14, and 16 μg/mL Amp at t = 12 hrs (blue points) with linear regression fit overlaid (black solid line). These experimental data were compared with the predicted mean GFP-CAT molecules per cell for inducible promoter model simulations at same time point (red points).

In parallel, we observed and modelled EGE behaviour in the context of constitutive expression of the *gfp-cat_T172A_* cassette, where we again saw a dose-dependent unimodal increase in *gfp-cat_T172A_* expression (Supplementary Data Fig. 4 and 5, Video 2). We found a similar theoretical banding pattern of maximal EGE, based on dependence between enzymatic activity of the fitness gene and the applied pressure (Fig 3b). Next, we were able to isolate and characterise dose-dependent EGE within an entirely unrelated fitness model system previously reported by Tsuru et al.^12^ Here, antibiotic exposure was replaced by histidine auxotrophy, which was relieved by expression and upregulation of a histidine biosynthesis pathway gene. Deletion of the native histidinol-phosphate aminotransferase *hisC*, and subsequent rewiring of the strain to encode a monostable *hisC-gfp* circuit, allowed for the uncoupling of *hisC* expression from its native operon. This resulted in a fully synthetic and tunable expression system. We applied a range of fitness pressures by gradually reducing the availability of histidine in the medium; we found a corresponding concentration-dependent stochastic upregulation of the *hisC* gene, again demonstrating ‘gene expression according to need’ (Supplementary Data Fig. 6). By coupling these experimental results to our theoretical framework, we were able to model this behaviour and predict an island of maximal emergence by modulating HisC enzymatic activity *in silico* (Fig 3c). Lastly, using our approach of varying fitness gene product activity, we identified dose dependent EGE in a bactericidal antibiotic resistance context. Here, cells expressing the wild-type or mutant (L74N) beta-lactamase (*bla*) gene from a plasmid template were treated with increasing amounts of ampicillin. In line with our findings in the chloramphenicol resistance framework, expression of wild type *bla* increased slightly with corresponding increase in Amp concentration, while expression of the mutant *bla*_L74N_ was more pronounced at much lower concentrations of the antibiotic (Supplementary Data Fig.7). Next, we derived a dose-responsive banding pattern of maximal EGE of *bla* through adjusting our modelling parameters to take into account the bactericidal activity of the beta-lactamase and its expression levels which allowed us to identify our experimental case within this map (Fig 3d). Thus, we report fitness-induced EGE and corresponding mathematical model banding patterns based on expression strength and fitness gene activity for both plasmid and genome integrated expression systems, in the case of bacteriostatic and bactericidal antibiotics as well as auxotrophy supplementation, indicating the general nature of this population-level behaviour.

Our mathematical and experimental results show that EGE is a global phenomenon that can be observed with some degree of tuning of the relevant parameters and likely plays a role in the expression of any fitness-conferring gene. It should be noted that even small changes in the expression of highly-active fitness genes, such as those encoding antibiotic resistance enzymes, may be crucially important for cells’ biology even where transient EGE may be difficult to detect with current experimental methods^24^. Mathematical modelling reveals these relationships and indeed confirms that an intermediate-activity mutant maximises EGE and gives a model output that fits remarkably well with the experimental Ohm’s law-like linear framework of dose dependent gene expression (Fig. 3 a-d). This can be observed within inducible and constitutive expression systems and across different fitness-conferring genes (Fig 3 e-h). Overall, our results indicate that the linear relationship between fitness pressure and gene expression relies on growth positive feedbacks, stochasticity and fitness gene activity. We propose that evolution could increase population survival in the presence of such stresses as antibiotics or cancer-targeting drugs, by tuning the parameters of a cellular system in the operating regime of EGE; this increases fitness through need-based expression and precludes the necessity for hardwired genetic changes.

## Materials and methods

### Cell growth

All MK01 *E. coli* strains were cultured in lysogeny broth (LB) and, when indicated, 0.005% (w/v) L-(+)-arabinose (Sigma) (stock concentration 5% (w/v) in water) and chloramphenicol (Cm) (Sigma, stock concentration 50mg/ml in ethanol) of ampicillin (Amp) (Sigma, stock concentration 100mg/ml in water), which were stored at −20°C and added to the medium at the beginning of each experiment. Histidine was purchased as a 100mM solution from Sigma. OSU11 and OSU12 *E. coli* strains were a gift from Saburo Tsuru and were cultured in M63 minimal media and supplemented with histidine as described previously^12^. Chloramphenicol, ampicillin and histidine were diluted so that 2μl of an intermediate concentration was added to 148μl of cells in growth media per well in 96-well plates.

All experiments were inoculated from 5ml overnight cultures grown in LB without antibiotics at a starting Absorbance_600_ (A_600_) of 0.01 as measured on the Tecan F200 PRO microplate reader. For experiments in OSU11 and OSU12 *E. coli* strains, the overnight culture in LB was inoculated into M63 media supplemented with histidine. The microplate reader format was used for all experiments; cells were grown at 37°C with orbital shaking at 335.8 RPM with an amplitude of 1.5mm. For the Rounds experiment, 2μl of each culture, grown for 12hrs, was transferred to 148μl of fresh LB media (± arabinose and Cm as indicated) to generate an exact replica of the parent plate with the diluted cultures, which were then growth again under the same conditions for 12hrs.

### Cassette and strain construction

The *catI* gene was cloned from pTKIP-cat which was a gift from Edward Cox & Thomas Kuhlman (Addgene plasmid # 41065; http://n2t.net/addgene:41065; RRID:Addgene 41065). Gibson assembly (NEB) was used for all cloning steps, and all constructs were transformed into Top10 *E. coli* (Invitrogen). The Biobrick part B0034 RBS-*gfp*-Gly4Ser-*cat* construct was inserted downstream of the AraC-pBad cassette in the pCola vector; pCola and GFP were obtained from Schaerli et al. 2014^25^. The *bla* gene was cloned from pUC18 which was a gift from Joachim Messing (Addgene plasmid # 50004; http://n2t.net/addgene:50004; RRID:Addgene 50004) and inserted to replace the *cat* gene downstream of *gfp* in the pBad-inducible expression cassette described above. Lox sites were inserted to flank a kanamycin resistance gene which was then cloned downstream of the GFP-CAT cassette to aid in selection of genomic integrations. For constitutive expression, the AraC gene and pBad promoter were replaced by the Biobrick promoter J23100 upstream of the GFP-CAT-lox-kan-lox cassette in the pCola vector. Site directed mutagenesis was performed to generate the *cat*-T_172_A and H_193_A and *bla*-L_74_N mutants within the expression cassettes. All expression cassette sequences used in this study are reported in Supplementary Fig. 26. A set of plasmids for the main constructs, along with maps and sequences, were deposited in Addgene (IDs are listed in **Supplementary Table X**).

The expression cassettes containing the *cat* gene were integrated into the *intC* locus of the *E. coli* strain MK01 (genotype: F-, Δ(araD-araB)567, ΔlacZ4787(::rrnB-3), λ-, Δ(araH-araF)570(::FRT), ΔaraEp-532::FRT, φPcp8-araE535, rph-1, Δ(rhaD-rhaB)568, hsdR514, ΔlacI)^26^. The strain was modified to decrease biofilm formation by knocking out the *flu* and *fim* genes. Briefly, MK01 cells were transformed with the pRed/ET expression plasmid (Gene Bridges kit K006). Transformants were grown up and recombinase expression was induced as described previously^27^. A kanamycin resistance cassette flanked by lox sites was amplified using primers containing sequences homologous to the 5’ and 3’ regions of the *flu* gene and electroporated into the recombinase expressing MK01 cells. Recombinants were selected on LB-agar containing 15μg/ml kanamycin (Sigma). Successful integration was confirmed via amplification and sequencing of the *flu* locus. The kanamycin resistance cassette was removed by transforming the cells with *Cre* recombinase (Gene Bridges, 706-Cre) according to the manufacturer’s instructions. This sequence was repeated in order to remove the *fim* locus, and subsequently, to introduce the various CAT-GFP expression cassettes into the *intC* locus. Genomic integration and sequencing verification primers used in this study are reported in Supplementary Table 2. All *intC* locus integrations were sequenced to ensure correct integration. Plasmids containing the expression cassettes containing the wild type and mutant *bla* gene were transformed into Top10 cells (Invitrogen) and treated with various Amp concentrations as described above.

### Fluorescence and Absorbance measurements

All experiments where GFP and RFP fluorescence and Absorbance_600_ (A_600_) were measured were performed in 96-well PS, flat bottom, μclear, black plates (Greiner Bio-One), with n=3 technical replicates per treatment, and n=3 biological replicates unless stated otherwise. GFP fluorescence was measured at ex485nm/em535nm and a gain value of 25. Measurements were taken at 15min intervals.

### Flow cytometry

MK01 cells carrying the *gfp-cat-T_172_A* genomic integration cassette were induced with 0.005% arabinose and treated with 0 and 5μg/ml Cm, and grown using the microplate reader with GFP and A_600_ measurements performed as above. At 12hr, 4% paraformaldehyde (Sigma) in PBS was added to each well to a final concentration of 2%, and pipetted up and down to mix. Cells were stored at 4°C in the dark for one to three days. Flow cytometry was performed on a BD Fortessa Analyzer (BD Biosciences) and sample data was analysed using FlowJo (v10) Software.

### RT-qPCR

MK01 cells carrying the *gfp-cat-T_172_A* genomic integration cassette were induced with 0.005% arabinose and treated with 0 and 5μg/ml Cm and grown using the microplate reader with GFP and A_600_ measurements performed as above with n=4 technical replicates. Cells were harvested at 1hr time intervals for 11hrs. Briefly, 600μl of culture of both 0 and 5μg/ml Cm treatments was removed from the plate, added to 1200μl of RNAprotect (Qiagen) in 2mL Eppendorf tubes, and processed according to the manufacturer’s protocol. All samples were stored at −80°C until the end of the time course. RNA was extracted using the RNeasy mini kit (Qiagen). The extracted RNA was treated with TURBO *DNA-free* Kit (Invitrogen) and cDNA was generated using the SuperScript IV First-Strand Synthesis System (Invitrogen). The LightCycler 480 SYBR Green I Master kit (Roche) was used as the qPCR master mix, and the experiments were performed on the Roche 480 LightCycler Instrument II. Housekeeping genes used in this study include *idnT, hcaT*, and *cysG*^28^, and were used to quantify *gfp* mRNA expression at each time point. The delta-delta Ct method was used to determine differences in gene expression, with significance determined using the unpaired Student’s t-test. The mean Ct value of the housekeeping genes was also used to normalise Ct values of the control genes *rpoD, rpoH, rpoE, rpoN, acrB, pntB, oppA*, and *cyoC*. Primer sequences used for pPCR amplification are reported in Table 3 in Supplementary Information.

### Mass Spectrometry

100μl of culture medium was mixed with 100μL of a solution containing a mixture of acetonitrile, methanol and water (40:40:20, v/v/v). After centrifugation at 17,000 x g, at 4°C, for 10 mins, 100μl of the supernatant was mixed 100μl of a solution of acetonitrile containing 0.2% acetic acid. After vortexing and centrifugation at 17,000 x g, at 4°C, for 10 mins, 100μl of the supernatant was loaded into LC-MS vials prior to analysis.

Aqueous normal phase liquid chromatography was performed using an Agilent 1290 Infinity II LC system equipped with binary pump, temperature-controlled auto-sampler (set at 4°C) and temperature-controlled column compartment (set at 25°C), containing a Cogent Diamond Hydride Type C silica column (150 mm × 2.1 mm; dead volume 315 μl). A flowrate of 0.4 ml/min was used. Elution of polar metabolites was carried out using solvent A (0.2% acetic acid in deionized water (Resistivity ~ 18 MW cm), and solvent B (acetonitrile and 0.2% acetic acid). Mass spectrometry was carried out using an Agilent Accurate Mass 6545 QTOF apparatus. Nozzle Voltage and fragmentor voltages were set at 2,000 V and 100 V, respectively. The nebulizer pressure was set at 50 psig and the nitrogen drying gas flow rate was set at 5 l/min. The drying gas temperature was maintained at 300°C. Data were collected in the centroid mode in the 4 GHz (extended dynamic range) mode^29^ and the values were normalized to the starting concentration measurement at 0hrs.

### Mathematical Modelling

We generated agent-based models of the genetic networks that includes stochastic simulation of gene expression inside growing and dividing cells to capture EGE. We used Approximate Bayesian Computation to fit the models to data^17,19^. The full details of the mathematical modelling are found in supplementary information. All the modelling was implemented in the Julia programming language.

## Acknowledgments

We thank Saburo Tsuru for sharing with us the *OSU11* and *OSU12-hisC E. coli* strains, Miles Priestman for technical assistance and Anthony Bowman for sharing the script for ABC-SMC model selection in Julia language. We also thank Martin Buck, Philipp Thomas and Peter Swain for comments on the manuscript. This work was funded by Wellcome Trust UK New Investigator Award No. WT102944 and was supported by a Royal Society International Exchanges Cost Share Award (IE170297).

## Author contributions

Conceived and designed the experiments: MC, MS, MI

Performed the experiments: MC, MS

Analyzed the data: MC, MS, GLM, VS, MI

Contributed reagents/material/analysis tools: GLM, VS, MI

Wrote the manuscript: MC, MS, AG, VS, MI

