## Supplementary material for "Ohm’s Law for increasing fitness gene expression with selection pressure"

3 **Sequence data** **58**

**4 Supplementary videos** **64**

### 1 Supplementary figures

| Model | 1 | 2 | 3 | 4 | 5 | 6 | 7 | 8 | 9 | 10 |
|---|---|---|---|---|---|---|---|---|---|---|
| mRNA | ✓ | ✓ | ✓ | ✓ | ✓ | ✓ | ✓ | ✓ | ✓ | ✓ |
| protein | ✗ | ✗ | ✗ | ✗ | ✓ | ✓ | ✓ | ✓ | ✓ | ✓ |
| biased partitioning | ✗ | ✗ | ✗ | ✗ | ✓ | ✗ | ✗ | ✗ | ✓ | ✗ |
| global transcription feedback | ✗ | ✗ | ✗ | ✓ | ✗ | ✓ | ✗ | ✓ | ✗ | ✗ |
| global translation feedback | ✗ | ✗ | ✗ | ✗ | ✗ | ✗ | ✓ | ✓ | ✗ | ✗ |
| constitutive | ✓ | ✓ | ✓ | ✗ | ✓ | ✓ | ✓ | ✓ | ✓ | ✗ |
| regulated | ✗ | ✗ | ✗ | ✓ | ✗ | ✗ | ✗ | ✗ | ✗ | ✓ |
| no. of parameters | 2 | 3 | 3 | 4 | 3 | 4 | 4 | 5 | 4 | 5 |
| Ohm's Law | ✗ | ✗ | ✗ | ✗ | ✗ | ✗ | ✓ | ✓ | ✗ | ✓ |
| unimodal shift | ✗ | ✗ | ✗ | ✗ | ✗ | ✗ | ✓ | ✓ | ✗ | ✗ |
| mRNA increase | ✗ | ✗ | ✗ | ✓ | ✗ | ✗ | ✗ | ✓ | ✗ | ✓ |
| max ratio | 1.0 | 1.0 | 1.0 | 1.0 | 1.0 | 1.0 | 1.75 | 1.8 | 1.0 | 2.1 |

Supplementary Table 1: The upper part of the table lists the different model assumptions made. A tick indicates that the model (for a given column) contains an assumption (given by the row) while a cross indicates that it does not. The lower part (highlighted in gray) summarises the model output. The first row indicates whether each model is capable of producing a linear relationship between the fitness protein and the fitness pressure. The second row shows whether we can observe a unimodal shift to the right (increase in the production) of the fitness protein in response to stress. The next row shows whether an increase in the mean mRNA level is observed in response to fitness pressure. Finally, the maximum mean ratio of the fitness protein to the reference protein for each model is displayed in the last row. The only model that captures the observed data (unimodal shift, mRNA increase and Ohm's Law) is model 8, which also worked when tested for the regulated case and which we adopt in the main paper.

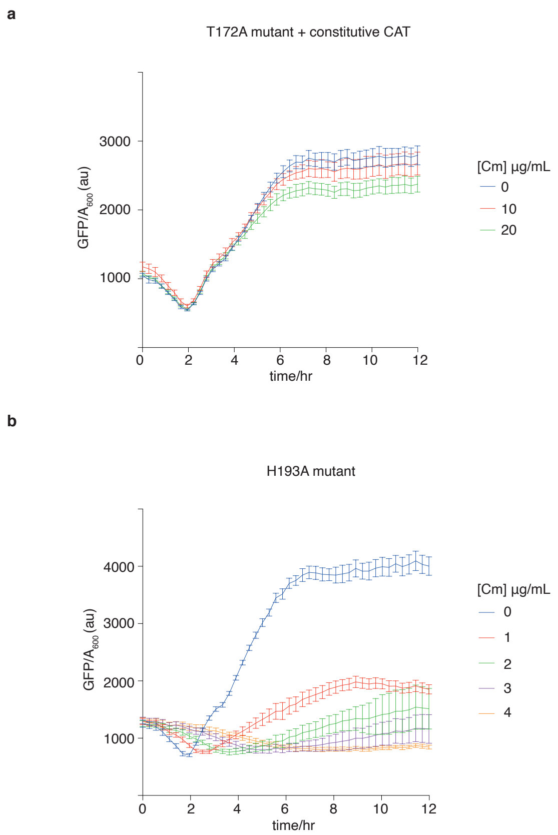

Supplementary Figure 1: Chloramphenicol treatment does not result in increased gene expression of *gfp-cat* cassette under conditions of relieved fitness pressure or inactivity of CAT enzyme. (a) The *E. coli* strain *MK01* intC::pBad-B0034-*gfp-cat-T*$_{172}$*A* was transfected with an extra plasmid constitutively expressing the CAT gene to neutralise the fitness selection pressure on the integrated antibiotic resistance gene. Cells were induced with arabinose and treated as before (Fig. 1f), with 0, 10 and 20 µg/mL Cm, but no longer displayed emergent gene expression. Mean (± SDm) GFP expression (GFP/$A_{600}$ per well) was monitored for 12hrs, representative of n=3 biological replicates. (b) An inactive GFP-CAT is not involved in upregulation of gene expression upon Cm challenge. The *E. coli* strain *MK01 intC::pBad-B0034-gfp-cat-H*$_{193}$*A* (inactive CAT mutant) was induced with 0.005% arabinose and treated with 0, 1, 2, 3, and 4 µg/mL Cm. Mean (± SDm) GFP expression (GFP/$A_{600}$ per well) was monitored for 12 hrs; n=3 biological replicates.

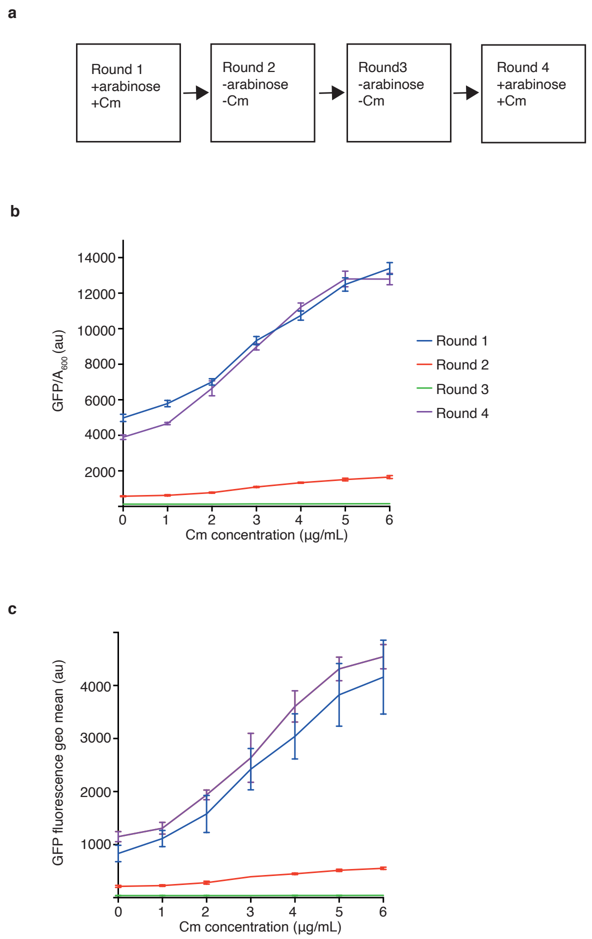

Supplementary Figure 2: Emergent gene expression (EGE) is reversible and repeatable upon rounds of growth after stressor addition, removal, and re-addition. (a) Schematic of experimental design. Round 1, 150 µl of *E. coli MK01* intC::pBad-B0034-*gfp-cat-*$T_{172}A$ culture was induced with 0.005% arabinose and treated with 0,1,2,3,4,5, or 6 µg/mL Cm, to apply different fitness pressures on the GFP-CAT. GFP expression and growth were monitored for 12hrs. Starting cultures for Round 2 were obtained from the 12hr time point wells of the Round 1 experiment, where 2µl of culture was used to inoculate 148µl LB media, now in the absence of arabinose or Cm (to test EGE reversibility). GFP expression and growth were monitored for 12hrs. Starter culture for Round 3 was obtained as above from the 12hr time point of Round 2 and grown in the same manner, in the absence of arabinose or Cm. Round 4 was inoculated from the 12hr time point of Round 3 and cells were cultured while restoring the presence of 0.005% arabinose and 0,1,2,3,4,5, or 6 µg/mL Cm. (b) Experimental results of reversibility experiment showing mean (± SDm) GFP expression (GFP/$A_{600}$ per well), monitored for 12hrs of Rounds 1-4; n=3 biological replicates. (c) Comparable flow cytometry data for the plate reader experiments in (b). Every condition in each round of growth was fixed with 2% paraformaldehyde at 12hrs, analysed by flow cytometry and graphed as the geometric mean of each culture in Rounds 1-4; n=3 biological replicates.

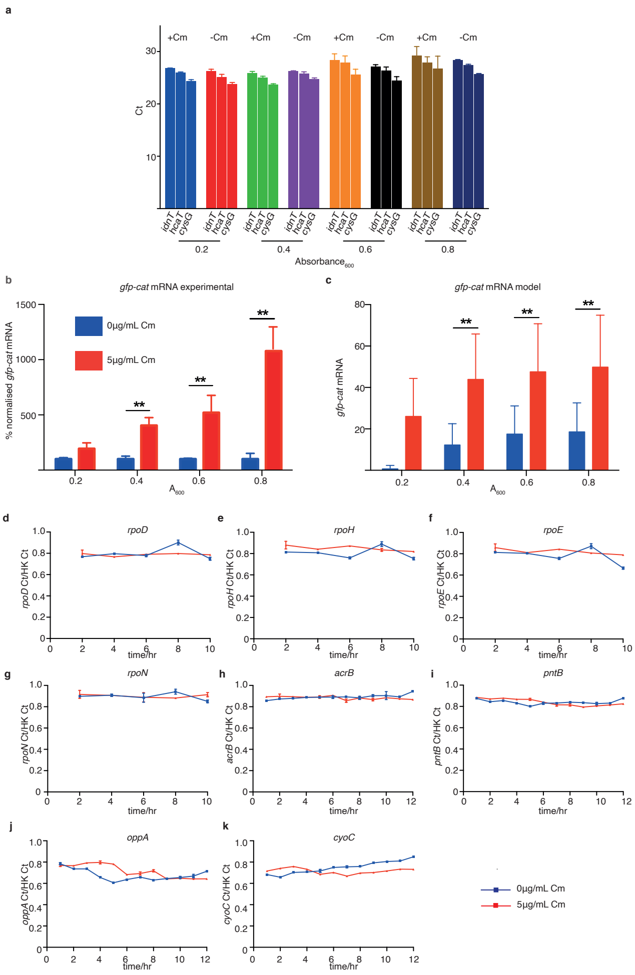

Supplementary Figure 3: Emergent gene expression-associated upregulation is specific to *gfp-cat* mRNA and is predicted by the model without further fitting. RT-qPCR assay showing (a) housekeeping gene *idnT*, *hcaT*, and *cysG* transcript raw Ct values used to analyse *gfp-cat-T172A* expression in cells induced with 0.005% arabinose, treated with 0 or 5 µg/mL Cm and harvested at $A_{600}$ of 0.2, 0.4, 0.6, and 0.8, mean (± SEm, n=3 biological replicates) and (b), *gfp-cat-T172A* expression normalised to the housekeeping genes in (a), Cm concentrations are indicated as 0 (blue) or 5 (red) µg/mL in each sample (± SEm, n=3 biological replicates). Asterisks represent p values: ** = $p<0.01$; delta-delta Ct value analysis was used for statistical analysis and values within comparison groups were normalised to the control 0 µg/mL Cm treatment. (c) *gfp-cat* mRNA values from simulations of the inducible promoter model treated with 0 (blue) or 5 (red) µg/mL Cm for cell populations corresponding to $A_{600}$ of 0.2, 0.4, 0.6 and 0.8. Asterisks represent p values: ** = $p<0.01$; Two-sample Kolmorgorov-Smirnov test was used. mRNA Expression of sigma factors (d) *rpoD*, (e) *rpoH*, (f) *rpoE*, (g) *rpoN* as well as genes with reported potential transcriptional level fluctuations in the presence of antibiotics [44] including (h) *acrB*, (i) *pntB*, (j) *oppA*, and (k) *cyoC*. Ratios of transcript Ct values to housekeeping transcript Ct values are reported (± SDm, n=1 biological replicate).

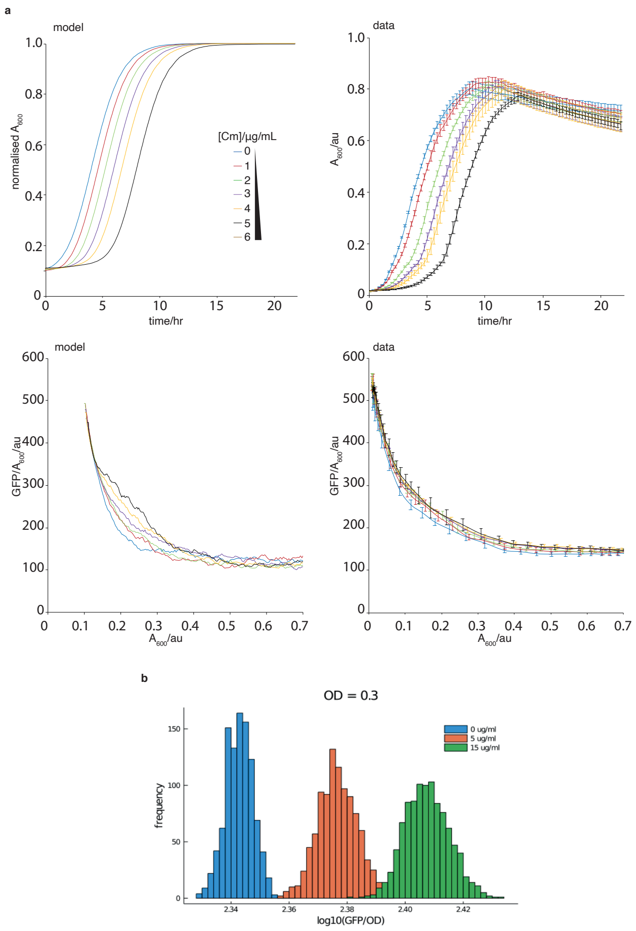

Supplementary Figure 4: Constitutive expression model of emergent gene expression (promoter BBaJ23100 with mutant $T_{172}$A GFP-CAT). (a) Graphs show mean (± SDm) GFP expression (GFP/$A_{600}$ per well) in populations constitutively expressing genome-integrated mutant *gfp-cat*$_{T172A}$. These were treated with 0-6 μg/mL Cm, to induce EGE, and monitored for 22 hrs; n=3 biological replicates. Time series showing total number of cells from simulation of a constitutive promoter model normalised by carrying capacity (upper left panel), corresponding experimental mean (± SDm) GFP expression (GFP/$A_{600}$ per well) in populations constitutively expressing genome-integrated mutant *gfp-cat*$_{T172A}$. (upper right panel), mean number of GFP-CAT molecules per cell from simulation of the constitutive promoter model (lower left panel) and corresponding experimental mean *gfp* expression data (lower right panel), for cells treated with 0-6 μg/mL Cm, for a time period of 22 hrs. Best fit parameters were used from ABC parameter inference and initial conditions (initial number of cells and *gfp* expression levels) are takenwere taken from data displayed in right hand side panels. mRNA levels were assumed to be zero initially. (b) Histograms show 1000 samples of Gaussian processes at $A_{600}$/au=0.3 which were fitted (via maximisation of log-likelihood) to GFP/$A_{600}$ and $A_{600}$ data presented in right panels of (a) for 0, 5 and 15 μg/mL Cm. GFP/$A_{600}$ curves were found to increase with Cm dosage in a statistically significant manner (using Student's T-test).

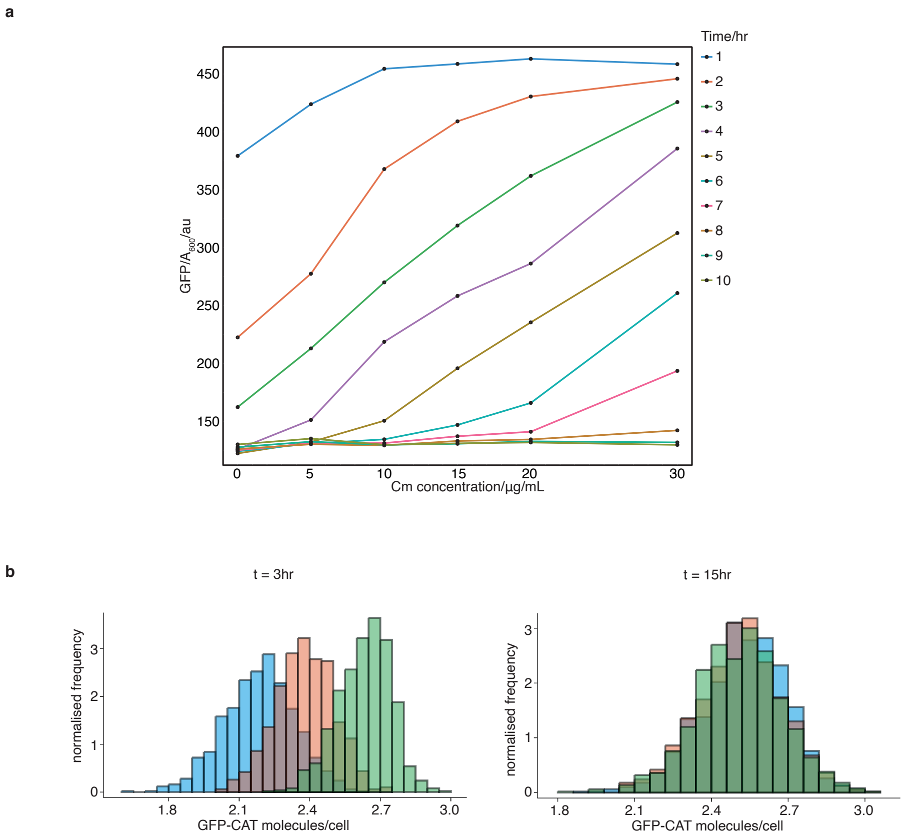

Supplementary Figure 5: Transient upregulation of *gfp-cat* observed under experimental and model conditions (constitutive emergent gene expression model). (a) The *E. coli* strain *MK01 intC::J23100-B0034-GFP-CAT*$_{T172A}$ was treated with 0-6 μg/mL Cm and mean (± SDm) GFP expression (GFP/$A_{600}$ per well) was monitored for 10hrs, representative of n=3 biological replicates, with each line corresponding to a different time point. Linear regression analysis of these lines reveals linearity is maximised at t = 3hrs. (b) Left panel shows GFP-CAT distributions from simulations of the constitutive promoter model at t = 3 hr, displayed in histograms for 2, 4 or 6 μg/mL Cm (coloured in blue, red and green respectively). Right panel shows GFP-CAT distributions from simulations of constitutive promoter model at t = 15 hr displayed in histograms for 2, 4 or 6 μg/mL Cm (coloured in blue, red and green respectively). X-axes are displayed on log10 scale and y-axes are scaled such that the total area of the histograms sum to 1.

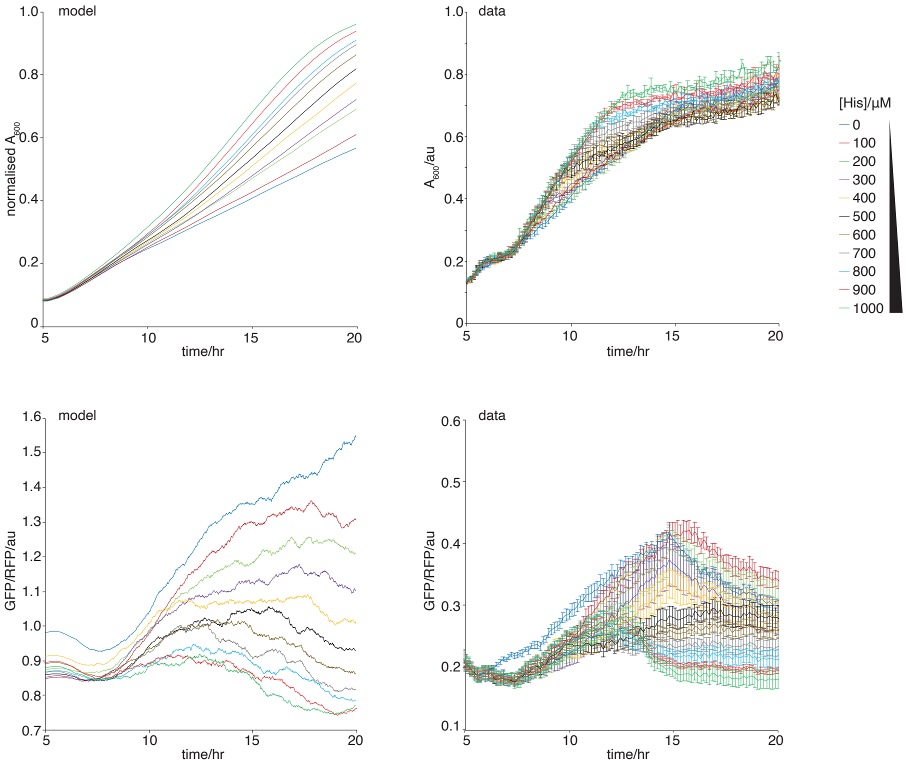

Supplementary Figure 6: Inducible expression model of emergent gene expression of *gfp-hisC* in *E. coli OSU12-hisC* grown in decreasing concentrations of histidine in the growth medium. Time series showing total number of cells from simulations of an inducible promoter model normalised by carrying capacity (upper left panel) and corresponding experimental mean $A_{600}$ data (upper right panel) for 11 histidine conditions ranging from 0 µM to 1000 µM for a time period of 20 hours. Lower plots show the corresponding time series of *gfp* expression normalised to *rfp* expression for the same conditions with the left plot showing the model output and the right plot showing the experimental mean. Best fit parameter were used from ABC parameter inference and initial conditions (initial number of cells, *gfp* expression and *rfp* expression levels) were taken from data displayed in right hand side panels. mRNA levels were assumed to be zero initially.

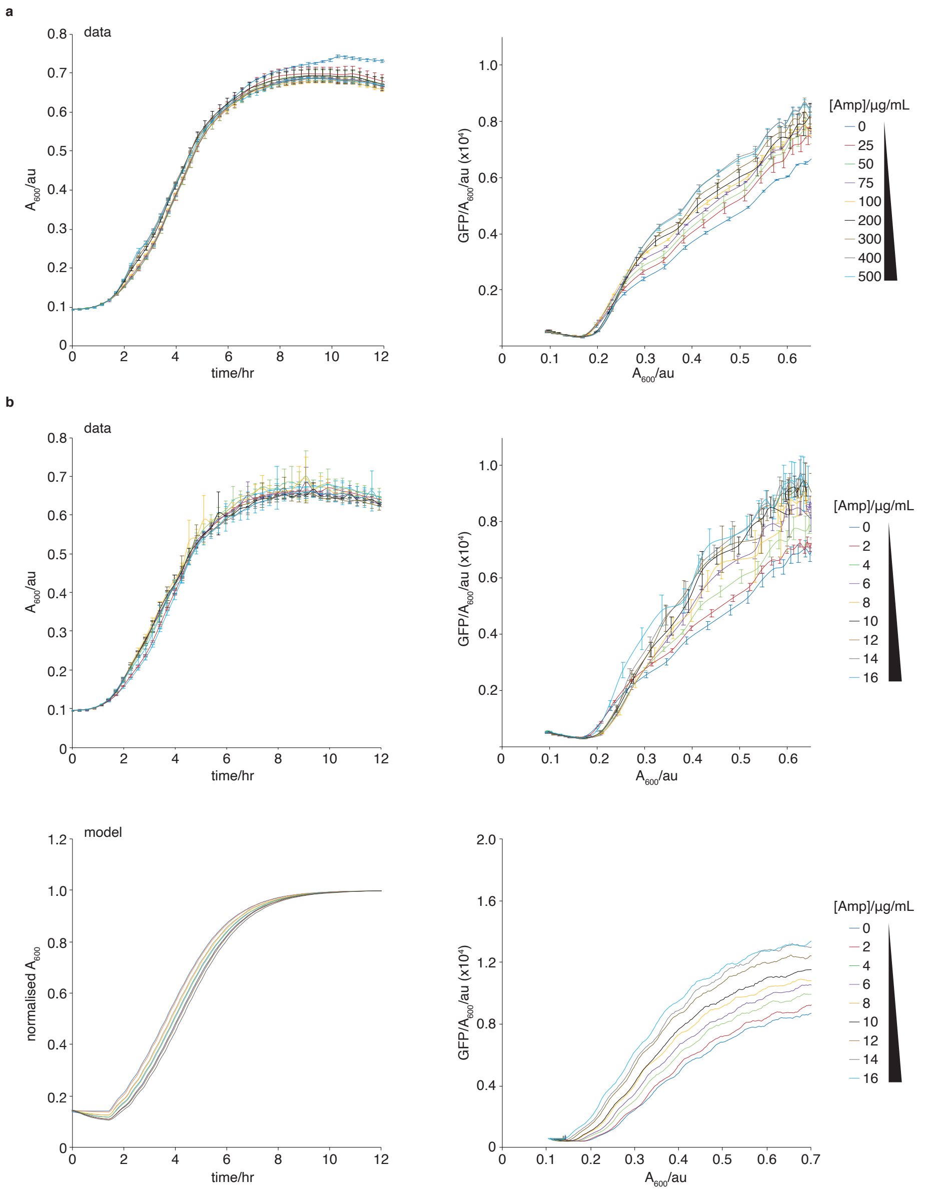

Supplementary Figure 7: Emergent gene expression (EGE) increases linearly with rising amplicillin concentration. (a) Emergent wild type *gfp-bla* expression with a weakly-induced $P_{BAD}$ promoter (0.005% arabinose). Graphs show growth kinetics $A_{600}$ (left panel) and GFP fluorescence per well (right panel) of populations expressing plasmid-encoded wt *gfp-bla* and treated with 0, 25, 50, 75, 100, 200, 300, 400, and 500 µg/mL Amp. (b) Mathematical model and experimental data of growth kinetics $Abs_{600}$ (upper panels) and GFP fluorescence per well of populations (lower panels) expressing plasmid-encoded mutant *gfp-bla74N* and treated with 0, 2, 4, 6, 8, 10, 12, 14, and 16 µg/mL Amp; n=3 biological replicates. Best fit parameters were used from ABC parameter inference and initial conditions (initial number of cells and *gfp* expression levels) were taken from data displayed in top panels of b. mRNA levels were assumed to be zero initially.

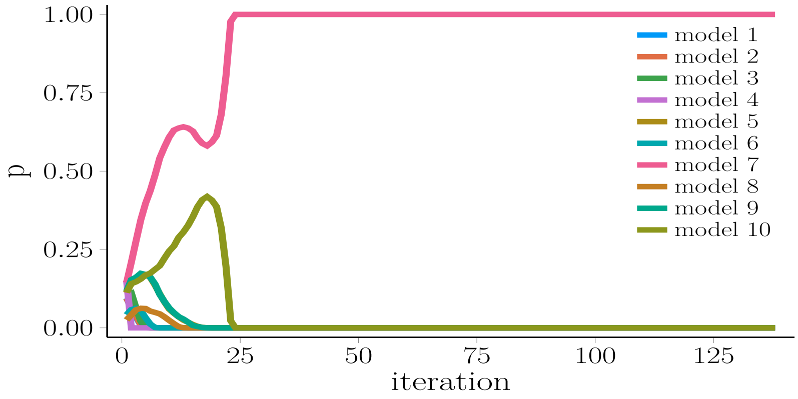

Supplementary Figure 8: Bayesian model selection result. Probability of models 1 to 10 varying with iteration of ABC-SMC algorithm. Model 7 is the most probable model.

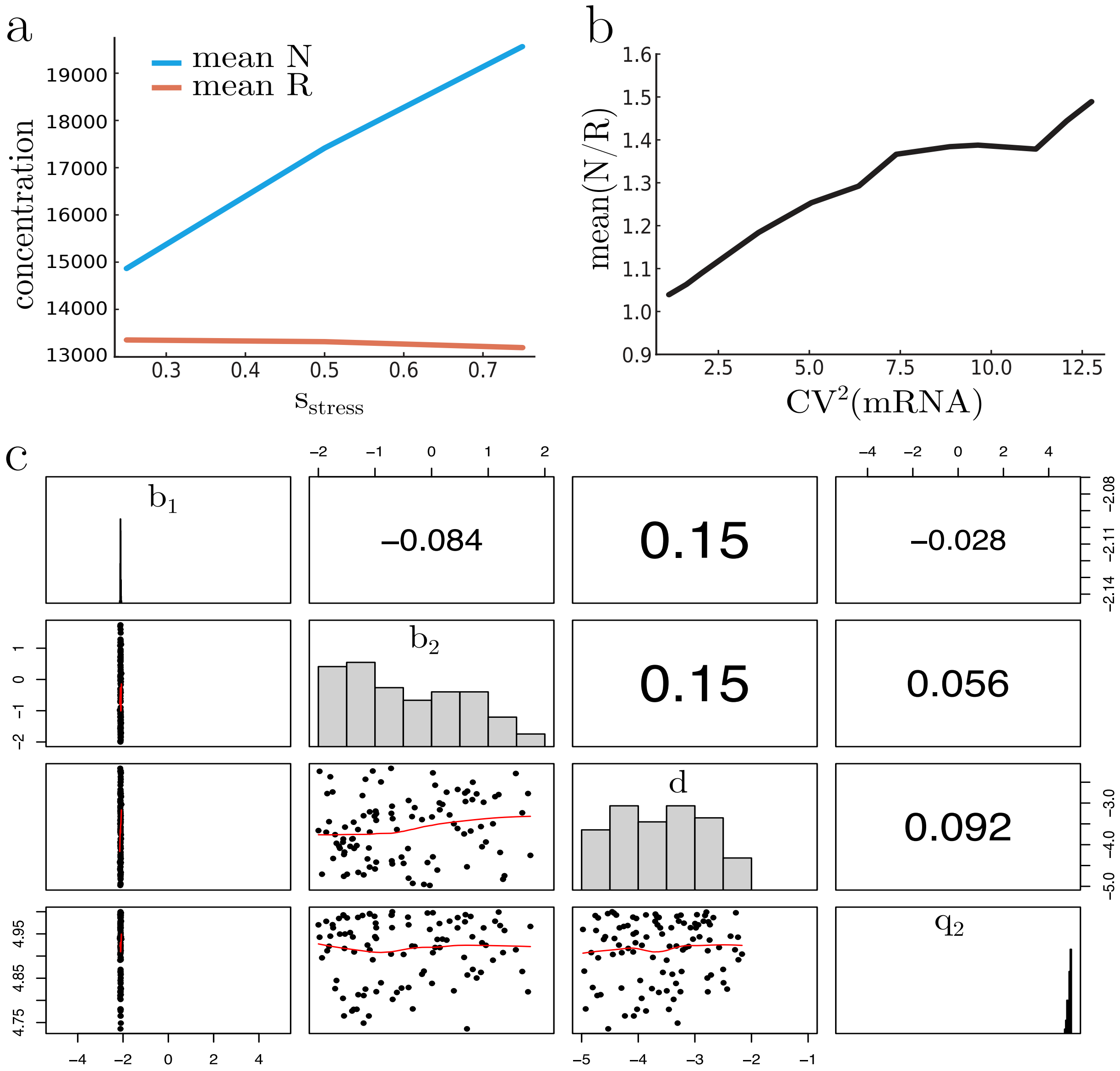

Supplementary Figure 9: Summary of ABC parameter study of model 7 using error defined in equation (8). (a) relationship between stress level and steady state concentration of protein species in model 7. Concentration is defined as the protein level divided by volume of cell. (b) relationship between steady state ratio of fitness protein and reference protein and mRNA noise level as defined by coefficient of variation squared. (c) final posterior distributions of model 7 parameters. X-axes limits correspond to uniform prior distribution ranges.

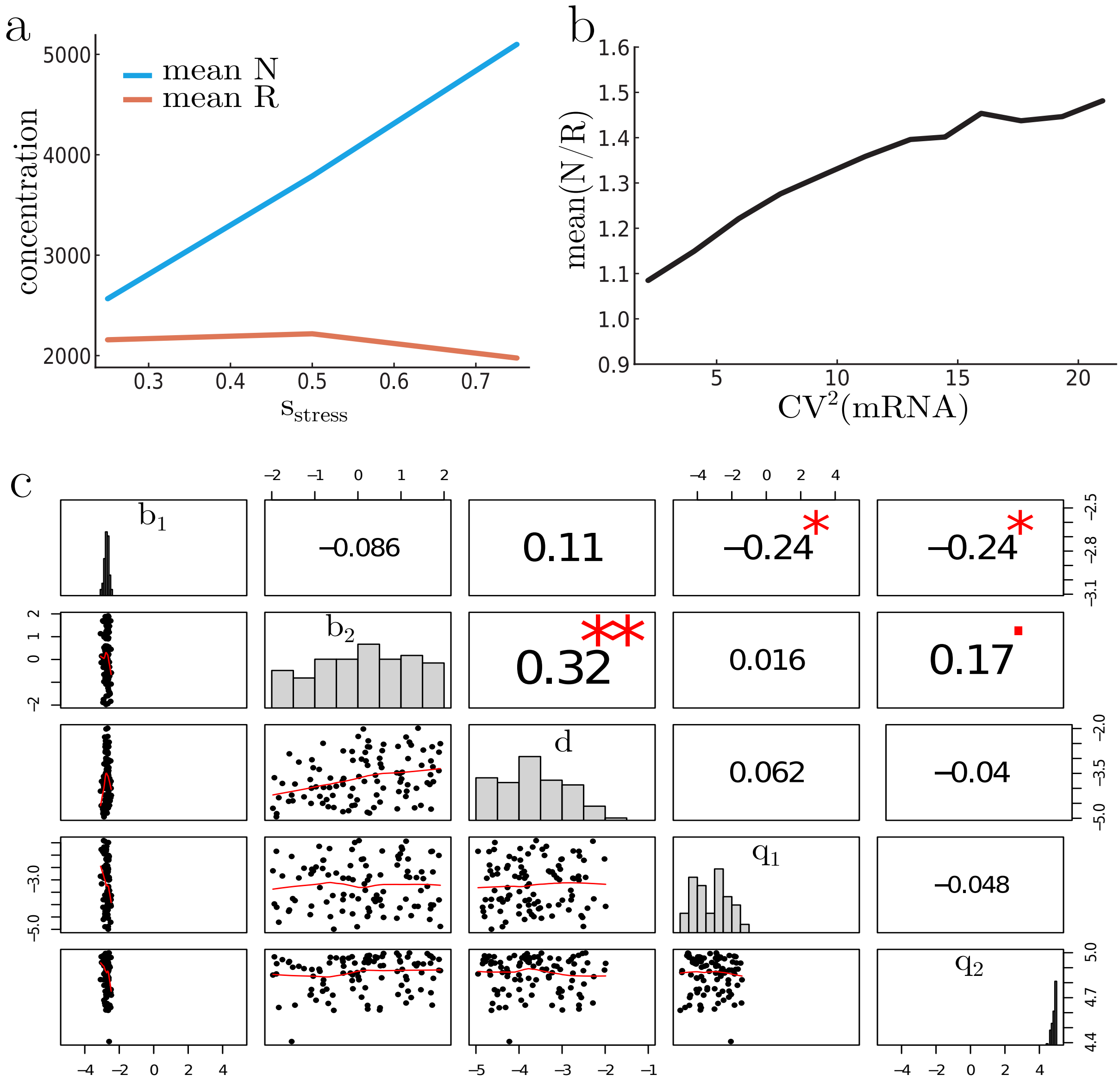

Supplementary Figure 10: Summary of ABC parameter study of model 8 using error defined in equation (8). (a) relationship between stress level and steady state concentration of protein species in model 8. Concentration is defined as the protein level divided by volume of cell. (b) relationship between steady state ratio of fitness protein and reference protein and mRNA noise level as defined by coefficient of variation squared. (c) final posterior distributions of model 8 parameters. X-axes limits correspond to uniform prior distribution ranges.

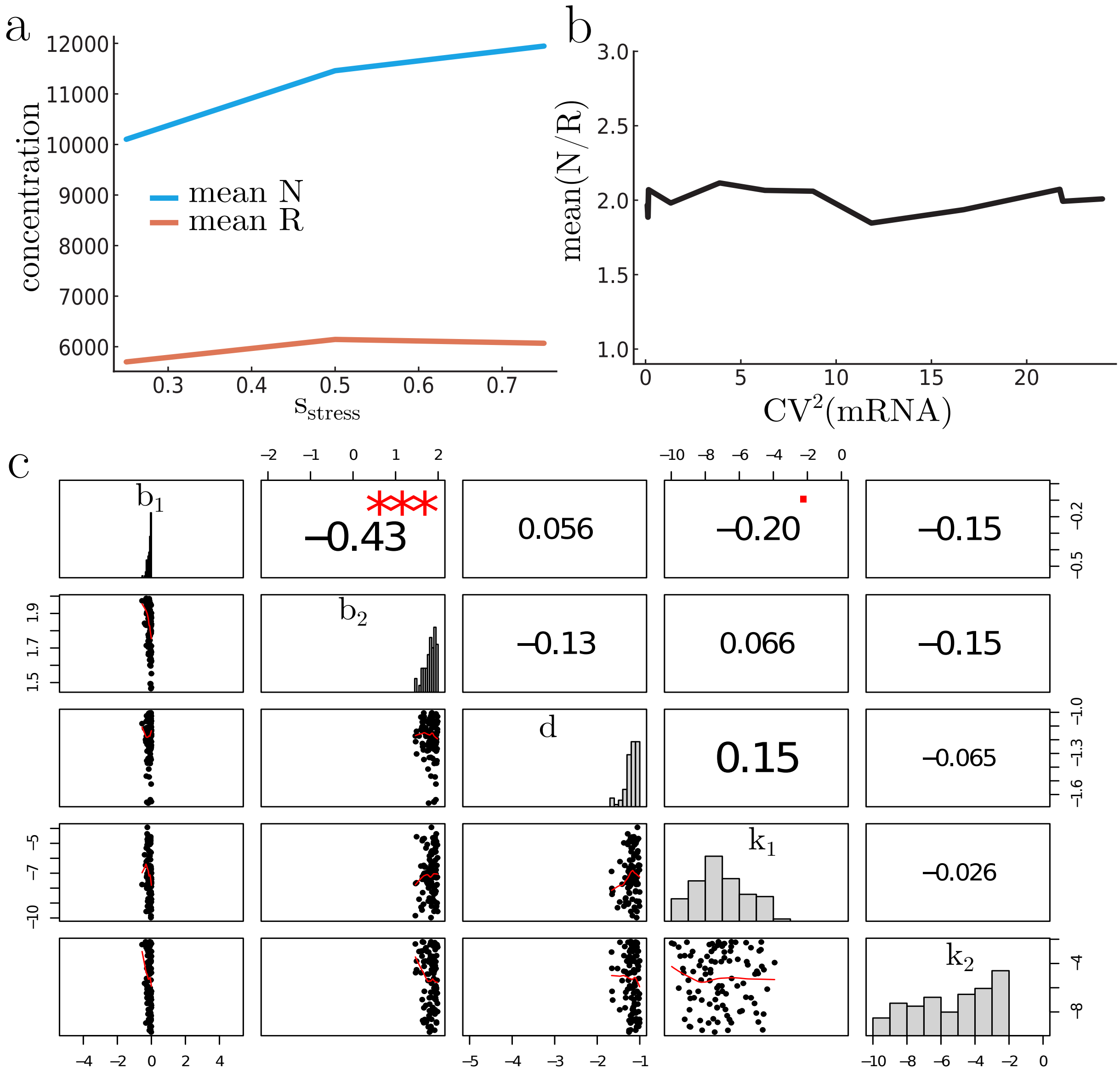

Supplementary Figure 11: Summary of ABC parameter study of model 10 using error defined in equation (8). (a) relationship between stress level and steady state concentration of protein species in model 10. Concentration is defined as the protein level divided by volume of cell. (b) relationship between steady state ratio of fitness protein and reference protein and mRNA noise level as defined by coefficient of variation squared. (c) final posterior distributions of model 10 parameters. X-axes limits correspond to uniform prior distribution ranges.

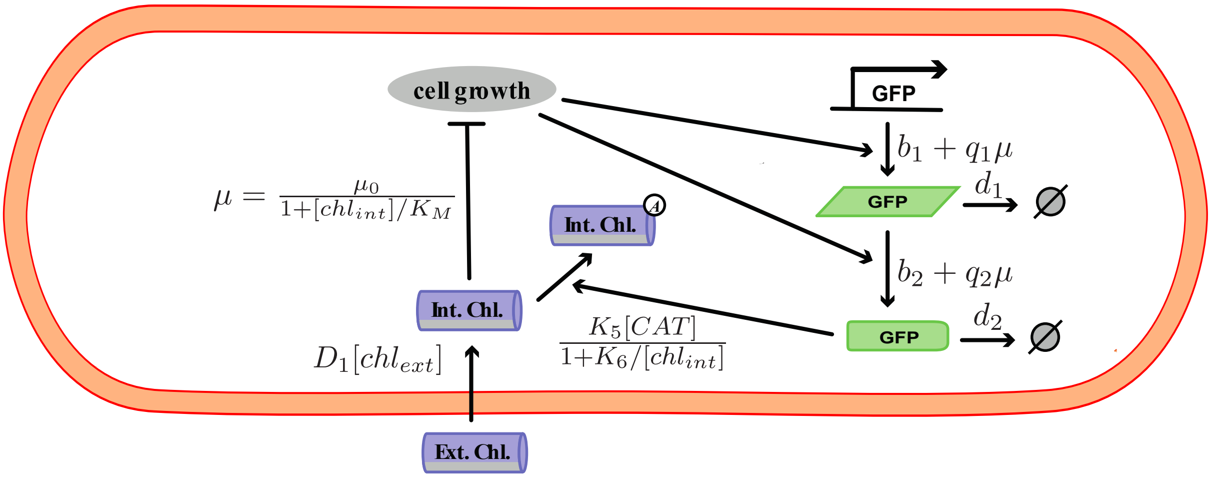

Supplementary Figure 12: Schematic summarising constitutively expressed promoter model. Square brackets denote concentration, i.e., the species is divided by the cell volume, V. For further details on individual reactions, see text. Parallelograms represent mRNA species and rounded rectangles represent protein species.

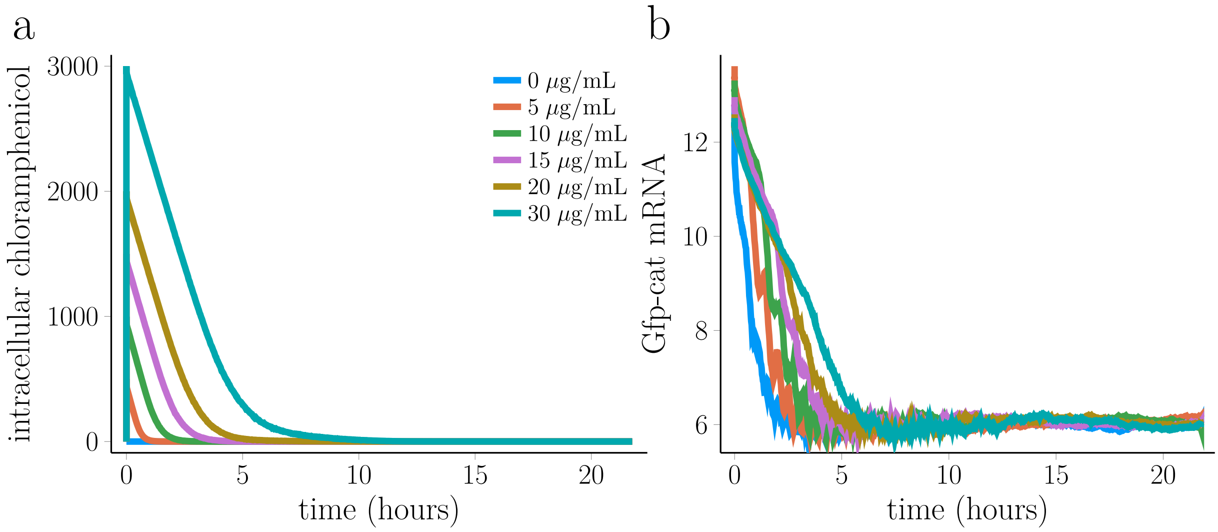

Supplementary Figure 13: Simulations of constitutive promoter model defined in section 1.6. Parameters used are obtained from best fit of microplate reader data shown in Supplementary Data Figure 3. (a) mean intracellular chloramphenicol levels varying in time over a period of 21 hours for 6 different doses of chloramphenicol defined in legend. (b) mean Cat mRNA levels varying in time over a period of 21 hours for 6 different doses of chloramphenicol.

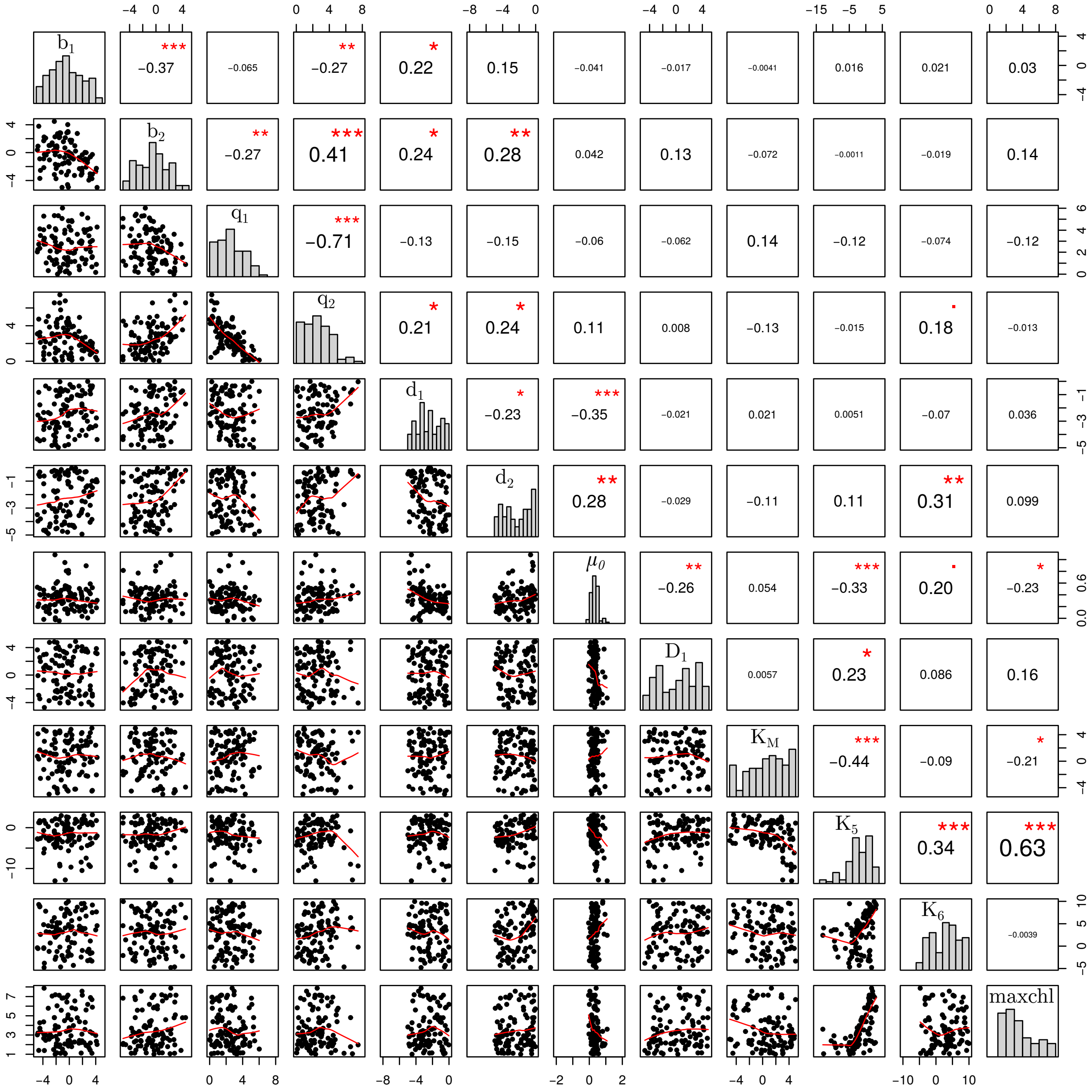

Supplementary Figure 14: Posterior parameter distributions found by fitting the constitutive promoter model to microplate reader data presented in Supplementary Data Figure 3. Limits on x-axes correspond to ranges of Uniform prior distributions used to initialise ABC inference. Lower triangular plots show scatter plots of parameter distributions with lowess smoothed line overlaid. Upper triangular plots show pearson correlation with a point representing a p-value less than 0.05, an asterisk representing a p-value less than 0.01, two asterisks representing a p-value less than 1e-3 and three asterisks representing a p-value less than 1e-4.

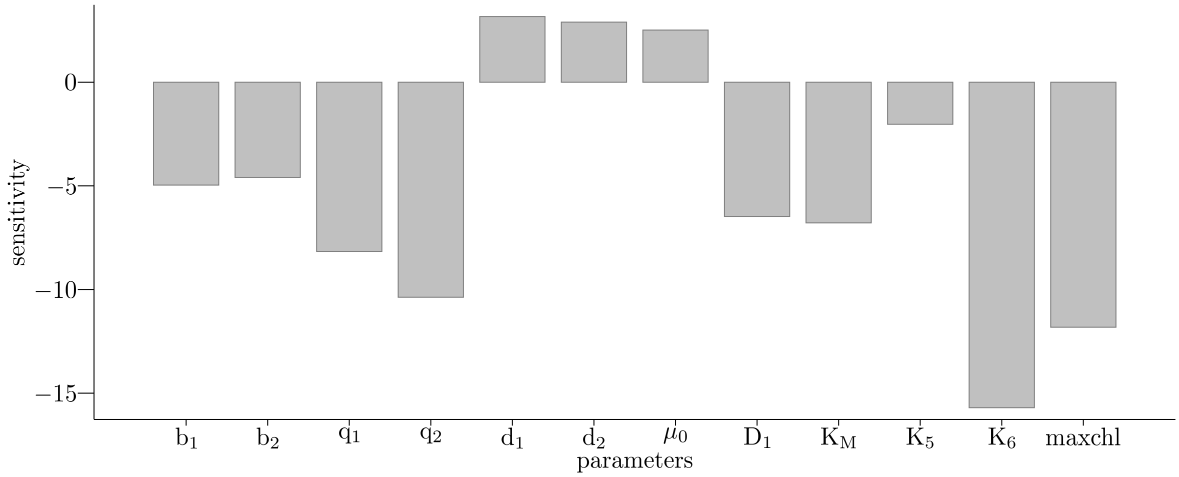

Supplementary Figure 15: Sensitivity of parameters for model fitted to microplate reader experiments shown in Supplementary Data Figure 3 as computed by inverting the covariance matrix of the final probability distribution shown on the diagonal of Supplementary Figure 14. The y-axis is plotted on a log-scale and the parameters on the x-axis are explained in section 1.6.

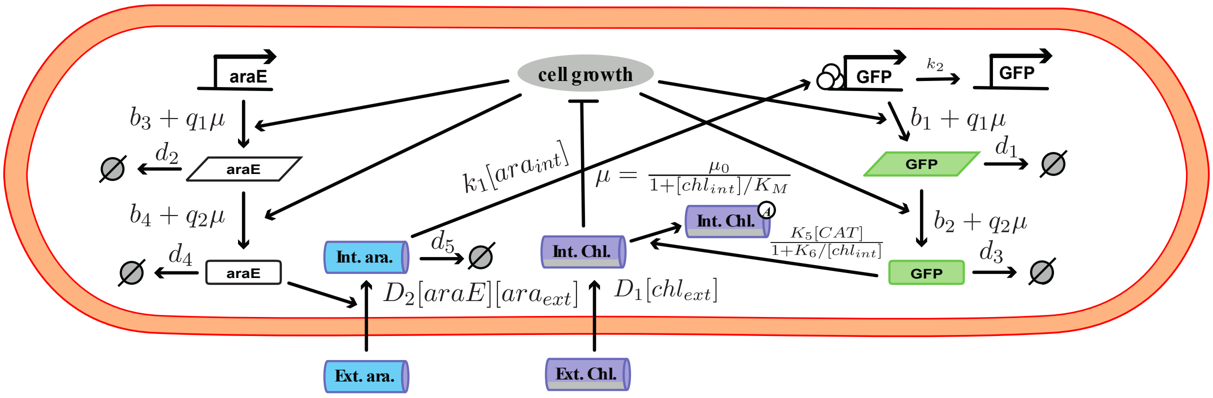

Supplementary Figure 16: Schematic summarising inducible promoter model. Square brackets denote concentration, i.e., the species is divided by the cell volume, V. For further details on individual reactions, see text. Parallelograms represent mRNA species and rounded rectangles represent protein species.

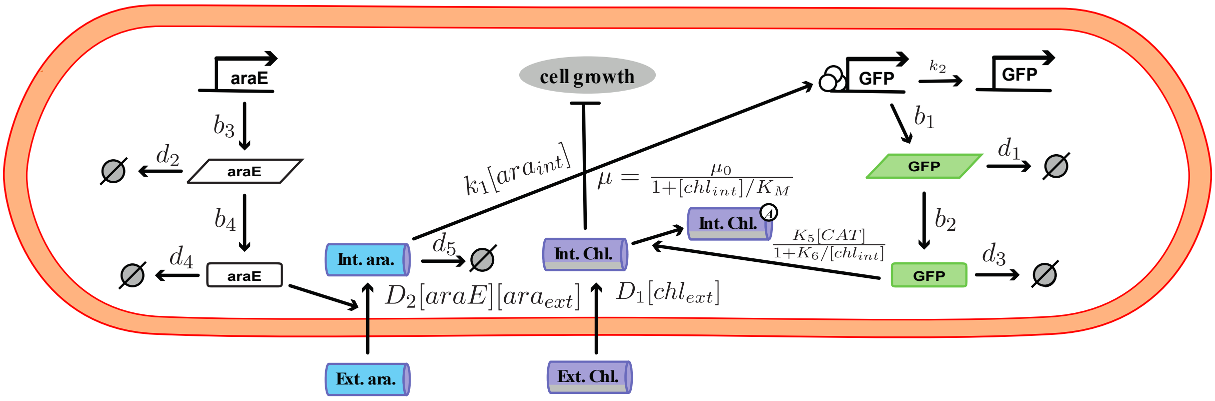

Supplementary Figure 17: Schematic of control arabinose inducible model. This model is unable to capture the same dynamics as the previous one, this is used to ensure the same underlying process is happening as in the constitutive csae (positive feedback mediated selection).

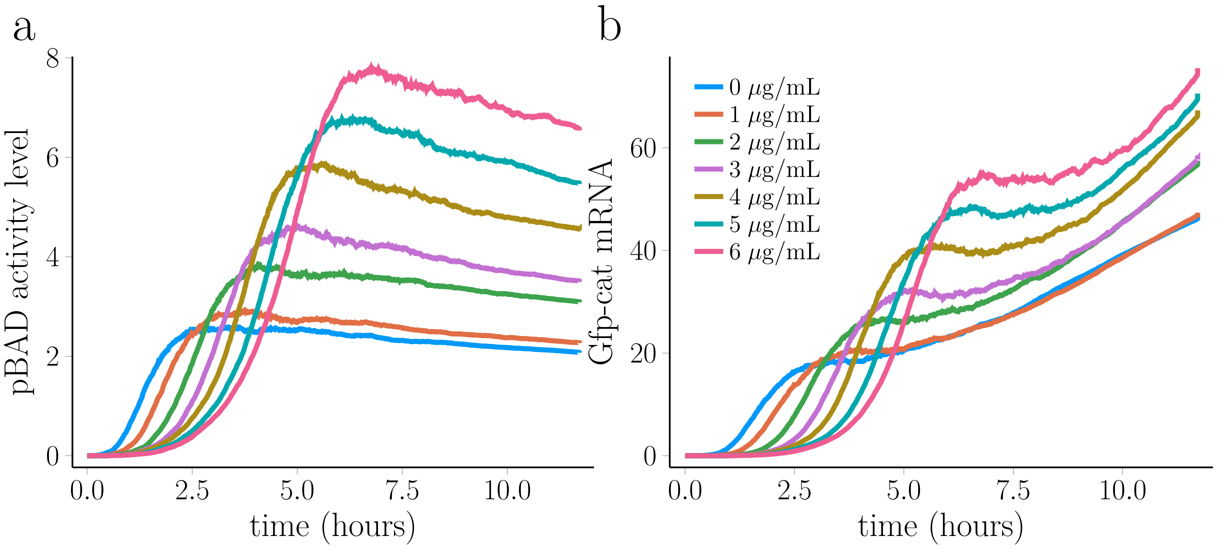

Supplementary Figure 18: Simulations of inducible promoter model defined in section 1.7. Parameters used are obtained from best fit of microplate reader data shown in Figure 2b of main paper. (a) mean pBAD activity levels varying in time over a period of 12 hours for 7 different doses of chloramphenicol defined in legend. (b) mean Cat mRNA levels varying in time over a period of 12 hours for 7 different doses of chloramphenicol.

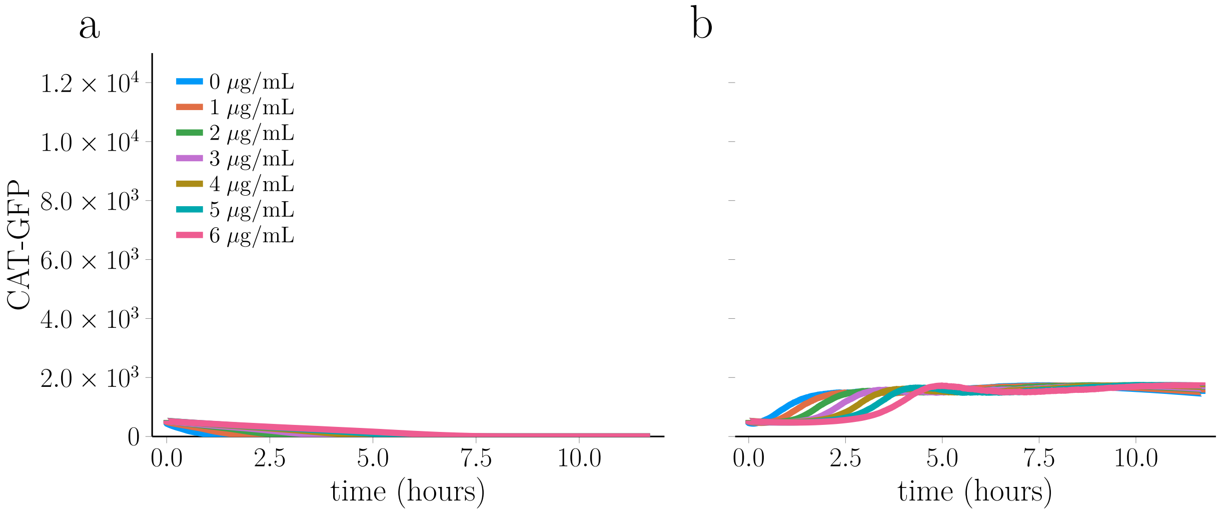

Supplementary Figure 19: Control simulations of inducible promoter model defined in section 1.7. Parameters used are best fit parameters from fitting to microplate reader data shown in Figure 2 of main paper unless otherwise stated. (a) repeating simulations with transcriptional and translational global positive feedbacks removed, i.e., with parameters $q_1$ and $q_2$ set to zero. (b) repeating simulations with pBAD activity levels fixed to a constant, 1.

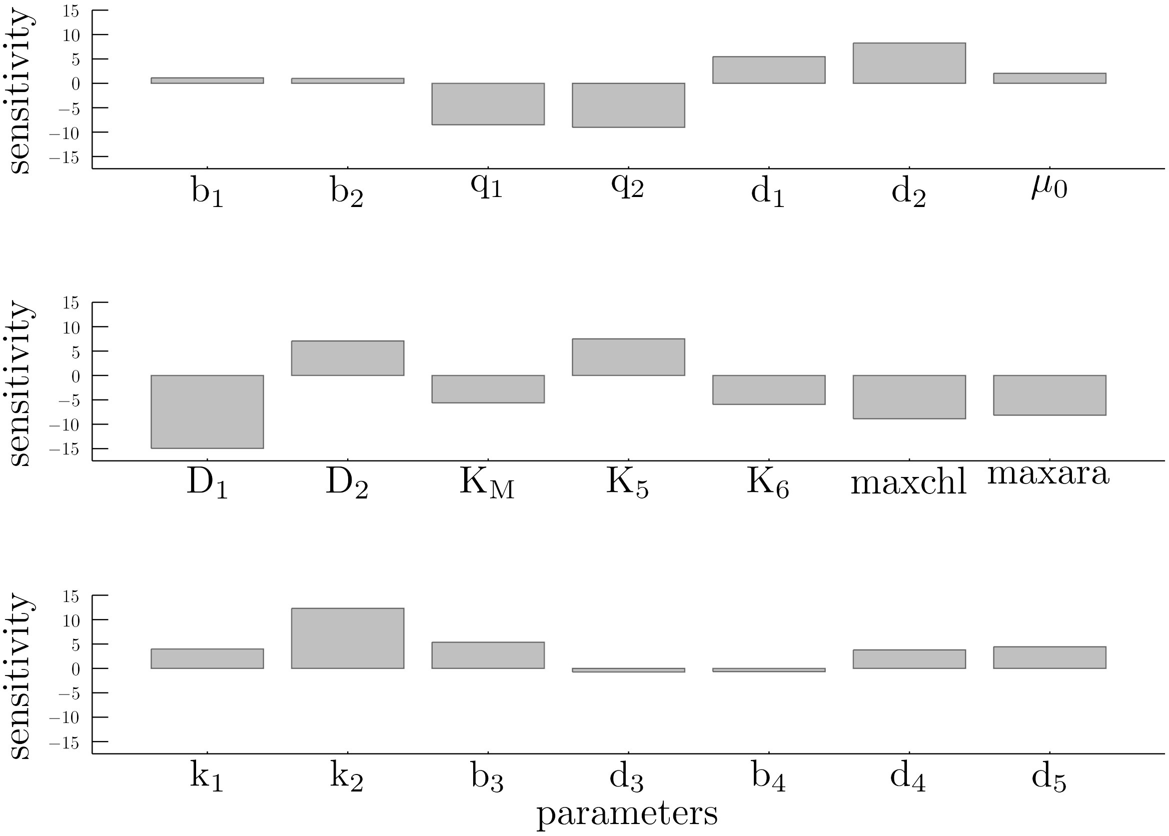

Supplementary Figure 20: Sensitivity of parameters for model fitted to microplate reader experiments shown in Figure 2b of the main paper as computed by inverting the covariance matrix of the final probability distributions obtained from ABC. The y-axis is plotted on a log-scale and the parameters on the x-axis are explained in section 1.7.

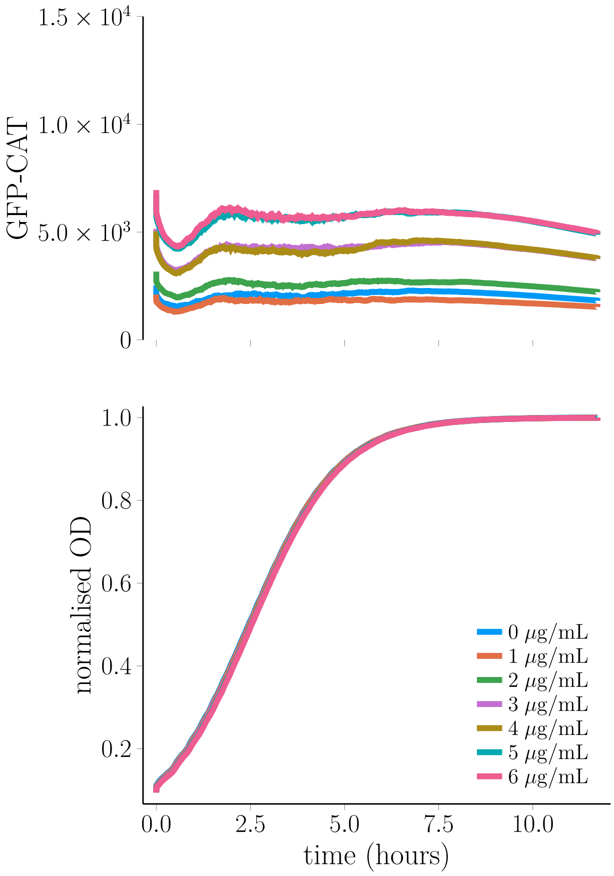

Supplementary Figure 21: Simulating round 2 washout experiment with inducible promoter model. Cell states are taken from the end of simulations presented in Figure 2b of main paper and used as initial conditions for a new simulation without arabinose or chloramphenicol. Upper panel shows GFP-CAT time series simulated over a time period of 12 hours and the lower panel shows corresponding OD levels varying over the same time period.

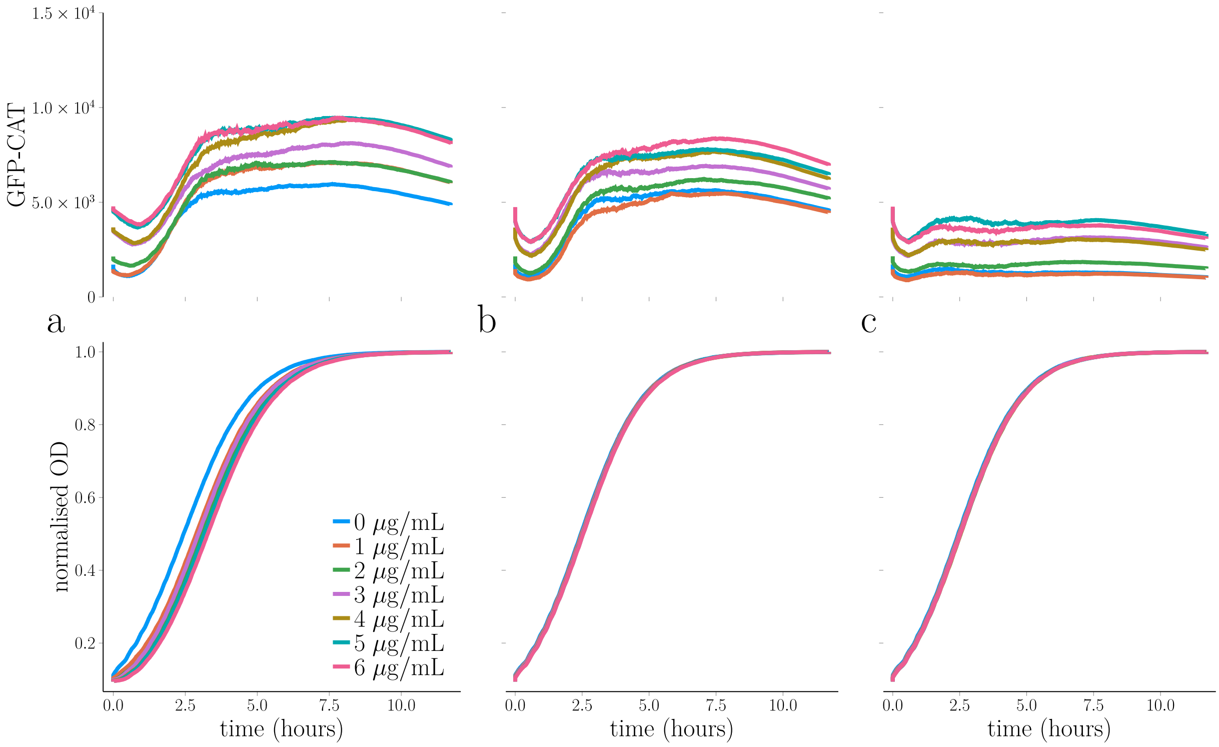

Supplementary Figure 22: Simulating round 3 washout experiment with inducible promoter model. Cell states are taken from the end of simulations presented in Supplementary Figure 21 and used as initial conditions for 3 additional scenarios. Upper panels shows GFP-CAT time series simulated over a time period of 12 hours and lower panels shows corresponding OD levels varying over the same time period. (a) shows the case where arabinose and 7 different chloramphenicol levels are added at the beginning of the simulation. (b) shows the case where only arabinose is added at the beginning of the simulation. (c) shows the case where neither arabinose or chloramphenicol is added.

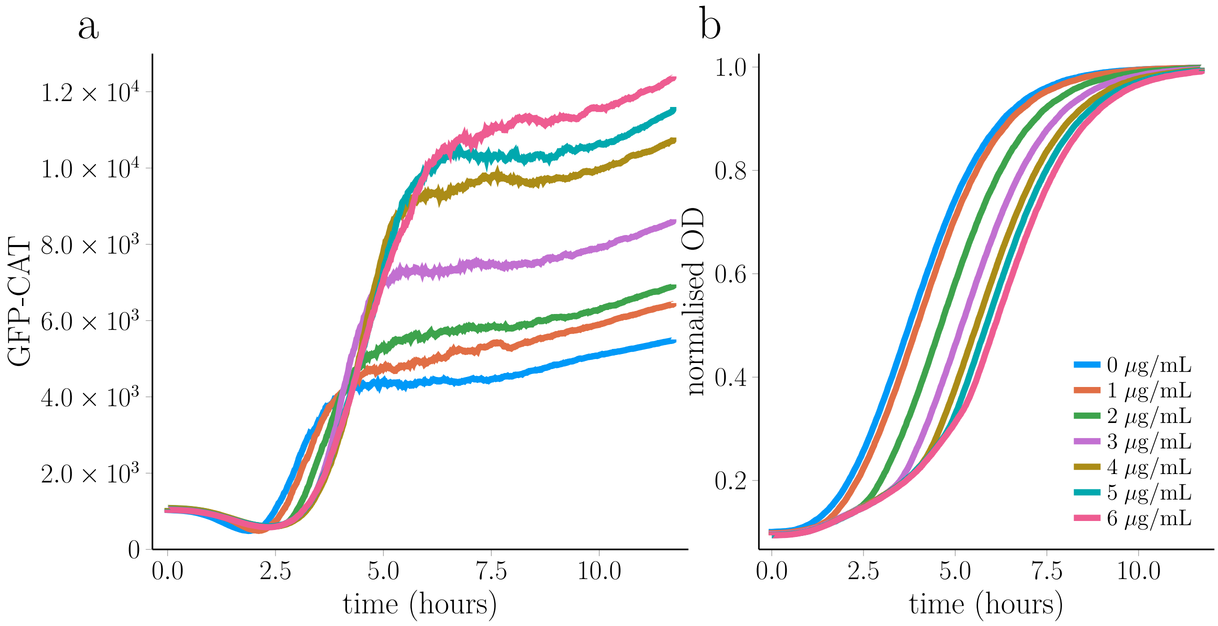

Supplementary Figure 23: Simulations of inducible promoter model defined in section 1.7 with chloraphenicol import function defined in equation (106). Parameters used are obtained from best fit of microplate reader data shown in Figure 2b of main paper and mass spectrometry data shown in Figure 2d. (a) mean GFP-CAT levels varying in time over a period of 12 hours for 7 different doses of chloramphenicol defined in legend. (b) normalised OD levels varying in time over a period of 12 hours for 7 different doses of chloramphenicol.

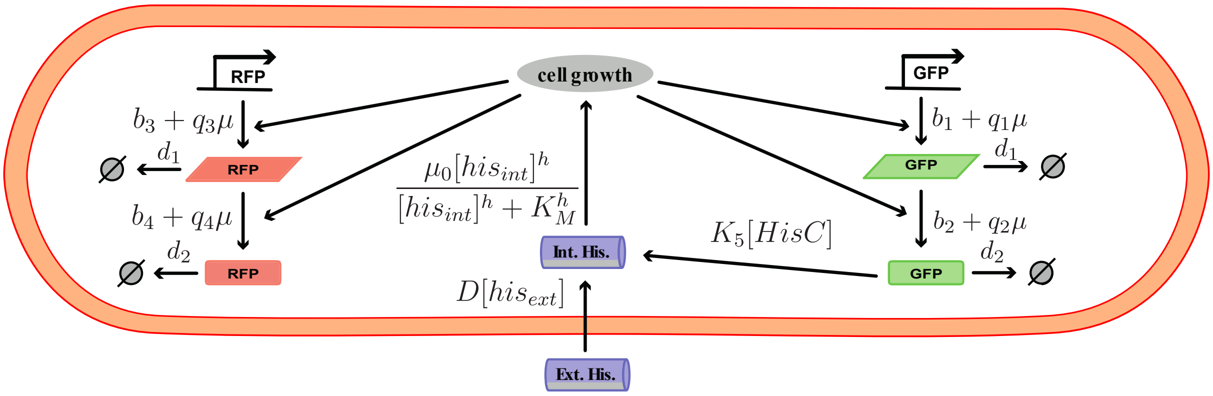

Supplementary Figure 24: Schematic summarising Histidine depletion model. Square brackets denote concentration, i.e., the species is divided by the cell volume, V. For further details on individual reactions, see text. Parallelograms represent mRNA species and rounded rectangles represent protein species.

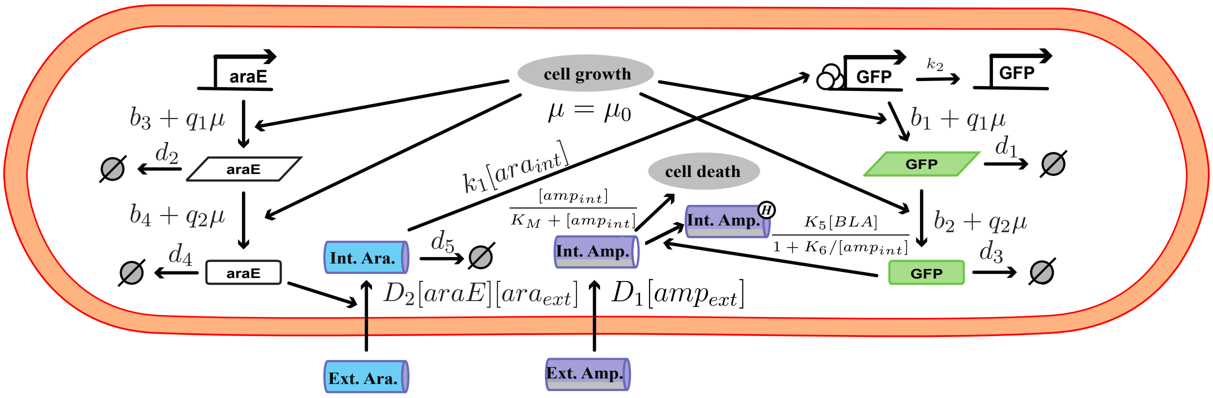

Supplementary Figure 25: Schematic summarising ampicillin model. Square brackets denote concentration, i.e., the species is divided by the cell volume, V. For further details on individual reactions, see text. Parallelograms represent mRNA species and rounded rectangles represent protein species.

### 2 Mathematical modelling supplementary methods and results

#### 2.1 Agent-based simulation of stochastic gene expression and cell growth

Gene expression is noisy due to intrinsic and extrinsic stochasticity [32, 43]. Intrinsic stochasticity arises from the probabilistic timing of the biochemical reactions, resulting in cell-to-cell variation in gene expression levels within clonal cell populations, even if the environment is completely homogeneous [45]. Its effects are most pronounced when the number of biomolecules in the system is small. Extrinsic stochasticity is generated from interactions of the system of interest with other stochastic systems in the cell such as cell cycle or its environment, generating stochastic kinetic parameters, and it is responsible for cell-to-cell variation beyond what is expected from intrinsic variability generated by the reactions within the gene regulatory network.

In order to capture the coupling of stochastic gene expression and cell-to-cell variability in growth rate required for phenotypic selection that underlies emergent gene expression (EGE), we resort to an agent-based modelling approach [30]. In our simulations, agents are single cells that are growing and dividing and inside the cells there are biochemical reactions that take place. A single cell's growth rate is coupled to stochastic expression of a fitness inducing gene inside that cell. In order to simulate our model we use a mixture of the stochastic simulation algorithm [34] to capture gene expression dynamics and analytical solutions of exponential or logistic growth models to capture the cell growth dynamics. The Tau-leaping algorithm [33] is a commonly used approximation of the original stochastic simulation algorithm that allows for longer time steps to be calculated and therefore reduces the computational cost of the simulation. We employ both the Tau-leaping algorithm (when conducting Bayesian parameter estimation) and a stochastic simulation algorithm throughout our study.

As in [41], to manage the computational complexity, we simulate a fixed number of cells ($C$), upon any cell division, the new offspring replaces one of the old cells in the population at random. We simulate the evolution of the state-matrix $M_{i,j}(t)$, a matrix containing the quantities of the molecular species $j$ in cell $i$ at time $t$ in the model. Another matrix, $P_{i,j}(t)$ stores the propensities of the reactions in the system. A third matrix $K_{i,j}(t)$, represents the state-change matrix which stores the changes in the number of the different molecular species at each time step and is used to update the state-matrix $M_{i,j}(t)$. Matrix $K_{i,j}(t)$ is computed using the propensities from $P_{i,j}(t)$ and the tau leaping algorithm.

The approximation of the tau leaping algorithm is formulated around the following consideration: if there is an infinitesimal time $\tau$ such that when time progresses from $t$ to $t+\tau$ there is no significant change in any of the propensity functions, the number of successful reactions of type $j$ within that time $\tau$ can be approximated by a Poisson random variable, $K_{i,j}(t+\tau)$ = $Pois(P_{i,j}(x)\tau)$, i.e., a Poisson number is selected with mean and variance of $P_{i,j}(x)$. We used a constant leap of $\tau = 4seconds$. The time of the state-matrix is then updated to $t+\tau$ as well as its content, using matrix $K_{i,j}(t+\tau)$. With the new state-matrix propensities re-calculated using the new molecular numbers. This is repeated until the volume of the cell reaches its final volume and division occurs. In the event of a negative state variable being detected, we simply repeat the simulation with a smaller $\tau$ or switch to the SSA [33]. We have provided pseudocode in Algorithm 1 for the constitutively expressed model which we define in Section S1.6.

#### 2.2 Cell growth, size control and molecular partitioning

We assumed exponential cell growth using the following deterministic differential equation:

$$\frac{dV_i(t)}{dt} = \mu_i(t)V_i(t) \tag{1}$$

where $V_i(t)$ is the size of cell $i$ at time $t$ and $\mu_i(t)$ is the growth rate of cell $i$ at time $t$. In order to model different experimental setups, we considered two different forms of $\mu_i(t)$. We consider a static environment that enables constant growth, which can be modelled as follows:

$$\mu_i = \mu_0. \tag{2}$$

In this case the growth rate is kept constant throughout the whole experiment, which resembles a chemostat setting. To model the experimental setup of the The microplate reader using our fix cell number agent-based simulations, We used logistic growth, where growth saturates as the population of cells reaches their carrying capacity which can be modelled as follows:

$$\mu_i(t) = \mu_0 N_D(t)\Big(1-\frac{N_D(t)}{k_N}\Big) \tag{3}$$

where $k_N$ is the carrying capacity, which is the maximum number of cell divisions the culture can support, $N_D(t)$ is the number of cell divisions up to time $t$ and $\mu_0$ is the maximal growth constant. In both growth modes, $\mu_0$ could be a function of the level of fitness inducing genes, when that is being modelled. The solution of the constant growth case can be written as

$$V_i(t) = V_i(0)\exp(\mu_0 t) \tag{4}$$

while for the logistic growth case it becomes

$$V_i(t) = V_i(0)\exp\Big(\mu_0 N_D(t)\Big(1-\frac{N_D(t)}{k_N}\Big)t\Big) \tag{5}$$

**Algorithm 1** SSA algorithm for constitutively expressed model

1. Create cell array with $C \times 5$ empty entries, where $C$ corresponds to the number of cells tracked and 5 corresponds to the number of molecular species of interest. Set $t = 0$ and for each cell $i$ set $V_i(0) = 1$, $DNA_i(0) = 1$, $mRNA_i(0) = 10$, $protein_i(0) = 100$, $chl_{int_i}(0) = 0$ and compute $L_F$.

2. Generate two random numbers $\xi_1$ and $\xi_2$ uniformly distributed in (0,1).

3. For each cell $i = 1$ to $C$ evaluate the propensity functions for the following reactions:
   $\alpha_{i,1}(t) = (b_1 + q_1\mu)DNA_i(t)$,
   $\alpha_{i,2}(t) = d_1 mRNA_i(t)$,
   $\alpha_{i,3}(t) = (b_2 + q_2\mu)mRNA_i(t)$,
   $\alpha_{i,4}(t) = D_1 chl_{ext}(t)$,
   $\alpha_{i,5}(t) = \frac{K_5 protein_i(t)}{1+\frac{K_6}{chl_{int_i}(t)}}$,
   $\alpha_{i,6}(t) = d_2 protein_i(t)$,
   and evaluate $\alpha_0 = \sum_{i=1}^{N}\sum_{j=1}^{6} \alpha_{i,j}(t)$.

4. Compute the time when the next reaction takes place as $t + \tau$ where $\tau$ is given by $\tau = \frac{1}{\alpha_0}\ln[\frac{1}{\xi_1}]$.

5. Set $(I,J)$ to be the smallest integers satisfying $\sum_{i=1}^{I}\sum_{j=1}^{J} \alpha_{i,j}(t) > \xi_2\alpha_0$.

6. If $J = 1$, set $mRNA_I(t+\tau) = mRNA_I(t) + 1$.
   If $J = 2$, set $mRNA_I(t+\tau) = mRNA_I(t) - 1$.
   If $J = 3$, set $protein_I(t+\tau) = protein_I(t) + 1$.
   If $J = 4$, set $chl_{int_I}(t+\tau) = chl_{int_I}(t) + 1$.
   If $J = 5$, set $chl_{int_I}(t+\tau) = chl_{int_I}(t) - 1$.
   If $J = 6$, set $protein_I(t+\tau) = protein_I(t) - 1$.
   Update propensities.

7. For $i = 1$ to $N$, set $V_i(t+\tau) = V_i(t)\exp\Big(\mu_0 N_D(t+\tau)\Big(1 - \frac{N_D(t+\tau)}{k_N}\Big)\Big)$. Check if $V_i(t+\tau) \geq L_F$ and then binomially distribute contents of cell $i$ between cells $i$ and a randomly selected cell in the population.

8. Set $t = t + \tau$, if $t >= F$ then end.

To have a realistic cell size distribution, we model size control through a noisy linear map [36], i.e., the final volume $L_F$ of a given cell was assumed to follow

$$L_F = aL_I + b + \eta_1 \tag{6}$$

where $L_I$ is the initial volume of the cell, $a$ and $b$ are linear function parameters and $\eta_1$ is sampled from $\mathcal{N}(0, \sigma_1)$. The dividing cell of volume $L_F$ gives rise to two daughter cells with initial volume $L_I = L_F \times \eta_2$ and $L_I = L_F \times (1 - \eta_2)$ where $\eta_2$ is sampled from $\mathcal{N}(0.5, \sigma_2)$. For all simulations presented, we fix these parameters to $a = 1$, $b = 1\mu m^3$, $\sigma_1 = 0.2\mu m^3$ and $\sigma_2 = 0.05$.

DNA and DNA replication is not explicitly modelled and the new daughter cells each inherit one copy of the genes. Other molecules are binomially partitioned in the daughter cells [35].

#### 2.3 Approximate Bayesian Computation inference

Parameter inference and model selection were carried out using Approximate Bayesian Computation (ABC) embedded in Sequential Monte-Carlo. Our approach is based on that of [40] incorporating the model selection procedure outlined by [46]. Briefly, a set of parameter values associated with a certain model (particle) is evolved through a sequence of distributions until it approximates a sample from the posterior distribution over the joint model and parameter space. At each iteration, each particle is assigned an error value, based on a squared Euclidean distance measure between the experimental data and the output from a simulation using its parameter values. 50% of particles with lowest errors are kept and new particles are generated. These new particles are then concatenated with those from the previous iteration, and this is repeated until reaching a desired level of accuracy. The ABC-SMC and the agent based simulations are implemented in Julia programming language. Codes are available upon request.

#### 2.4 Conditions for observing emergent gene expression

In this section, we investigate the minimal model ingredients necessary to account for the selective upregulation of a fitness inducing protein. Emergent gene expression occurs when cells expressing a fitness inducing protein ($N$) become over-represented in a population compared to a reference protein ($R$) which is expressed in a kinetically identical manner. We use the chemostat growth set up for the investigations of this section. Our minimal model consists of up to 4 variables, the fitness mRNA ($n$), fitness protein ($N$), reference mRNA ($r$) and reference protein ($R$). More specifically, the error function we try to minimise using the ABC approach is defined for model $m$ and parameter set $p$ as

$$error_n(m,p) = \frac{\varepsilon + mean(r)}{\varepsilon + mean(n)}, \tag{7}$$

for models with only an mRNA level, while for models with a protein level, we define it as

$$error_N(m,p) = \frac{\varepsilon + mean(R)}{\varepsilon + mean(N)}, \tag{8}$$

The error functions maximises the average fitness mRNA or protein level while penalising the case where the fitness mRNA or protein are close to zero. To achieve this penalisation of the low copy number case, we set $\varepsilon = 100$. Below we present the 10 models which we considered in the model selection. We run the simulation for 1000 minutes and consider the average mean fitness mRNA or protein level at the final time point to avoid any initial condition effect. We use zero expression as initial conditions.

###### 2.4.1 Model 1 - constitutively expressed mRNA impacts cell growth rate

The first model ($m = 1$) is defined by the following 4 reactions which model transcription and mRNA decay of the fitness and reference gene:

$$\mathrm{DNA^n} \xrightarrow{b_1} \mathrm{DNA^n} + \mathrm{n}_i(t) \tag{9}$$

$$\mathrm{n}_i(t) \xrightarrow{d} \varnothing \tag{10}$$

$$\mathrm{DNA^r} \xrightarrow{b_1} \mathrm{DNA^r} + \mathrm{r}_i(t) \tag{11}$$

$$\mathrm{r}_i(t) \xrightarrow{d} \varnothing \tag{12}$$

Additionally, we assumed that the cells grow at a rate

$$\mu_i(t) = s_0 - s_{stress} + \frac{s_{stress} n(t)}{K + n(t)} \tag{13}$$

where $s_0$ is fixed to give a doubling time of 20 minutes and $s_{stress}$ is chosen to be 75% of $s_0$. At cell division the mRNAs are partitioned into daughter cells binomially with $p = 0.5$. We set $K$ equal to 100 and sample $log_{10}(b_1)$ and $log_{10}(d)$ from prior distributions Uniform(-5,5) and Uniform(-5,-1) respectively.

##### 2.4.2 Model 2 - constitutively expressed mRNA impacts cell growth rate and cell division is biased

The second model ($m = 2$) is defined by the following 4 reactions which model transcription and mRNA decay of the fitness and reference protein:

$$\mathrm{DNA^n} \xrightarrow{b_1} \mathrm{DNA^n} + \mathrm{n}_i(t) \tag{14}$$

$$\mathrm{n}_i(t) \xrightarrow{d} \varnothing \tag{15}$$

$$\mathrm{DNA^r} \xrightarrow{b_1} \mathrm{DNA^r} + \mathrm{r}_i(t) \tag{16}$$

$$\mathrm{r}_i(t) \xrightarrow{d} \varnothing \tag{17}$$

Additionally, we assumed that the cells grow at a rate

$$\mu_i(t) = s_0 - s_{stress} + \frac{s_{stress} n(t)}{K + n(t)} \tag{18}$$

where $s_0$ is fixed to give a doubling time of 20 minutes and $s_{stress}$ is chosen to be 75% of $s_0$. At cell division the mRNAs are partitioned into daughter cells binomially with $p = 0.5 + \xi$. We set $K$ equal to 100 and sample $log_{10}(b_1)$, $log_{10}(d)$ and $\xi$ from prior distributions Uniform(-5,5), Uniform(-5,-1) and Uniform(0,0.5) respectively.

##### 2.4.3 Model 3 - constitutively expressed mRNA impacts cell growth rate with global transcriptional feedback

The third model ($m = 3$) is defined by the following 4 reactions which model transcription and mRNA decay of the fitness and reference protein:

$$\mathrm{DNA^n} \xrightarrow{b_1 + q_1 \mu_i(t)} \mathrm{DNA^n} + \mathrm{n}_i(t) \tag{19}$$

$$\mathrm{n}_i(t) \xrightarrow{d} \varnothing \tag{20}$$

$$\mathrm{DNA^r} \xrightarrow{b_1 + q_1 \mu_i(t)} \mathrm{DNA^r} + \mathrm{r}_i(t) \tag{21}$$

$$\mathrm{r}_i(t) \xrightarrow{d} \varnothing \tag{22}$$

Global feedback is modelled by coupling transcription rate to growth rate. Additionally, we assumed that the cells grow at a rate

$$\mu_i(t) = s_0 - s_{stress} + \frac{s_{stress} n(t)}{K + n(t)} \tag{23}$$

where $s_0$ is fixed to give a doubling time of 20 minutes and $s_{stress}$ is chosen to be 75% of $s_0$. At cell division the mRNAs are partitioned into daughter cells binomially with $p = 0.5$. We set $K$ equal to 100 and sample $log_{10}(b_1)$, $log_{10}(d)$ and $log_{10}(q_1)$ from prior distributions Uniform(-5,5), Uniform(-5,-1) and Uniform(-5,5) respectively.

###### 2.4.4 Model 4 - regulated expression of mRNA impacts cell growth rate

The fourth model ($m = 4$) is defined by the following 8 reactions which model promoter activity, transcription and mRNA decay of the fitness and reference protein:

$$\mathrm{DNA}^{\mathrm{n}}_{\mathrm{off}}(t) \xrightarrow{k_1} \mathrm{DNA}^{\mathrm{n}}_{\mathrm{on}}(t) \tag{24}$$

$$\mathrm{DNA}^{\mathrm{n}}_{\mathrm{on}}(t) \xrightarrow{k_2} \mathrm{DNA}^{\mathrm{n}}_{\mathrm{off}}(t) \tag{25}$$

$$\mathrm{DNA}^{\mathrm{n}}_{\mathrm{on}}(t) \xrightarrow{b_1} \mathrm{DNA}^{\mathrm{n}}_{\mathrm{on}}(t) + \mathrm{n}_i(t) \tag{26}$$

$$\mathrm{n}_i(t) \xrightarrow{d} \varnothing \tag{27}$$

$$\mathrm{DNA}^{\mathrm{r}}_{\mathrm{off}}(t) \xrightarrow{k_1} \mathrm{DNA}^{\mathrm{r}}_{\mathrm{on}}(t) \tag{28}$$

$$\mathrm{DNA}^{\mathrm{r}}_{\mathrm{on}}(t) \xrightarrow{k_2} \mathrm{DNA}^{\mathrm{r}}_{\mathrm{off}}(t) \tag{29}$$

$$\mathrm{DNA}^{\mathrm{r}}_{\mathrm{on}}(t) \xrightarrow{b_1} \mathrm{DNA}^{\mathrm{r}}_{\mathrm{on}}(t) + \mathrm{r}_i(t) \tag{30}$$

$$\mathrm{r}_i(t) \xrightarrow{d} \varnothing \tag{31}$$

Additionally, we assumed that the cells grow at a rate

$$\mu_i(t) = s_0 - s_{stress} + \frac{s_{stress} n(t)}{K + n(t)} \tag{32}$$

where $s_0$ is fixed to give a doubling time of 20 minutes and $s_{stress}$ is chosen to be 75% of $s_0$. At cell division the mRNAs are partitioned into daughter cells binomially with $p = 0.5$. We set $K$ equal to 100 and sample $log_{10}(b_1)$, $log_{10}(d)$, $log_{10}(k_1)$ and $log_{10}(k_2)$ from prior distributions Uniform(-5,5), Uniform(-5,-1), Uniform(-10,0) and Uniform(-10,0) respectively.

##### 2.4.5 Model 5 - constitutively expressed fitness protein impacts cell growth rate

The fifth model ($m = 5$) is defined by the following 6 reactions which model transcription, mRNA decay and translation of the fitness and reference protein:

$$\mathrm{DNA}^{\mathrm{n}} \xrightarrow{b_1} \mathrm{DNA}^{\mathrm{n}} + \mathrm{n}_i(t) \tag{33}$$

$$\mathrm{n}_i(t) \xrightarrow{d} \varnothing \tag{34}$$

$$\mathrm{n}_i(t) \xrightarrow{b_2} \mathrm{n}_i(t) + \mathrm{N}_i(t) \tag{35}$$

$$\mathrm{DNA}^{\mathrm{r}} \xrightarrow{b_1} \mathrm{DNA}^{\mathrm{r}} + \mathrm{r}_i(t) \tag{36}$$

$$\mathrm{r}_i(t) \xrightarrow{d} \varnothing \tag{37}$$

$$\mathrm{r}_i(t) \xrightarrow{b_2} \mathrm{r}_i(t) + \mathrm{R}_i(t) \tag{38}$$

Additionally, we assumed that the cells grow at a rate

$$\mu_i(t) = s_0 - s_{stress} + \frac{s_{stress} N(t)}{K + N(t)} \tag{39}$$

where $s_0$ is fixed to give a doubling time of 20 minutes and $s_{stress}$ is chosen to be 75% of $s_0$. At cell division the mRNAs are partitioned into daughter cells binomially with $p = 0.5$. We sample $log_{10}(b_1)$, $log_{10}(b_2)$ and $log_{10}(d)$ from prior distributions Uniform(-5,5), Uniform(-5,5) and Uniform(-5,-1) respectively.

##### 2.4.6 Model 6 - constitutively expressed fitness protein impacts cell growth rate with global transcriptional feedback

The sixth model ($m = 6$) is defined by the following 6 reactions which model transcription, mRNA decay and translation of the fitness and reference protein:

$$\mathrm{DNA}^{\mathrm{n}} \xrightarrow{b_1 + q_1 \mu_i(t)} \mathrm{DNA}^{\mathrm{n}} + \mathrm{n}_i(t) \tag{40}$$

$$\mathrm{n}_i(t) \xrightarrow{d} \varnothing \tag{41}$$

$$\mathrm{n}_i(t) \xrightarrow{b_2} \mathrm{n}_i(t) + \mathrm{N}_i(t) \tag{42}$$

$$\mathrm{DNA}^{\mathrm{r}} \xrightarrow{b_1 + q_1 \mu_i(t)} \mathrm{DNA}^{\mathrm{r}} + \mathrm{r}_i(t) \tag{43}$$

$$\mathrm{r}_i(t) \xrightarrow{d} \varnothing \tag{44}$$

$$\mathrm{r}_i(t) \xrightarrow{b_2} \mathrm{r}_i(t) + \mathrm{R}_i(t) \tag{45}$$

Global feedback is modelled by coupling transcription rate to growth rate. Additionally, we assumed that the cells grow at a rate

$$\mu_i(t) = s_0 - s_{stress} + \frac{s_{stress} N(t)}{K + N(t)} \tag{46}$$

where $s_0$ is fixed to give a doubling time of 20 minutes and $s_{stress}$ is chosen to be 75% of $s_0$. At cell division the mRNAs are partitioned into daughter cells binomially with $p = 0.5$. We sample $log_{10}(b_1)$, $log_{10}(b_2)$, $log_{10}(q_1)$ and $log_{10}(d)$ from prior distributions Uniform(-5,5), Uniform(-5,5), Uniform(-5,5) and Uniform(-5,-1) respectively.

###### 2.4.7 Model 7 - constitutively expressed fitness protein impacts cell growth rate with global translational feedback

The seventh model ($m = 7$) is defined by the following 6 reactions which model transcription, mRNA decay and translation of the fitness and reference protein:

$$\mathrm{DNA}^{\mathrm{n}} \xrightarrow{b_1} \mathrm{DNA}^{\mathrm{n}} + \mathrm{n}_i(t) \tag{47}$$

$$\mathrm{n}_i(t) \xrightarrow{d} \varnothing \tag{48}$$

$$\mathrm{n}_i(t) \xrightarrow{b_2 + q_2 \mu_i(t)} \mathrm{n}_i(t) + \mathrm{N}_i(t) \tag{49}$$

$$\mathrm{DNA}^{\mathrm{r}} \xrightarrow{b_1} \mathrm{DNA}^{\mathrm{r}} + \mathrm{r}_i(t) \tag{50}$$

$$\mathrm{r}_i(t) \xrightarrow{d} \varnothing \tag{51}$$

$$\mathrm{r}_i(t) \xrightarrow{b_2+q_2\mu_i(t)} \mathrm{r}_i(t) + \mathrm{R}_i(t) \tag{52}$$

Global feedback is modelled by coupling translation rate to growth rate. Additionally, we assumed that the cells grow at a rate

$$\mu_i(t) = s_0 - s_{stress} + \frac{s_{stress}N(t)}{K+N(t)} \tag{53}$$

where $s_0$ is fixed to give a doubling time of 20 minutes and $s_{stress}$ is chosen to be 75% of $s_0$. At cell division the mRNAs are partitioned into daughter cells binomially with $p = 0.5$. We sample $log_{10}(b_1)$, $log_{10}(b_2)$, $log_{10}(q_2)$ and $log_{10}(d)$ from prior distributions Uniform(-5,5), Uniform(-5,5), Uniform(-5,5) and Uniform(-5,-1) respectively.

###### 2.4.8 Model 8 - constitutively expressed fitness protein impacts cell growth rate with global transcriptional and translational feedback

The eighth model ($m = 8$) is defined by the following 6 reactions which model transcription, mRNA decay and translation of the fitness and reference protein:

$$\mathrm{DNA^n} \xrightarrow{b_1+q_1\mu_i(t)} \mathrm{DNA^n} + \mathrm{n}_i(t) \tag{54}$$

$$\mathrm{n}_i(t) \xrightarrow{d} \varnothing \tag{55}$$

$$\mathrm{n}_i(t) \xrightarrow{b_2+q_2\mu_i(t)} \mathrm{n}_i(t) + \mathrm{N}_i(t) \tag{56}$$

$$\mathrm{DNA^r} \xrightarrow{b_1+q_1\mu_i(t)} \mathrm{DNA^r} + \mathrm{r}_i(t) \tag{57}$$

$$\mathrm{r}_i(t) \xrightarrow{d} \varnothing \tag{58}$$

$$\mathrm{r}_i(t) \xrightarrow{b_2+q_2\mu_i(t)} \mathrm{r}_i(t) + \mathrm{R}_i(t) \tag{59}$$

Global feedback is modelled by coupling transcription and translation rate to growth rate. Additionally, we assumed that the cells grow at a rate

$$\mu_i(t) = s_0 - s_{stress} + \frac{s_{stress}N(t)}{K+N(t)} \tag{60}$$

where $s_0$ is fixed to give a doubling time of 20 minutes and $s_{stress}$ is chosen to be 75% of $s_0$. At cell division the mRNAs are partitioned into daughter cells binomially with $p = 0.5$. We sample $log_{10}(b_1)$, $log_{10}(b_2)$, $log_{10}(q_1)$, $log_{10}(q_2)$ and $log_{10}(d)$ from prior distributions Uniform(-5,5), Uniform(-5,5), Uniform(-5,5), Uniform(-5,5) and Uniform(-5,-1) respectively.

##### 2.4.9 Model 9 - constitutively expressed fitness protein impacts cell growth rate and cell division is biased

The ninth model ($m = 9$) is defined by the following 6 reactions which model transcription, mRNA decay and translation of the fitness and reference protein:

$$\mathrm{DNA^n} \xrightarrow{b_1} \mathrm{DNA^n} + \mathrm{n}_i(t) \tag{61}$$

$$\mathrm{n}_i(t) \xrightarrow{d} \varnothing \tag{62}$$

$$\mathrm{n}_i(t) \xrightarrow{b_2} \mathrm{n}_i(t) + \mathrm{N}_i(t) \tag{63}$$

$$\mathrm{DNA^r} \xrightarrow{b_1} \mathrm{DNA^r} + \mathrm{r}_i(t) \tag{64}$$

$$\mathrm{r}_i(t) \xrightarrow{d} \varnothing \tag{65}$$

$$\mathrm{r}_i(t) \xrightarrow{b_2} \mathrm{r}_i(t) + \mathrm{R}_i(t) \tag{66}$$

Additionally, we assumed that the cells grow at a rate

$$\mu_i(t) = s_0 - s_{stress} + \frac{s_{stress} N(t)}{K + N(t)} \tag{67}$$

where $s_0$ is fixed to give a doubling time of 20 minutes and $s_{stress}$ is chosen to be 75% of $s_0$. At cell division the mRNAs are partitioned into daughter cells binomially with $p = 0.5 + \xi$. We sample $log_{10}(b_1)$, $log_{10}(b_2)$, $\xi$ and $log_{10}(d)$ from prior distributions Uniform(-5,5), Uniform(-5,5), Uniform(0,0.5) and Uniform(-5,-1) respectively.

##### 2.4.10 Model 10 - regulated expression of fitness protein impacts cell growth rate

The final model ($m = 10$) is defined by the following 10 reactions which model promoter activity, transcription, mRNA decay and translation of the fitness and reference protein:

$$\mathrm{DNA}^{\mathrm{n}}_{\mathrm{off}}(t) \xrightarrow{k_1} \mathrm{DNA}^{\mathrm{n}}_{\mathrm{on}}(t) \tag{68}$$

$$\mathrm{DNA}^{\mathrm{n}}_{\mathrm{on}}(t) \xrightarrow{k_2} \mathrm{DNA}^{\mathrm{n}}_{\mathrm{off}}(t) \tag{69}$$

$$\mathrm{DNA}^{\mathrm{n}}_{\mathrm{on}} \xrightarrow{b_1} \mathrm{DNA}^{\mathrm{n}}_{\mathrm{on}}(t) + \mathrm{n}_i(t) \tag{70}$$

$$\mathrm{n}_i(t) \xrightarrow{d} \varnothing \tag{71}$$

$$\mathrm{n}_i(t) \xrightarrow{b_2} \mathrm{n}_i(t) + \mathrm{N}_i(t) \tag{72}$$

$$\mathrm{DNA}^{\mathrm{r}}_{\mathrm{off}}(\mathrm{t}) \xrightarrow{k_1} \mathrm{DNA}^{\mathrm{r}}_{\mathrm{on}}(\mathrm{t}) \tag{73}$$

$$\mathrm{DNA}^{\mathrm{r}}_{\mathrm{on}}(\mathrm{t}) \xrightarrow{k_2} \mathrm{DNA}^{\mathrm{r}}_{\mathrm{off}}(\mathrm{t}) \tag{74}$$

$$\mathrm{DNA}^{\mathrm{r}}_{\mathrm{on}} \xrightarrow{b_1} \mathrm{DNA}^{\mathrm{r}}_{\mathrm{on}} + \mathrm{r}_i(t) \tag{75}$$

$$\mathrm{r}_i(t) \xrightarrow{d} \varnothing \tag{76}$$

$$\mathrm{r}_i(t) \xrightarrow{b_2} \mathrm{r}_i(t) + \mathrm{R}_i(t) \tag{77}$$

Additionally, we assumed that the cells grow at a rate

$$\mu_i(t) = s_0 - s_{stress} + \frac{s_{stress} N(t)}{K + N(t)} \tag{78}$$

where $s_0$ is fixed to give a doubling time of 20 minutes and $s_{stress}$ is chosen to be 75% of $s_0$. At cell division the mRNAs are partitioned into daughter cells binomially with $p = 0.5$. We sample $log_{10}(b_1)$, $log_{10}(b_2)$, $log_{10}(d)$, $log_{10}(k_1)$ and $log_{10}(k_2)$ from prior distributions Uniform(-5,5), Uniform(-5,5), Uniform(-5,-1), Uniform(-10,0) and Uniform(-10,0) respectively.

##### 2.4.11 Bayesian model selection result

We performed a Bayesian model selection of the previously listed ten models and found model 7 to be the winning model at producing a bias in the mean fitness protein level relative to the mean reference protein level (c.f. Supplementary Figure 8). Model 7 wins the model selection as it is also one of the simplest and Bayesian model selection naturally penalises complexity. We wanted to further investigate which of the models are capable of producing a selective upregulation of the fitness protein, so we also peformed individual ABC parameter estimation studies for each of the 10 models. These results are summarised in Table 1. We also show some representative simulations from the models that were sucessful, their final posterior distributions, and the effect of varying mRNA noise on the mean bias.

Only those models that had both mRNA and protein species were capable of producing emergent gene expression. We noticed from the parameter sets which yielded EGE that a low transcription rate ($b_1$) appeared in the final posterior distributions which minimised the error function given by equation (8). From this we suspected that the reason why we need mRNA species in the models to produce EGE is to increase noise in gene expression. To probe this further, we took the parameter set from our ABC test which gave the least error and used the fact that noise in a Poisson process varies as an inverse of the mean. We then decreased the transcription rate in order to decrease the mean mRNA level and simultaneously increased the translation rate to keep the mean protein levels constant. We found that models 7 and 8 showed a linear relationship between mRNA noise (as defined by the coefficient of variation squared) and level of bias, with more noise leading to higher levels of EGE. The skewness of a Poisson distribution is defined as the inverse of the square root of the mean, therefore we could also manipulate the skewness of the mRNA distribution by decreasing the transcription rate. By doing this we found a similar relationship between the mRNA skewness of level of EGE (not shown). Furthermore, we found model 10 did not yield a relationship between mRNA noise and level of EGE, but its final posterior distributions seem to suggest that a slow switching between on and off gene states was required for producing a memory effect. We also note that although model 9 failed to produce any significant EGE when maximising the mean ratio of fitness protein to reference protein, it did produce a significant bias when using the median ratio of fitness protein to reference protein as the error.

While we did not explicitly define our error functions in such a way as to yield a linear relationship between the level of stress ($s_{stress}$) and the expression level of the fitness protein, nevertheless, we found such a relationship (see Supplementary Figures 9(a), 10(a) and 11(a)). We confirmed this using linear regression and found that the reference protein either showed less of a response to varying $s_{stress}$ or occasionally showed a slightly negative relationship.

#### 2.5 Mathematical model for constitutively expressed promoter

We defined each cell at each time point $t$ by the number of Gfp-cat mRNAs, GFP-CAT proteins, the volume of the cell and the number of chloramphenicol molecules. When two genes are at opposite sides of the origin of replication and at a similar distance from it, no systematic bias in the copy number is observed [32]. We assumed that the gene copy number for GFP-CAT was constant throughout the simulation and equal to 1 for simplicity. We assumed mRNAs are transcribed from the DNA and mRNAs are in turn translated into protein:

$$\text{DNA} \xrightarrow{b_1+q_1\mu} \text{mRNA}_i(t) \tag{79}$$

$$\text{mRNA}_i(t) \xrightarrow{d_1} \varnothing \tag{80}$$

$$\text{mRNA}_i(t) \xrightarrow{b_2+q_2\mu} \text{mRNA}_i(t) + \text{protein}_i(t) \tag{81}$$

$$\text{protein}_i(t) \xrightarrow{d_2} \varnothing \tag{82}$$

where the subscript $i$ denotes the cell number within the simulation. DNA is transcribed at rate $b_1 + q_1\mu$ which is comprised of a basal transcription rate $b_1$ in addition to a growth dependent one which is consistent with recent experimental evidence [37, 38]. The amount of protein expressed per cell is decreased in the data for the constitutive case from the beginning to the end of experiment. This could imply an initial high level protein that is diluted throughout the experiment. For this we set the initial conditions taken directly from the data and used transcription rates that only depend on the growth rate (as shown above). But we note that alternatively, we could have assumed a decreasing dependence of the transcription rate on OD. This could be due to lower metabolic activitiy of cells at high OD. Specifically, we found that using steady state initial conditions, coupled with a transcription term of the form

$$b_1 + q_1\mu + b_3 \frac{K_{OD}}{K_{OD} + N_D(t)/k_D}, \tag{83}$$

where $k_{OD}$ is an inhibitory threshold constant was able to capture the data equally well (results not shown).

We assumed mRNA is translated at a rate $b_2 + q_2\mu$ which, like the transcription rate, is comprised of a basal translation rate $b_2$ and a growth dependent rate which is also consistent with recent experimental evidence. We assumed mRNA is degraded at a rate $d_1$ and protein is degraded at a rate $d_2$. The equation for growth was modelled following [31] as follows:

$$\mu_i(t) = \frac{\mu_0}{1 + \frac{chl_{int_i}(t)/V_i(t)}{K_M}} N_D(t)\left(1 - \frac{N_D(t)}{k_N}\right) \tag{84}$$

where $\mu_0$ is the maximal growth rate in the absence of chloramphenicol, $chl_{int_i}(t)$ is the internal concentration of chloramphenicol of cell $i$ at time $t$ and $K_M$ is the concentration of chloramphenicol that reduces the growth rate by half, the half-inhibition concentration. We can clearly see from equation (84) that the growth rate is negatively affected by chloramphenicol which indirectly affects both transcription and translation as they contain the growth term $\mu_i(t)$ (equations (79) and (81)) which is consistent with the model of [31]. Essentially there are two positive feedbacks through the growth term, one on transcription and one on translation. Chloramphenicol was modelled as either a constant or depleting external pool with chloramphenicol molecules translocating inside individual cells as in [31]. The influx and acetylation of chloramphenicol were modelled as follows:

$$chl_{ext}(t) \xrightarrow{Chl_{influx}} chl_{int_i}(t) \tag{85}$$

$$chl_{int_i} + protein_i \xrightarrow{Chl_{acetylation}} \varnothing \tag{86}$$

where $chl_{ext}(t)$ represents the external chloramphenicol pool, $chl_{int_i}(t)$ is the internal chloramphenicol in cell $i$ and time $t$, $Chl_{influx}$ is the rate of chloramphenicol influx and $Chl_{acetylation}$ is the rate of chloramphenicol acetylation. The uptake of chloramphenicol into bacteria cells is known not to be passive but instead mediated by protein-coupled oligopetide transporter Ydgr [42], which for simplicity we model as a linear process

$$Chl_{influx} = D_1 chl_{ext}(t) N_D(t)/k_N \tag{87}$$

where $D_1$ is the import rate of chloramphenicol. As a further model simplification, we ignore any export of chloramphenicol. The dependence on $N_D/k_N$ captures the fact that the import of chloramphenicol increases with the size of the cell population. the The acetylation of chloramphenicol was modelled following the Michaelis menten rate [31]:

$$Chl_{acetylation} = \frac{K_5 protein_i(t)/V_i(t)}{1 + \frac{K_6}{chl_{int_i}(t)/V_i(t)}} \tag{88}$$

where $K_5$ is the acetylation strength of GFP-CAT and $K_6$ is the affinity of GFP-CAT acetylation of chloramphenicol. All of these reactions are summarised in a schematic diagram, see Supplementary Figure 12.

##### 2.5.1 Additional data from simulations of constitutive promoter model

In this section, we present time series of Cat mRNA and chloramphenicol levels of the constitutive promoter model defined in section 1.6. These data present corresponding Cat mRNA and chloramphenicol levels for the model simulations presented in Supplementary Data Figure 3. The parameters used represent the best fit parameters obtained from fitting the constitutive promoter model to the microplate reader data. Supplementary Figure 13(a) shows how the mean intracellular chloramphenicol levels vary in time. As in the case of pBAD system, the strongest growth occurs after the chloramphenicol is completely acetylated, with a larger lag in the growth being associated with a larger dose of chloramphenicol. As is observed at the protein level, we also observe an upregulation of GFP-CAT at the mRNA level. This also occurs in a dose dependent manner, with more sustained upregulation of mRNA associated with larger levels of chloramphenicol.

In Supplementary Figure 14 we show the posterior parameter distributions obtained from using an Approximate Bayesian Computation algorithm with microplate reader data presented in Supplementary Data Figure 3. Approximate Bayesian computation (ABC) or likelihood-free methods were developed to deal with models where likelihood calculations fail. Through sampling from unbiased uniform prior distribution we store parameter sets that yield simulated data sufficiently resembling the gene expression and OD microplate reader data, thus generating an approximation for the posterior distribution. In particular, we use an ABC algorithm that specialises in minimising the number of simulations for reaching a given quality of the posterior approximation [40]. For our error function, we used a squared Euclidean distance metric on the GFP-CAT and OD microplate reader time series data (shown in Figures 2A and B) from the main paper. The final posterior distributions are presented in Supplementary Figure 14 as a correlation plot which shows the relationships between each of the 12 model parameters. We highlight that the two strongest relationships we found were a positive relationship between the maximum number of chloramphenicol molecules and the acetylation rate (correlation coefficient of 0.63) and a negative relationship between the growth dependent transcriptional positive feedback and the growth dependent translational positive feedback (correlation coefficient of -0.71). We also found that the growth rate was particularly well constrained by the data. By inverting the covariance matrix of the final probability distribution shown on the diagonal of Supplementary Figure 14 we were able to find the most sensitive model parameters (see Supplementary Figure 15). The protein and mRNA degradation rates are the most sensitive, followed by the cell growth rate. The least sensitive parameter was $K_6$, a parameter which represents the affinity of GFP-CAT acetylation of chloramphenicol.

#### 263 2.6 Mathematical model for inducible promoter

We defined each cell at each time point $t$ by the number of Gfp-cat mRNAs, GFP-CAT proteins, araE mRNAs, araE proteins, the volume of the cell, the number of arabinose molecules and the number of chloramphenicol molecules. We assumed that the activity level of the GFP-CAT pBAD promoter is a function of arabinose or in other words the gene copy number for GFP-CAT varied as a function of arabinose. For the constitutive AraE gene (which we label DNA), we assumed that the gene copy number was constant throughout the simulation and equal to 1 for simplicity. As in the constitutive model we assumed mRNA is transcribed from DNA and then subsequently translated into protein.

$$\varnothing \xrightarrow{k_1[ara_{int}]} \text{pBAD}_i(t) \tag{89}$$

$$\text{pBAD}_i(t) \xrightarrow{k_2} \varnothing \tag{90}$$

$$\text{pBAD}_i(t) \xrightarrow{b_1+q_1\mu} \text{Cat-mRNA}_i(t) \tag{91}$$

$$\text{Cat-mRNA}_i(t) \xrightarrow{d_1} \varnothing \tag{92}$$

$$\text{Cat-mRNA}_i(t) \xrightarrow{b_2+q_2\mu} \text{Cat-mRNA}_i(t) + \text{Cat-protein}_i(t) \tag{93}$$

$$\text{CAT-protein}_i(t) \xrightarrow{d_3} \varnothing \tag{94}$$

$$\text{DNA} \xrightarrow{b_3+q_1\mu} \text{AraE-mRNA}_i(t) \tag{95}$$

$$\text{AraE-mRNA}_i(t) \xrightarrow{d_2} \varnothing \tag{96}$$

$$\text{AraE-mRNA}_i(t) \xrightarrow{b_4+q_2\mu} \text{AraE-mRNA}_i(t) + \text{AraE-protein}_i(t) \tag{97}$$

$$\text{AraE-protein}_i(t) \xrightarrow{d_4} \varnothing \tag{98}$$

where squarebrackets denote concentration (copy number divided by volume) and the subscript $i$ denotes the cell number within the simulation. The pBAD promoter is activated at a rate $k_1[ara_{int}]$ and becomes inactive at a rate $k_2$. Cat mRNA is transcribed at rate $b_1 + q_1\mu$ which is comprised of a basal transcription rate $b_1$ in addition to a growth rate dependent one. mRNA is translated at a rate $b_2 + q_2\mu$ which, like the transcription rate, is comprised of a basal translation rate $b_2$ and a growth dependent rate. We assumed Cat mRNA is degraded at a rate $d_1$ and CAT proteins degrade at a rate $d_2$. As in the constitutive case the equation for growth was modelled following [31] as follows:

$$\mu_i(t) = \frac{\mu_0}{1 + \frac{chl_{int_i}(t)/V_i(t)}{K_M}} N_D(t)\left(1 - \frac{N_D(t)}{k_N}\right) \tag{99}$$

where $\mu_0$ is the maximal growth rate in the absence of chloramphenicol, $chl_{int_i}(t)$ is the internal concentration of chloramphenicol of cell $i$ at time $t$ and $K_M$ is the concentration of chloramphenicol that reduces the growth rate by half, the half-inhibition concentration. We can clearly see from equation (99) that the growth rate is negatively affected by chloramphenicol which indirectly affects both transcription and translation as they contain the growth term $\mu_i(t)$ (equations (79) and (81)) which is consistent with the model of [31]. Essentially there are two positive feedbacks through the growth term, one on transcription and one on translation. Chloramphenicol and arabinose were modelled as depleting external pools. The import and acetylation of chloramphenicol were modelled as follows:

$$chl_{ext}(t) \xrightarrow{Chl_{influx}} chl_{int_i}(t) \tag{100}$$

$$chl_{int_i} + protein_i \xrightarrow{Chl_{acetylation}} \varnothing \tag{101}$$

where $chl_{ext}(t)$ represents the external chloramphenicol pool, $chl_{int_i}(t)$ is the internal chloramphenicol in cell $i$ and time $t$, $Chl_{influx}$ is the rate of chloramphenicol influx and $Chl_{acetylation}$ is the rate of chloramphenicol acetylation. The influx of chloramphenicol was modelled as in the constitutive promoter case:

$$Chl_{influx} = D_1 chl_{ext}(t) N_D(t)/k_N \tag{102}$$

where $D_1$ is the import rate of chloramphenicol and we assumed chloramphenicol is not exported. We again assumed that the magnitude of the influx of chloramphenicol depends on the size of the cell population. The acetylation of chloramphenicol was modelled following the Michaelis menten rate [31]:

$$Chl_{acetylation} = \frac{K_5 protein_i(t)/V_i(t)}{1 + \frac{K_6}{chl_{int_i}(t)/V_i(t)}} \tag{103}$$

where $K_5$ is the acetylation strength of GFP-CAT and $K_6$ is the affinity of GFP-CAT acetylation of chloramphenicol. Finally, the import of arabinose was modelled by the following reaction:

$$ara_{ext}(t) \xrightarrow{Ara_{influx}} ara_{int_i}(t) \tag{104}$$

where

$$Ara_{influx} = D_2 ara_{ext}(t)\text{AraE-protein}_i(t) N_D(t)/k_N. \tag{105}$$

This second order reaction captures the co-dependence of the import of arabinose on the size of the arabinose pool and the amount of the arabinose transporter protein in the cell. The rate at which this reaction occurs is represented by the parameter $D_2$. All of these reactions are summarised in a schematic diagram, see Supplementary Figure 16. We also present a model schematic where the global growth dependent transcriptional and translational positive feedback loops are omitted, see Supplementary Figure 17.

##### 2.6.1 Additional results from simulations of inducible promoter model

In this section, we present time series data of pBAD promoter activity level and mRNA levels from the inducible promoter model defined in section 1.7. These data present corresponding pBAD promoter activity levels and chloramphenicol levels for the model simulations presented in Figure 2b of the main paper. The parameters used represent the best fit parameters obtained from fitting the inducible promoter model to the microplate reader data. Supplementary Figure 18(a) shows how the mean pBAD promoter activity levels vary in time. The pBAD activity level increases in a chloramphenicol dose dependent manner. It also diminishes slowly on a time-scale related to the unbinding rate of arabinose from the pBAD promoter ($k_2$). Supplementary Figure 18(b) shows how the mean Cat mRNA level varies in time. As is observed at the protein level, we also observe an upregulation of Gfp-cat at the mRNA level. This also occurs in a dose dependent manner, with more sustained upregulation of mRNA associated with larger levels of chloramphenicol. Consistent with the data presented in Figure 2f of the main paper, we can observe that mRNA levels are elevated for all time points and for all non-zero chloramphenicol doses.

In order to probe the relationship between the constitutive and inducible models, we performed some control simulations with the inducible promoter model. From our modelling study presented in section 1.4, we know that the constitutive model needed links between cell growth and gene expression to produce emergent gene expression. However, since the inducible promoter model is more complex, it may not need these links to yield the microplate reader data. In Supplementary Figure 19(a) we show the result of removing these links (setting parameters $q_1$ and $q_2$ to zero, see Supplementary Figure 17) from the inducible promoter model where the other parameters are set to the best fit parameters. Clearly we see that the GFP-CAT simulated data no longer agrees with the data and without these links between gene expression and cell growth, the upregulation is no longer observed. In addition to this numerical experiment, we also performed a Bayesian model selection between the models with and without these gene expression-cell growth links (see schematics presented in Supplementary Figures 16 and 17) to see which model was the best at reproducing the microplate reader data presented in Figure 2b. This resulted in the model with these gene expression-cell growth links being chosen with 100% probability. In Supplementary Figure 19(b) we show the effect of fixing the pBAD activity level to a constant, 1, which reduces the inducible promoter model to the constitutive promoter model. We keep the parameters used fixed to those which were the best fit for the microplate reader data presented in Figure 2b of the main paper. This results in a transient emergent gene expression behaviour.

We also studied the posterior parameter distributions obtained from using an Approximate Bayesian Computation algorithm with microplate reader data presented in Figure 2b of the main paper. We used the same ABC algorithm we did for the constitutive promoter case and for our error function, we used a squared Euclidean distance metric on the GFP-CAT and OD microplate reader time series data. We performed a correlation analysis of the final posterior distributions which can uncover relationships between each of the 21 model parameters. The strongest relationships we found were a positive one between the maximum number of arabinose molecules and the acetylation rate (correlation coefficient 0.52) and as we found in the constitutive case, a negative relationship between the growth dependent transcriptional positive feedback and the growth dependent translational positive feedback (correlation coefficient of -0.62). Furthermore, as in the constitutive case, we also found that the growth rate was particularly well constrained by the data. By inverting the covariance matrix of the final probability distribution we were able to find the most sensitive model parameters (see Supplementary Figure 20). The pBAD promoter deactivation rate ($k_2$) was the most sensitive parameter, followed by the degradation rate of CAT protein ($d_2$) and the acetylation rate ($K_5$). The least sensitive parameter was $D_1$ which represents the import rate of chloramphenicol molecules.

##### 2.6.2 Reversibility of emergent gene expression for inducible promoter system

In this section we reproduce the washout experiments shown in Supplementary Data Figure 2 using simulations of the inducible promoter model. The purpose of these experiments was to show the reversibility of the emergent gene expression phenomenon for the inducible promoter case. While in the constitutive case, we can observe this reversibility as soon as the chloramphenicol is acetylated, in the case of the inducible promoter it depends on how quickly the pBAD promoter deactivates. To reproduce these experiments computationally, we use the cell states (i.e. number of mRNA, protein, size of cells, pBAD activity level) at the final time step, and use this as the initial condition for subsequent computational experiments. Though we did not fit the washout experiments explicitly using the model, we find that the model reproduces the observed data qualitatively. In Supplementary Figure 21 we see that the GFP-CAT levels diminish but still maintain the chloramphenicol dose dependent ordering from the initial round of arabinose and chloramphenicol. Since there is no additional chloramphenicol, we see that the cells all grow at the same rate and produce similar OD curves (lower panel of Supplementary Figure 21).

We simulated an additional round of washout experiments and present the results in Supplementary Figure 22. This time we took the cell states from the final time point of the simulation presented in Supplementary Figure 21 and used these states as the initial conditions for three different scenarios. In the first simulation, we add arabinose and the same 7 different dosages of chloramphenicol as in the initial experiment. We find that while the GFP-CAT levels rise according to the level of chloramphenicol, they do not rise as much as in the initial experiment. This is due to the initial condition of GFP-CAT allowing for acetylation of chloramphenicol immediately, therefore negating the need for GFP-CAT levels to rise as much as they did in the initial experiment. As a consequence of this, we only observe a small impact on the corresponding OD curves. In the second simulation, we add arabinose but don't add chloramphenicol. This causes the GFP-CAT levels to rise, but not as much they do in the case of chloramphenicol and arabinose addition. The growth rate is unaffected in this scenario and all the OD curves have a similar appearance. In the third simulation, we once again simulate the scenario we did in round 2, where neither arabinose or chloramphenicol are added. This causes the GFP-CAT levels to decrease further, but we note that in the simulation the observed GFP-CAT levels do not decrease as much as they did in the experiments presented in Supplementary Data Figure 2. This may be due to small delays introduced into restarting the next round of experiments allowing mRNA species to decay which are not simulated in our computational simulations.

##### 2.6.3 Capturing mass spectrometry data accurately

Fitting the inducible promoter model to the data presented in Figure 2b of the main paper achieved good agreement with our experimental data in terms of gene expression and OD. However, we found that without explicitly accounting for mass spectrometry data in our error function which we used for parameter inference, the resulting parameter sets predicted that the external chloramphenicol pool depleted too rapidly from simulations of our model compared with the data we obtained from mass spectrometry experiments. To remedy this, we modified our model so that the chloramphenicol import function from having a linear dependence on the external chloramphenicol pool to one that saturates was necessary to capture the external chloramphenicol pool depletion rate accurately. Specifically, we changed the influx rate from

$$Chl_{influx} = D_1 chl_{ext}(t) N_D(t)/k_N \tag{106}$$

to

$$Chl_{influx}* = D_1 \frac{chl_{ext}(t)}{K_{chl} + chl_{ext}(t)} N_D(t)/k_N \tag{107}$$

where $K_{chl}$ is a half saturation constant. After making this modification, we repeated our ABC parameter inference and found good agreement with gene expression data, OD data and mass spectrometry data. This model produced a reasonable fit to the mass spectrometry data (Figure 2f). The other model predictions such as GFP levels and OD levels remain consistent with the previous fitting (Supplementary Figure 23).

#### 2.7 Mathematical model for Histidine depletion system

We defined each cell at each time point $t$ by the number of Gfp-hisC mRNAs, GFP-HisC proteins, rfp mRNAS, RFP proteins, the volume of the cell and the number of histidine molecules. As in the other models presented, we assumed that the gene copy number for each gene was constant throughout the simulation and equal to 1 for simplicity. We assumed GFP-hisC and rfp mRNAs are transcribed from the DNA as per the following reactions

$$\text{DNA}_{\text{gfp}} \xrightarrow{b_1+q_1\mu} \text{GFP-hisC mRNA}_i(t) \tag{108}$$

$$\text{DNA}_{\text{rfp}} \xrightarrow{b_3+q_3\mu} \text{RFP mRNA}_i(t). \tag{109}$$

These mRNA molecules are then translated into proteins and both mRNA and proteins are assumed to degrade via the following reactions

$$\text{GFP-hisC mRNA}_i(t) \xrightarrow{b_2+q_2\mu} \text{GFP-hisC mRNA}_i(t) + \text{GFP-HisC protein}_i(t) \tag{110}$$

$$\text{RFP mRNA}_i(t) \xrightarrow{b_4+q_4\mu} \text{RFP mRNA}_i(t) + \text{RFP protein}_i(t) \tag{111}$$

$$\text{GFP-hisC mRNA}_i(t) \xrightarrow{d_1} \varnothing \tag{112}$$

$$\text{RFP mRNA}_i(t) \xrightarrow{d_3} \varnothing \tag{113}$$

$$\text{GFP-HisC protein}_i(t) \xrightarrow{d_2} \varnothing \tag{114}$$

$$\text{RFP protein}_i(t) \xrightarrow{d_4} \varnothing \tag{115}$$

where the subscript $i$ denotes the cell number within the simulation. We assumed both the fitness protein, HisC and the reference protein, RFP obey the same equation but with different rates. For example, we assumed HisC mRNA is translated at a rate $b_2 + q_2\mu$ while RFP mRNA is translated at a rate $b_4 + q_4\mu$. For simplicity we assumed both mRNA and protein are degraded at the same rates for both fitness and reference species. In order to simulate Histidine depletion experiments, we take advantage of that fact that the growth rate of bacteria cells is known to depend on the intracellular Histidine levels. Hence, we let the growth rate take the following form

$$\mu(t) = \mu_0 \frac{([His_{int_i}(t)]/V_i(t))^h}{([His_{int_i}(t)]/V_i(t))^h + K_M^h} N_D(t)\Big(1 - \frac{N_D(t)}{k_N}\Big) \tag{116}$$

where $\mu_0$ is the maximal growth rate which can be found when cells are saturated with Histidine, $his_{int_i}(t)$ is the internal concentration of histidine of cell $i$ at time $t$ and $K_M$ is the concentration of Histidine that represents half-maximal growth rate, the half-saturation concentration. The import and production of Histidine were modelled as follows:

$$His_{ext}(t) \xrightarrow{His_{influx}} His_{int_i}(t) \tag{117}$$

$$\text{GFP-HisC protein}_i(t) \xrightarrow{His_{production}} \varnothing \tag{118}$$

where $His_{ext}(t)$ represents the external Histidine pool, $His_{int_i}(t)$ is the internal Histidine in cell $i$ and time $t$, $His_{influx}$ is the rate of Histidine influx and $His_{production}$ is the rate of Histidine production by HisC. The uptake of Histidine into E. Coli cells is known not to be passive but instead active [39], which for simplicity we model as a linear process

$$His_{influx} = DHis_{ext}(t)N_D(t)/k_N \tag{119}$$

where $D$ is the import rate of Histidine. As a further model simplification, we ignore any export of Histidine. The dependence on $N_D/k_N$ captures the fact that the import of Histidine increases with the size of the cell population. The rate of HisC mediated production of intracellular Histidine was modelled by the following simple reaction

$$His_{production} = K_5 \text{GFP-HisC protein}_i(t) \tag{120}$$

where $K_5$ is the production rate of intracellular Histidine. All of these reactions are summarised in a schematic diagram, see Supplementary Figure 24.

#### 2.8 Mathematical model for Ampicillin system

We defined each cell at each time point $t$ by the number of Gfp-bla mRNAs, GFP-BLA proteins, araE mRNAs, araE proteins, the volume of the cell, the number of arabinose molecules and the number of chloramphenicol molecules. We assumed that the activity level of the GFP-BLA pBAD promoter is a function of arabinose or in other words the gene copy number for GFP-BLA varied as a function of arabinose. For the constitutive AraE gene (which we label DNA), we assumed that the gene copy number was constant throughout the simulation and equal to 1 for simplicity. As in the constitutive model we assumed mRNA is transcribed from DNA and then subsequently translated into protein.

$$\varnothing \xrightarrow{k_1[ara_{int}]} \text{pBAD}_i(t) \tag{121}$$

$$\text{pBAD}_i(t) \xrightarrow{k_2} \varnothing \tag{122}$$

$$\text{pBAD}_i(t) \xrightarrow{b_1+q_1\mu} \text{Bla-mRNA}_i(t) \tag{123}$$

$$\text{Bla-mRNA}_i(t) \xrightarrow{d_1} \varnothing \tag{124}$$

$$\text{Bla-mRNA}_i(t) \xrightarrow{b_2+q_2\mu} \text{Bla-mRNA}_i(t) + \text{Bla-protein}_i(t) \tag{125}$$

$$\text{BLA-protein}_i(t) \xrightarrow{d_3} \varnothing \tag{126}$$

$$\text{DNA} \xrightarrow{b_3+q_1\mu} \text{AraE-mRNA}_i(t) \tag{127}$$

$$\text{AraE-mRNA}_i(t) \xrightarrow{d_2} \varnothing \tag{128}$$

$$\text{AraE-mRNA}_i(t) \xrightarrow{b_4+q_2\mu} \text{AraE-mRNA}_i(t) + \text{AraE-protein}_i(t) \tag{129}$$

$$\text{AraE-protein}_i(t) \xrightarrow{d_4} \varnothing \tag{130}$$

where squarebrackets denote concentration (copy number divided by volume) and the subscript $i$ denotes the cell number within the simulation. The pBAD promoter is activated at a rate $k_1[ara_{int}]$ and becomes inactive at a rate $k_2$. Bla mRNA is transcribed at rate $b_1 + q_1\mu$ which is comprised of a basal transcription rate $b_1$ in addition to a growth rate dependent one. mRNA is translated at a rate $b_2 + q_2\mu$ which, like the transcription rate, is comprised of a basal translation rate $b_2$ and a growth dependent rate. We assumed Bla mRNA is degraded at a rate $d_1$ and BLA proteins degrade at a rate $d_2$. The growth rate

$$\mu_i(t) = \mu_0 N_D(t)\Big(1 - \frac{N_D(t)}{k_N}\Big) \tag{131}$$

where $\mu_0$ is the maximal growth rate. We note that from equation (131) as the growth rate slows so too do both transcription and translation as they contain the growth term $\mu_i(t)$ (equations (123) and (125)). As in other models, there are two positive feedbacks through the growth term, one on transcription and one on translation. Ampicillin and arabinose were modelled as depleting external pools. The import and hydrolysis of ampicillin were modelled as follows:

$$amp_{ext}(t) \xrightarrow{Amp_{influx}} amp_{int_i}(t) \tag{132}$$

$$amp_{int_i} + protein_i \xrightarrow{Amp_{hydrolysis}} \varnothing \tag{133}$$

where $amp_{ext}(t)$ represents the external ampicillin pool, $amp_{int_i}(t)$ is the internal ampicillin in cell $i$ and time $t$, $Amp_{influx}$ is the rate of ampicillin influx and $Amp_{hydrolysis}$ is the rate of ampicillin hydrolysis. The influx of ampicillin was modelled as follows

$$Amp_{influx} = D_1 amp_{ext}(t) N_D(t)/k_N \tag{134}$$

where $D_1$ is the import rate of ampicillin and we assumed ampicillin is not exported. We again assumed that the magnitude of the influx of ampicillin depends on the size of the cell population. The cleavage of ampicillin was modelled following the Michaelis menten rate:

$$Amp_{cleavage} = \frac{K_5 protein_i(t)/V_i(t)}{1 + \frac{K_6}{amp_{int_i}(t)/V_i(t)}} \tag{135}$$

where $K_5$ is the cleavage rate of GFP-BLA and $K_6$ is the affinity of GFP-BLA cleavage of ampicillin. Finally, the import of arabinose was modelled by the following reaction:

$$ara_{ext}(t) \xrightarrow{Ara_{influx}} ara_{int_i}(t) \tag{136}$$

where

$$Ara_{influx} = D_2 ara_{ext}(t)\text{AraE-protein}_i(t) N_D(t)/k_N. \tag{137}$$

This second order reaction captures the co-dependence of the import of arabinose on the size of the arabinose pool and the amount of the arabinose transporter protein in the cell. The rate at which this reaction occurs is represented by the parameter $D_2$. Finally, uniquely in this model in contrast to the prior models considered, the cells are assumed to undergo lysis in an ampicillin dependent manner according to the following reaction

$$cell_i \xrightarrow{Amp_{lysis}} \varnothing \tag{138}$$

where $Amp_{lysis}$ is given by

$$Amp_{lysis} = \frac{[amp_{int}]}{[amp_{int}] + K_M}, \tag{139}$$

where $K_M$ is the dose of ampicillin at which the cell lysis rate is half maximal. After a cell death reaction is triggered we remove one cell from the constant population of tracked cells, $C$ and we also update the number of divisions at time point t by $N_D(t) = N_D(t) - N_D(t)/C$. Hence if all tracked cells are removed in one timestep then our proxy for the size of the cell population, $N_D(t)$ also becomes zero. Upon cell division, we preferentially place new daughter cells into spaces occupied by dead cells (in order to keep the number of cells tracked as close to constant as possible). If there are no dead cells in the population at time point $t$ then instead we revert to randomly replacing a cell in the population with a daughter cell. All of these reactions are summarised in a schematic diagram, see Supplementary Figure 25.

### 3 Sequence data

Supplementary Table 2: Genomic integration primer sequences

| primer name | primer sequence |
|---|---|
| intC-Red-F | ATAGTTGTTAAGGTCGCTCACTCCACCTTCTCATCAAGCCAGTCCGCCCAAGAGC AGGAGATTACGACGATC |
| intC-Red-R | CCGTAGATTTACAGTTCGTCATGGTTCGCTTCAGATCGTTGACAGCCGCAGTATC CGCTCATGAGACAATAACCC |
| fim-Red-F | TATTGTCTTATTCTGTTGGCATATCGGCATGGGATGCGTATTAGTGAACCTCTTC AAATGTAGCACCTG |
| fim-Red-R | GCTAACGTGCAGGTTTTTAGCTTCAGGTAATATTGCGTACCAGCATTAGCGGAG ACCATTCCAACCCTTC |
| flu-Red-F | CTCCGGCACTGTAACCCTTTACCTGCCGGTATCCACGTTTGTGGGTACCGCTCTT CAAATGTAGCACCTG |
| flu-Red-R | AGGCGATGGTTCTGTCAGAAGGTCACATTCAGTGTGGCCTGACCGTTATACCTC TGGTAAGGTTGGGAAGC |

Supplementary Table 3: qPCR primer sequences

| primer name | sequence |
|---|---|
| idnT F | ATCCTCATCTGTTTAGCGAAGAGGAGATGC |
| idnT R | AATGATATCCATGATTTGCTCGATGGTGCG |
| hcaT F | AATGATATCCATGATTTGCTCGATGGTGCG |
| hcaT R | TATTACTCAGCGCAAAGATAATGACTTCCGC |
| cysG F | TATTACTCAGCGCAAAGATAATGACTTCCGC |
| cysG R | TAAGCGTACGCTCTGGGCATAATCGCGATG |
| gfp F | GGAAGGCTATGTGCAGGAACGTACGATTAG |
| gfp R | ATCACCAATCGGCGTGTTTTGCTGATAGTG |
| rpoD F | TGAAGACGAAGAAGATGGCGATGACGACAG |
| rpoD R | TTGCACTGCTCAACGCAGAGCTTCATGATC |
| rpoH F | TCTGGAAGCAGCTAAAACGCTGATCCTGTC |
| rpoH R | AGACGCTGCTTGGTTTTACGCAGGTTGAAG |
| rpoE F | GTTGAACGGGTCCAGAAGGGAGATCAGAAAG |
| rpoE R | GCCGCCACTTTCGAAGTTTTCAGCTTCAATGG |
| rpoN F | ATTACCTGATGTGGCAGGTTGAGCTGACAC |
| rpoN R | ATCGCTAATGATCAGTCTGGCCTCTTCCAG |
| acrB F | CAAAGTTGAAGCGATTACCATGCGTGCAAC |
| acrB R | AGAGTGGTGTTAATGTCGTTGATAGAAACAC |
| pntB F | GCGATGAACCGTTCCTTTATCAGCGTTATTG |
| pntB R | TCAGCCAGCAATACGTTCATATGTCCAGGC |
| oppA F | GCAGATCGGAGGTGTTATACAACAGGTTG |
| oppA R | GCAGATCGGAGGTGTTATACAACAGGTTG |
| cyoC F | TGGATCTACCTGATGAGCGACTGCATTCTG |
| cyoC R | CCCATGCCGTTAACAATCAGGTGATGGAATTC |

Supplementary Figure 26: GFP-CAT expression cassettes

Inducible expression: *araC*-pBad-B0034-*gfp*-G4S-*cat*-lox-*kan*-lox cassette

Legend

AraC gene
pBad promoter
B0034 ribosome binding site
*gfp* gene
Gycine(4)-Serine linker
*cat* gene
lox recombination sites
kanamycin resistance gene

GCTCCTAGGTCTGATTCGTTACCAATTATGACAACTTGACGGCTACATCATTCACTTTTTCTTCACAACCG
GCACGGAACTCGCTCGGGCTGGCCCCGGTGCATTTTTTAAATACCCGCGAGAAATAGAGTTGATCGTCA
AAACCAACATTGCGACCGACGGTGGCGATAGGCATCCGGGTGGTGCTCAAAAGCAGCTTCGCCTGGCT
GATACGTTGGTCCTCGCGCCAGCTTAAGACGCTAATCCCTAACTGCTGGCGGAAAAGATGTGACAGACG
CGACGGCGACAAGCAAACATGCTGTGCGACGCTGGCGATATCAAAATTGCTGTCTGCCAGGTGATCGCT
GATGTACTGACAAGCCTCGCGTACCCGATTATCCATCGGTGGATGGAGCGACTCGTTAATCGCTTCCATG
CGCCGCAGTAACAATTGCTCAAGCAGATTTATCGCCAGCAGCTCCGAATAGCGCCCTTCCCCTTGCCCGG
CGTTAATGATTTGCCCAAACAGGTCGCTGAAATGCGGCTGGTGCGCTTCATCCGGGCGAAAGAACCCCG
TATTGGCAAATATTGACGGCCAGTTAAGCCATTCATGCCAGTAGGCGCGCGGACGAAAGTAAACCCACT
GGTGATACCATTCGCGAGCCTCCGGATGACGACCGTAGTGATGAATCTCTCCTGGCGGGAACAGCAAAA
TATCACCCGGTCGGCAAACAAATTCTCGTCCCTGATTTTTCACCACCCCCTGACCGCGAATGGTGAGATT
GAGAATATAACCTTTCATTCCCAGCGGTCGGTCGATAAAAAAATCGAGATAACCGTTGGCCTCAATCGG
CGTTAAACCCGCCACCAGATGGGCATTAAACGAGTATCCCGGCAGCAGGGGATCATTTTGCGCTTCAGC
CATACTTTTCATACTCCCGCCATTCAGAGGAAGAAACCAATTGTCCATATTGCATCAGACATTGCCGTCAC
TGCGTCTTTTACTGGCTCTTCTCGCTAACCAAACCGGTAACCCCGCTTATTAAAAGCATTCTGTAACAAAG
CGGGACCAAAGCCATGACAAAAACGCGTAACAAAAGTGTCTATAATCACGGCAGAAAAGTCCACATTG
ATTATTTGCACGGCGTCACACTTTGCTATGCCATAGCATTTTTATCCATAAGATTAGCGGTTCCTACCTGA
CGCTTTTTATCGCAACTCTCTACTGTTTCTCCGGGTCCCTATCAGTGATAGAGAGAGCTCGTTGAGAAAG
AGGAGAAATACTAGATGCGTAAAGGCGAAGAACTGTTTACCGGTGTGGTTCCGATTCTGGTGGAACTG
GACGGCGATGTTAATGGTCATAAATTCAGTGTTCGCGGCGAAGGTGAAGGCGATGCGACGAACGGCAA
ACTGACCCTGAAATTTATCTGCACCACGGGTAAACTGCCGGTCCCGTGGCCGACGCTGGTGACCACGCT
GACCTATGGCGTTCAATGTTTTGCGCGTTACCCGGATCACATGAAACAGCACGACTTTTTCAAATCGGCC
ATGCCGGAAGGCTATGTGCAGGAACGTACGATTAGCTTTAAAGACGATGGTACGTATAAAACCCGCGC
GGAAGTGAAATTCGAAGGCGATACCCTGGTTAACCGTATCGAACTGAAAGGTATCGATTTCAAAGAAGA
CGGCAATATTCTGGGTCATAAACTGGAATATAACTTCAATTCCCACAACGTGTACATCACCGCGGATAAA
CAGAAAAACGGCATTAAAGCCAATTTCAAAATCCGCCATAATGTGGAAGATGGTAGCGTTCAGCTGGCC
GACCACTATCAGCAAAACACGCCGATTGGTGATGGCCCGGTCCTGCTGCCGGACAATCACTACCTGAGT
ACCCAGTCCGTGCTGTCAAAAGATCCGAACGAAAAACGTGACCACATGGTCCTGCTGGAATTTGTGACG
GCTGCGGGTATCACCCACGGCATGGACGAACTGTATAAAGGTGGAGGTGGCAGTATGGAGAAAAAAAT
CACTGGATATACCACCGTTGATATATCCCAATGGCATCGTAAAGAACATTTTGAGGCATTTCAGTCAGTT
GCTCAATGTACCTATAACCAGACCGTTCAGCTGGATATTACGGCCTTTTTAAAGACCGTAAAGAAAAATA
AGCACAAGTTTTATCCGGCCTTTATTCACATTCTTGCCCGCCTGATGAATGCTCATCCGGAATTCCGTATG
GCAATGAAAGACGGTGAGCTGGTGATATGGGATAGTGTTCACCCTTGTTACACCGTTTTCCATGAGCAA
ACTGAAACGTTTTCATCGCTCTGGAGTGAATACCACGACGATTTCCGGCAGTTTCTACACATATATTCGC
AAGATGTGGCGTGTTACGGTGAAAACCTGGCCTATTTCCCTAAAGGGTTTATTGAGAATATGTTTTTCGT
CTCAGCCAATCCCTGGGTGAGTTTCACCAGTTTTGATTTAAACGTGGCCAATATGGACAACTTCTTCGCCC
CCGTTTTCACCATGGGCAAATATTATACGCAAGGCGACAAGGTGCTGATGCCGCTGGCGATTCAGGTTC
ATCATGCCGTTTGTGATGGCTTCCATGTCGGCAGAATGCTTAATGAATTACAACAGTACTGCGATGAGTG
GCAGGGCGGGGCGTGATAAAGCTGGATCCGGCGCCGCTCATTCGCTAATCGCCACGACGCGTAGTCGA
CCCGCCAATTCCCATGGGGCGCGCCTGCTGCCACCGCTGAGCAATAACTAGCATAACCCCTTGGGGCCT
CTAAACGGGTCTTGAGGGGTTTTTTGCCAGGCATCAAATAAAACGAAAGGCTCAGTCGGAAGACTGGG
CCTTTCGTTTTATCTGTTGTTTGTCGGTGAACGCTCTCCTGAGTAGGACAAATCCGCCGGGAGCGGATTT
GAACGTTGTGAAGCAACGGCCCGGAGGGTGGCGGGCAGGACGCCCGCCATAAACTGCCAGGCATCAA
ACTAAGCAGAAGGCCATCCTGACGGATGGCCTTTTTGCGTTTCAGATCTACCGGTAAACCAGCAATAGA
CATAAGCGGCTATTTAACGACCCTGCCCTGAACCGACGACAAGCTGACGACCGGGTCTCCGCAAGTGGC
ACTTTTCGGGGAAATGTGCGCGGAACCCCTATTTGTTTATTTTTCTAAATACATTCAAATATGTATCCGCT
CATGAATTAATTCCTCTTCAAATGTAGCACCTGAAGTCAGCCCCATACGATATAAGTTGTTAATAACTTCG
TATAGCATACATTATACGAAGTTATCTAGTGCTTGGATTCTCACCAATAAAAAACGCCCGGCGGCAACCG
AGCGTTCTGAACAAATCCAGATGGAGTTCTGAGGTCATTACTGGATCTATCAACAGGAGTCCAAGCGAG
CTCTCGAACCCCAGAGTCCCGCTCAGAAGAACTCGTCAAGAAGGCGATAGAAGGCGATGCGCTGCGAA
TCGGGAGCGGCGATACCGTAAAGCACGAGGAAGCGGTCAGCCCATTCGCCGCCAAGCTCTTCAGCAAT
ATCACGGGTAGCCAACGCTATGTCCTGATAGCGGTCCGCCACACCCAGCCGGCCACAGTCGATGAATCC

AGAAAAGCGGCCATTTTCCACCATGATATTCGGCAAGCAGGCATCGCCATGGGTCACGACGAGATCCTC
GCCGTCGGGCATGCGCGCCTTGAGCCTGGCGAACAGTTCGGCTGGCGCGAGCCCCTGATGCTCTTCGTC
CAGATCATCCTGATCGACAAGACCGGCTTCCATCCGAGTACGTGCTCGCTCGATGCGATGTTTCGCTTGG
TGGTCGAATGGGCAGGTAGCCGGATCAAGCGTATGCAGCCGCCGCATTGCATCAGCCATGATGGATAC
TTTCTCGGCAGGAGCAAGGTGAGATGACAGGAGATCCTGCCCCGGCACTTCGCCCAATAGCAGCCAGTC
CCTTCCCGCTTCAGTGACAACGTCGAGCACAGCTGCGCAAGGAACGCCCGTCGTGGCCAGCCACGATAG
CCGCGCTGCCTCGTCCTGCAGTTCATTCAGGGCACCGGACAGGTCGGTCTTGACAAAAAGAACCGGGCG
CCCCTGCGCTGACAGCCGGAACACGGCGGCATCAGAGCAGCCGATTGTCTGTTGTGCCCAGTCATAGCC
GAATAGCCTCTCCACCCAAGCGGCCGGAGAACCTGCGTGCAATCCATCTTGTTCAATCATGCGAAACGA
TCCTCATCCTGTCTCTTGATCAGATCTTGATCCCCTGCGCCATCAGATCCTTGGCGGCAAGAAAGCCATCC
AGTTTAATAACTTCGTATAGCATACATTATACGAAGTTATCTTTGCAGGGCTTCCCAACCTTACCAGAGG
AC

Inducible expression: *araC*-pBad-B0034-*gfp*-G4S-*bla*-lox-*kan*-lox cassette

AraC gene
pBad promoter
B0034 ribosome binding site
*gfp* gene
Gycine(4)-Serine linker
*cat* gene
lox recombination sites
kanamycin resistance gene

GCTCCTAGGTCTGATTCGTTACCAATTATGACAACTTGACGGCTACATCATTCACTTTTTCTTCACAACCG
GCACGGAACTCGCTCGGGCTGGCCCCGGTGCATTTTTTAAATACCCGCGAGAAATAGAGTTGATCGTCA
AAACCAACATTGCGACCGACGGTGGCGATAGGCATCCGGGTGGTGCTCAAAAGCAGCTTCGCCTGGCT
GATACGTTGGTCCTCGCGCCAGCTTAAGACGCTAATCCCTAACTGCTGGCGGAAAAGATGTGACAGACG
CGACGGCGACAAGCAAACATGCTGTGCGACGCTGGCGATATCAAAATTGCTGTCTGCCAGGTGATCGCT
GATGTACTGACAAGCCTCGCGTACCCGATTATCCATCGGTGGATGGAGCGACTCGTTAATCGCTTCCATG
CGCCGCAGTAACAATTGCTCAAGCAGATTTATCGCCAGCAGCTCCGAATAGCGCCCTTCCCCTTGCCCGG
CGTTAATGATTTGCCCAAACAGGTCGCTGAAATGCGGCTGGTGCGCTTCATCCGGGCGAAAGAACCCCG
TATTGGCAAATATTGACGGCCAGTTAAGCCATTCATGCCAGTAGGCGCGCGGACGAAAGTAAACCCACT
GGTGATACCATTCGCGAGCCTCCGGATGACGACCGTAGTGATGAATCTCTCCTGGCGGGAACAGCAAAA
TATCACCCGGTCGGCAAACAAATTCTCGTCCCTGATTTTTCACCACCCCCTGACCGCGAATGGTGAGATT
GAGAATATAACCTTTCATTCCCAGCGGTCGGTCGATAAAAAAATCGAGATAACCGTTGGCCTCAATCGG
CGTTAAACCCGCCACCAGATGGGCATTAAACGAGTATCCCGGCAGCAGGGGATCATTTTGCGCTTCAGC
CATACTTTTCATACTCCCGCCATTCAGAGGAAGAAACCAATTGTCCATATTGCATCAGACATTGCCGTCAC
TGCGTCTTTTACTGGCTCTTCTCGCTAACCAAACCGGTAACCCCGCTTATTAAAAGCATTCTGTAACAAAG
CGGGACCAAAGCCATGACAAAAACGCGTAACAAAAGTGTCTATAATCACGGCAGAAAAGTCCACATTG
ATTATTTGCACGGCGTCACACTTTGCTATGCCATAGCATTTTTATCCATAAGATTAGCGGTTCCTACCTGA
CGCTTTTTATCGCAACTCTCTACTGTTTCTCCGGGTCCCTATCAGTGATAGAGAGAGCTCGTTGAGAAAG
AGGAGAAATACTAGATGCGTAAAGGCGAAGAACTGTTTACCGGTGTGGTTCCGATTCTGGTGGAACTG
GACGGCGATGTTAATGGTCATAAATTCAGTGTTCGCGGCGAAGGTGAAGGCGATGCGACGAACGGCAA
ACTGACCCTGAAATTTATCTGCACCACGGGTAAACTGCCGGTCCCGTGGCCGACGCTGGTGACCACGCT
GACCTATGGCGTTCAATGTTTTGCGCGTTACCCGGATCACATGAAACAGCACGACTTTTTCAAATCGGCC
ATGCCGGAAGGCTATGTGCAGGAACGTACGATTAGCTTTAAAGACGATGGTACGTATAAAACCCGCGC
GGAAGTGAAATTCGAAGGCGATACCCTGGTTAACCGTATCGAACTGAAAGGTATCGATTTCAAAGAAGA
CGGCAATATTCTGGGTCATAAACTGGAATATAACTTCAATTCCCACAACGTGTACATCACCGCGGATAAA
CAGAAAAACGGCATTAAAGCCAATTTCAAAATCCGCCATAATGTGGAAGATGGTAGCGTTCAGCTGGCC
GACCACTATCAGCAAAACACGCCGATTGGTGATGGCCCGGTCCTGCTGCCGGACAATCACTACCTGAGT
ACCCAGTCCGTGCTGTCAAAAGATCCGAACGAAAAACGTGACCACATGGTCCTGCTGGAATTTGTGACG
GCTGCGGGTATCACCCACGGCATGGACGAACTGTATAAAGGTGGAGGTGGCAGTATGAGTATTCAACATTTCC
GTGTCGCCCTTATTCCCTTTTTTGCGGCATTTTGCCTTCCTGTTTTTGCTCACCCAGAAACGCTGGTGAAAGTAA
AAGATGCTGAAGATCAGTTGGGTGCACGAGTGGGTTACATCGAACTGGATCTCAACAGCGGTAAGATCCTTGA
GAGTTTTCGCCCCGAAGAACGTTTTCCAATGATGAGCACTTTTAAAGTTCTGCTATGTGGCGCGGTATTATCCC
GTATTGACGCCGGGCAAGAGCAACTCGGTCGCCGCATACACTATTCTCAGAATGACTTGGTTGAGTACTCACC
AGTCACAGAAAAGCATCTTACGGATGGCATGACAGTAAGAGAATTATGCAGTGCTGCCATAACCATGAGTGA
TAACACTGCGGCCAACTTACTTCTGACAACGATCGGAGGACCGAAGGAGCTAACCGCTTTTTTGCACAACATG
GGGGATCATGTAACTCGCCTTGATCGTTGGGAACCGGAGCTGAATGAAGCCATACCAAACGACGAGCGTGAC
ACCACGATGCCTGTAGCAATGGCAACAACGTTGCGCAAACTATTAACTGGCGAACTACTTACTCTAGCTTCCC
GGCAACAATTAATAGACTGGATGGAGGCGGATAAAGTTGCAGGACCACTTCTGCGCTCGGCCCTTCCGGCTGG
CTGGTTTATTGCTGATAAATCTGGAGCCGGTGAGCGTGGGTCTCGCGGTATCATTGCAGCACTGGGGCCAGAT
GGTAAGCCCTCCCGTATCGTAGTTATCTACACGACGGGGAGTCAGGCAACTATGGATGAACGAAATAGACAG
ATCGCTGAGATAGGTGCCTCACTGATTAAGCATTGGTGATAAAGCTGGATCCGGCGCCGCTCATTCGCTAATC
GCCACGACGCGTAGTCGACCCGCCAATTCCCATGGGGCGCGCCTGCTGCCACCGCTGAGCAATAACTAGCATA

ACCCCTTGGGGCCTCTAAACGGGTCTTGAGGGGTTTTTTGCCAGGCATCAAATAAAACGAAAGGCTCAGTCGG
AAGACTGGGCCTTTCGTTTTATCTGTTGTTTGTCGGTGAACGCTCTCCTGAGTAGGACAAATCCGCCGGGAGCG
GATTTGAACGTTGTGAAGCAACGGCCCGGAGGGTGGCGGGCAGGACGCCCGCCATAAACTGCCAGGCATCAA
ACTAAGCAGAAGGCCATCCTGACGGATGGCCTTTTTGCGTTTCAGATCTACCGGTAAACCAGCAATAGACATA
AGCGGCTATTTAACGACCCTGCCCTGAACCGACGACAAGCTGACGACCGGGTCTCCGCAAGTGGCACTTTTCG
GGGAAATGTGCGCGGAACCCCTATTTGTTTATTTTTCTAAATACATTCAAATATGTATCCGCTCATGAATTAAT
TCCTCTTCAAATGTAGCACCTGAAGTCAGCCCCATACGATATAAGTTGTTAATAACTTCGTATAGCATACATTA
TACGAAGTTATCTAGTGCTTGGATTCTCACCAATAAAAAACGCCCGGCGGCAACCGAGCGTTCTGAACAAATC
CAGATGGAGTTCTGAGGTCATTACTGGATCTATCAACAGGAGTCCAAGCGAGCTCTCGAACCCCAGAGTCCCG
CTCAGAAGAACTCGTCAAGAAGGCGATAGAAGGCGATGCGCTGCGAATCGGGAGCGGCGATACCGTAAAGC
ACGAGGAAGCGGTCAGCCCATTCGCCGCCAAGCTCTTCAGCAATATCACGGGTAGCCAACGCTATGTCCTGAT
AGCGGTCCGCCACACCCAGCCGGCCACAGTCGATGAATCCAGAAAAGCGGCCATTTTCCACCATGATATTCGG
CAAGCAGGCATCGCCATGGGTCACGACGAGATCCTCGCCGTCGGGCATGCGCGCCTTGAGCCTGGCGAACAG
TTCGGCTGGCGCGAGCCCCTGATGCTCTTCGTCCAGATCATCCTGATCGACAAGACCGGCTTCCATCCGAGTA
CGTGCTCGCTCGATGCGATGTTTCGCTTGGTGGTCGAATGGGCAGGTAGCCGGATCAAGCGTATGCAGCCGCC
GCATTGCATCAGCCATGATGGATACTTTCTCGGCAGGAGCAAGGTGAGATGACAGGAGATCCTGCCCCGGCA
CTTCGCCCAATAGCAGCCAGTCCCTTCCCGCTTCAGTGACAACGTCGAGCACAGCTGCGCAAGGAACGCCCGT
CGTGGCCAGCCACGATAGCCGCGCTGCCTCGTCCTGCAGTTCATTCAGGGCACCGGACAGGTCGGTCTTGACA
AAAAGAACCGGGCGCCCCTGCGCTGACAGCCGGAACACGGCGGCATCAGAGCAGCCGATTGTCTGTTGTGCC
CAGTCATAGCCGAATAGCCTCTCCACCCAAGCGGCCGGAGAACCTGCGTGCAATCCATCTTGTTCAATCATGC
GAAACGATCCTCATCCTGTCTCTTGATCAGATCTTGATCCCCTGCGCCATCAGATCCTTGGCGGCAAGAAAGC
CATCCAGTTTAATAACTTCGTATAGCATACATTATACGAAGTTATCTTTGCAGGGCTTCCCAACCTTACCAGAG
GAC

Constitutive expression: J23100-B0034-*gfp*-G4S-*cat*-lox-*kan*-lox cassette

Legend

J23100 promoter
B0034 ribosome binding site
*gfp* gene
Gycine(4)-Serine linker
*cat* gene
lox recombination sites
kanamycin resistance gene

ATACGATATAAGTTGTAATTCTCATGTTAGTCATGCCCCGCGCCCACCGGAAGGAGCTGACTGGGTTGCT
CCTAGGTCTGATTCGTTACCAATTGACGGCTAGCTCAGTCCTAGGTACAGTGCTAGCTTTCTCCGGGTCC
CTATCAGTGATAGAGAGAGCTCGTTGAGAAAGAGGAGAAATACTAGATGCGTAAAGGCGAAGAACTGT
TTACCGGTGTGGTTCCGATTCTGGTGGAACTGGACGGCGATGTTAATGGTCATAAATTCAGTGTTCGCG
GCGAAGGTGAAGGCGATGCGACGAACGGCAAACTGACCCTGAAATTTATCTGCACCACGGGTAAACTG
CCGGTCCCGTGGCCGACGCTGGTGACCACGCTGACCTATGGCGTTCAATGTTTTGCGCGTTACCCGGAT
CACATGAAACAGCACGACTTTTTCAAATCGGCCATGCCGGAAGGCTATGTGCAGGAACGTACGATTAGC
TTTAAAGACGATGGTACGTATAAAACCCGCGCGGAAGTGAAATTCGAAGGCGATACCCTGGTTAACCGT
ATCGAACTGAAAGGTATCGATTTCAAAGAAGACGGCAATATTCTGGGTCATAAACTGGAATATAACTTC
AATTCCCACAACGTGTACATCACCGCGGATAAACAGAAAAACGGCATTAAAGCCAATTTCAAAATCCGC
CATAATGTGGAAGATGGTAGCGTTCAGCTGGCCGACCACTATCAGCAAAACACGCCGATTGGTGATGGC
CCGGTCCTGCTGCCGGACAATCACTACCTGAGTACCCAGTCCGTGCTGTCAAAAGATCCGAACGAAAAA
CGTGACCACATGGTCCTGCTGGAATTTGTGACGGCTGCGGGTATCACCCACGGCATGGACGAACTGTAT
AAAGGTGGAGGTGGCAGTATGGAGAAAAAAATCACTGGATATACCACCGTTGATATATCCCAATGGCAT
CGTAAAGAACATTTTGAGGCATTTCAGTCAGTTGCTCAATGTACCTATAACCAGACCGTTCAGCTGGATA
TTACGGCCTTTTTAAAGACCGTAAAGAAAAATAAGCACAAGTTTTATCCGGCCTTTATTCACATTCTTGCC
CGCCTGATGAATGCTCATCCGGAATTCCGTATGGCAATGAAAGACGGTGAGCTGGTGATATGGGATAGT
GTTCACCCTTGTTACACCGTTTTCCATGAGCAAACTGAAACGTTTTCATCGCTCTGGAGTGAATACCACGA
CGATTTCCGGCAGTTTCTACACATATATTCGCAAGATGTGGCGTGTTACGGTGAAAACCTGGCCTATTTC
CCTAAAGGGTTTATTGAGAATATGTTTTTCGTCTCAGCCAATCCCTGGGTGAGTTTCACCAGTTTTGATTT
AAACGTGGCCAATATGGACAACTTCTTCGCCCCCGTTTTCACCATGGGCAAATATTATACGCAAGGCGAC
AAGGTGCTGATGCCGCTGGCGATTttcTTCATCATGCCGTTTGTGATGGCTTCCATGTCGGCAGAATGCTT
AATGAATTACAACAGTACTGCGATGAGTGGCAGGGCGGGGCGTGATAAAGCTGGATCCGGCGCCGCTC
ATTCGCTAATCGCCACGACGCGTAGTCGACCCGCCAATTCCCATGGGGCGCGCCTGCTGCCACCGCTGA
GCAATAACTAGCATAACCCCTTGGGGCCTCTAAACGGGTCTTGAGGGGTTTTTTGCCAGGCATCAAATA
AAACGAAAGGCTCAGTCGGAAGACTGGGCCTTTCGTTTTATCTGTTGTTTGTCGGTGAACGCTCTCCTGA
GTAGGACAAATCCGCCGGGAGCGGATTTGAACGTTGTGAAGCAACGGCCCGGAGGGTGGCGGGCAGG
ACGCCCGCCATAAACTGCCAGGCATCAAACTAAGCAGAAGGCCATCCTGACGGATGGCCTTTTTGCGTT
TCAGATCTACCGGTAAACCAGCAATAGACATAAGCGGCTATTTAACGACCCTGCCCTGAACCGACGACA
AGCTGACGACCGGGTCTCCGCAAGTGGCACTTTTCGGGGAAATGTGCGCGGAACCCCTATTTGTTTATTT
TTCTAAATACATTCAAATATGTATCCGCTCATGAATTAATTCCTCTTCAAATGTAGCACCTGAAGTCAGCC

CCATACGATATAAGTTGTTAATAACTTCGTATAGCATACATTATACGAAGTTATCTAGTGCTTGGATTCTC
ACCAATAAAAAACGCCCGGCGGCAACCGAGCGTTCTGAACAAATCCAGATGGAGTTCTGAGGTCATTAC
TGGATCTATCAACAGGAGTCCAAGCGAGCTCTCGAACCCCAGAGTCCCGCTCAGAAGAACTCGTCAAGA
AGGCGATAGAAGGCGATGCGCTGCGAATCGGGAGCGGCGATACCGTAAAGCACGAGGAAGCGGTCAG
CCCATTCGCCGCCAAGCTCTTCAGCAATATCACGGGTAGCCAACGCTATGTCCTGATAGCGGTCCGCCAC
ACCCAGCCGGCCACAGTCGATGAATCCAGAAAAGCGGCCATTTTCCACCATGATATTCGGCAAGCAGGC
ATCGCCATGGGTCACGACGAGATCCTCGCCGTCGGGCATGCGCGCCTTGAGCCTGGCGAACAGTTCGGC
TGGCGCGAGCCCCTGATGCTCTTCGTCCAGATCATCCTGATCGACAAGACCGGCTTCCATCCGAGTACGT
GCTCGCTCGATGCGATGTTTCGCTTGGTGGTCGAATGGGCAGGTAGCCGGATCAAGCGTATGCAGCCGC
CGCATTGCATCAGCCATGATGGATACTTTCTCGGCAGGAGCAAGGTGAGATGACAGGAGATCCTGCCCC
GGCACTTCGCCCAATAGCAGCCAGTCCCTTCCCGCTTCAGTGACAACGTCGAGCACAGCTGCGCAAGGA
ACGCCCGTCGTGGCCAGCCACGATAGCCGCGCTGCCTCGTCCTGCAGTTCATTCAGGGCACCGGACAGG
TCGGTCTTGACAAAAAGAACCGGGCGCCCCTGCGCTGACAGCCGGAACACGGCGGCATCAGAGCAGCC
GATTGTCTGTTGTGCCCAGTCATAGCCGAATAGCCTCTCCACCCAAGCGGCCGGAGAACCTGCGTGCAA
TCCATCTTGTTCAATCATGCGAAACGATCCTCATCCTGTCTCTTGATCAGATCTTGATCCCCTGCGCCATCA
GATCCTTGGCGGCAAGAAAGCCATCCAGTTTAATAACTTCGTATAGCATACATTATACGAAGTTATCTTT
GCAGGGCTTCCCAACCTTACCAGAGGAC

*cat* gene mutants

*cat*-$T_{172}A$

ATGGAGAAAAAAATCACTGGATATACCACCGTTGATATATCCCAATGGCATCGTAAAGAACATTTTGAG
GCATTTCAGTCAGTTGCTCAATGTACCTATAACCAGACCGTTCAGCTGGATATTACGGCCTTTTTAAAGAC
CGTAAAGAAAAATAAGCACAAGTTTTATCCGGCCTTTATTCACATTCTTGCCCGCCTGATGAATGCTCATC
CGGAATTCCGTATGGCAATGAAAGACGGTGAGCTGGTGATATGGGATAGTGTTCACCCTTGTTACACCG
TTTTCCATGAGCAAACTGAAACGTTTTCATCGCTCTGGAGTGAATACCACGACGATTTCCGGCAGTTTCT
ACACATATATTCGCAAGATGTGGCGTGTTACGGTGAAAACCTGGCCTATTTCCCTAAAGGGTTTATTGAG
AATATGTTTTTCGTCTCAGCCAATCCCTGGGTGAGTTTCACCAGTTTTGATTTAAACGTGGCCAATATGGA
CAACTTCTTCGCCCCCGTTTTCGCGATGGGCAAATATTATACGCAAGGCGACAAGGTGCTGATGCCGCTG
GCGATTCAGGTTCATCATGCCGTTTGTGATGGCTTCCATGTCGGCAGAATGCTTAATGAATTACAACAGT
ACTGCGATGAGTGGCAGGGCGGGGCG

*cat*-$H_{193}Q$

ATGGAGAAAAAAATCACTGGATATACCACCGTTGATATATCCCAATGGCATCGTAAAGAACATTTTGAG
GCATTTCAGTCAGTTGCTCAATGTACCTATAACCAGACCGTTCAGCTGGATATTACGGCCTTTTTAAAGAC
CGTAAAGAAAAATAAGCACAAGTTTTATCCGGCCTTTATTCACATTCTTGCCCGCCTGATGAATGCTCATC
CGGAATTCCGTATGGCAATGAAAGACGGTGAGCTGGTGATATGGGATAGTGTTCACCCTTGTTACACCG
TTTTCCATGAGCAAACTGAAACGTTTTCATCGCTCTGGAGTGAATACCACGACGATTTCCGGCAGTTTCT
ACACATATATTCGCAAGATGTGGCGTGTTACGGTGAAAACCTGGCCTATTTCCCTAAAGGGTTTATTGAG
AATATGTTTTTCGTCTCAGCCAATCCCTGGGTGAGTTTCACCAGTTTTGATTTAAACGTGGCCAATATGGA
CAACTTCTTCGCCCCCGTTTTCACCATGGGCAAATATTATACGCAAGGCGACAAGGTGCTGATGCCGCTG
GCGATTCAGGTTCATCAGGCCGTTTGTGATGGCTTCCATGTCGGCAGAATGCTTAATGAATTACAACAGT
ACTGCGATGAGTGGCAGGGCGGGGCG

*bla* gene mutant

*bla*-$L_{74}N$

ATGAGTATTCAACATTTCCGTGTCGCCCTTATTCCCTTTTTTGCGGCATTTTGCCTTCCTGTTTTTGCTCACCCAG
AAACGCTGGTGAAAGTAAAAGATGCTGAAGATCAGTTGGGTGCACGAGTGGGTTACATCGAACTGGATCTCA
ACAGCGGTAAGATCCTTGAGAGTTTTCGCCCCGAAGAACGTTTTCCAATGATGAGCACTTTTAAAGTTCTGAA
CTGTGGCGCGGTATTATCCCGTATTGACGCCGGGCAAGAGCAACTCGGTCGCCGCATACACTATTCTCAGAAT
GACTTGGTTGAGTACTCACCAGTCACAGAAAAGCATCTTACGGATGGCATGACAGTAAGAGAATTATGCAGTG
CTGCCATAACCATGAGTGATAACACTGCGGCCAACTTACTTCTGACAACGATCGGAGGACCGAAGGAGCTAA
CCGCTTTTTTGCACAACATGGGGGATCATGTAACTCGCCTTGATCGTTGGGAACCGGAGCTGAATGAAGCCAT
ACCAAACGACGAGCGTGACACCACGATGCCTGTAGCAATGGCAACAACGTTGCGCAAACTATTAACTGGCGA
ACTACTTACTCTAGCTTCCCGGCAACAATTAATAGACTGGATGGAGGCGGATAAAGTTGCAGGACCACTTCTG
CGCTCGGCCCTTCCGGCTGGCTGGTTTATTGCTGATAAATCTGGAGCCGGTGAGCGTGGGTCTCGCGGTATCAT
TGCAGCACTGGGGCCAGATGGTAAGCCCTCCCGTATCGTAGTTATCTACACGACGGGGAGTCAGGCAACTATG
GATGAACGAAATAGACAGATCGCTGAGATAGGTGCCTCACTGATTAAGCATTGGTAA

### 4 Supplementary videos

Supplementary Video 1: Transient upregulation of gfp-cat at$f$ mRNA and protein level observed in constitutive emergent gene expression model (see section 2.5). Upper panel shows dynamic histograms of GFP-CAT protein distributions from simulations of the constitutive promoter model for 13.07 hours for cells treated with 2, 4 or 6 $\mu g/mL$ Cm (coloured in blue, red and green respectively). Lower panel shows dynamic histograms of gfp-cat mRNA distributions from simulations of the constitutive promoter model for 13.07 hours for cells treated with 2, 4 or 6 $\mu g/mL$ Cm (coloured in blue, red and green respectively). X-axes are displayed on log10 scale and y-axes are scaled such that the total area of the histograms sum to 1.

Supplementary Video 2: Sustained upregulation of gfp-cat at mRNA and protein level observed in inducible emergent gene expression model (see section 2.6). Upper panel shows dynamic histograms of GFP-CAT protein distributions from simulations of the constitutive promoter model for 11.71 hours for cells treated with 0, 15 or 30 $\mu g/mL$ Cm (coloured in blue, red and green respectively). Lower panel shows dynamic histograms of gfp-cat mRNA distributions from simulations of the constitutive promoter model for 11.71 hours for cells treated with 0, 15 or 30 $\mu g/mL$ Cm (coloured in blue, red and green respectively). X-axes are displayed on log10 scale and y-axes are scaled such that the total area of the histograms sum to 1.
