## Supplementary figures and images for "Ohm’s Law for increasing fitness gene expression with selection pressure"

### Supplementary video 1

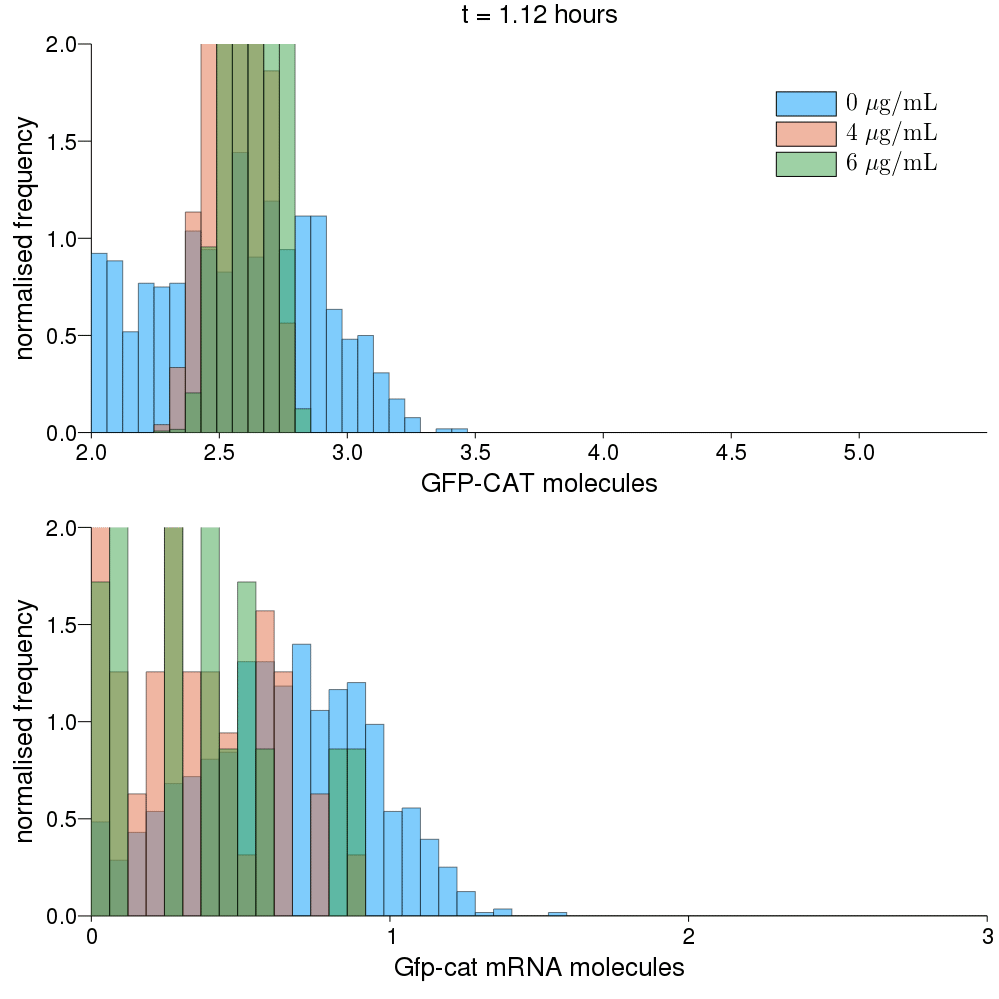

### Supplementary video 2

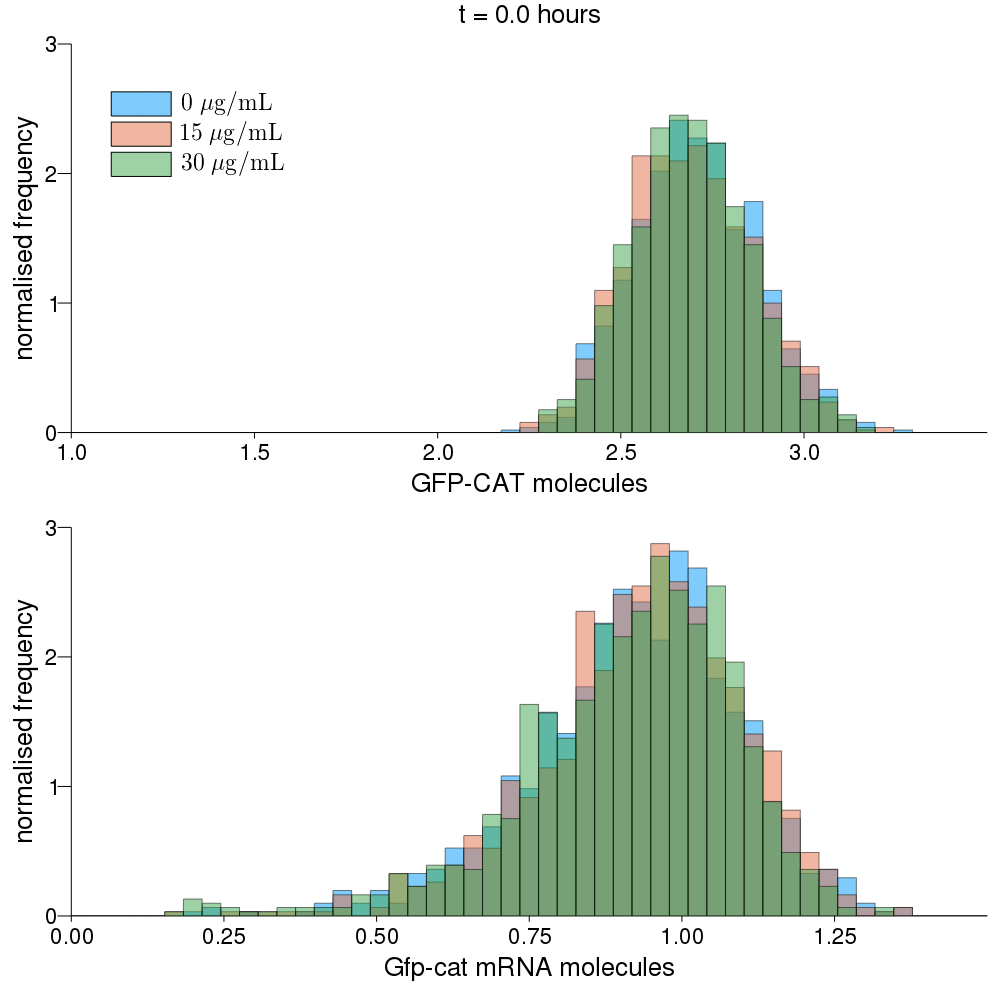
